# Identification and characterization of host-modulating effectors encoded by the Cluster F1 mycobacteriophage NormanBulbieJr

**DOI:** 10.64898/2025.12.12.693979

**Authors:** Bethany M. Wise, Kaylia Edwards, Cole R. Jirsa, Sabrina Abbruzzese, Amina Adebiyi, Sarika Bapat, Tyler Barnhardt, Naomi Bastiampillai, Ella Begovic, Mary G. Berchick, Gilda Bocco, John Bonoris, Elizabeth Boos, Margaret Cassady, Julia Chehab, Gene Cooper, Hunter Coyle, Joseph Davis, Natalie De la Cruz Vargas, Matthew Delach, Carly Dowiak, David Ferraro, Madison Fuller, Bootsie Glasser, Kyle Gordon, Hannah Hoch, Sarah Holleman, Luke Hood, Brett Hurrell, Nicolas Jacobs, Danny Jiang, Steven Kefalos, Salma Maher, Jocelyn Martin, Rohebot Mengesha, Daniel Merenich, Ashwin Nayak, Dasha Nesterova, Joshua Nguyen, Chukwudumebi Okonkwo, Taylor Pompan, Lauren Redwood, Christina Scanlon, Lily Schneider, Aparna Shenai, Colton Siatkowski, Lisa Strausner, Joseph Vizzarri, Jordan Watson, Angela Wilson, Catherine Wynne, Anthony Zygmunt, Michelle Kanther, Kimberly M. Payne, Richard S. Pollenz, C. N. Sunnen, Danielle M. Heller

**Affiliations:** Department of Science Education, Howard Hughes Medical Institute, Chevy Chase, Maryland 20815, USA; Department of Biology, Saint Joseph’s University, Philadelphia, Pennsylvania 19131, USA; Department of Biological Sciences, Dietrich School of Arts and Sciences, University of Pittsburgh, Pittsburgh, Pennsylvania 15260, USA; Department of Bio-Medical Sciences, Philadelphia College of Osteopathic Medicine, Philadelphia, Pennsylvania 19131, USA; Department of Molecular Biosciences, University of South Florida, Tampa, Florida 33620, USA

## Abstract

NormanBulbieJr (NBJ) is a temperate siphovirus isolated on the host *Mycobacterium smegmatis* mc^2^155 that encodes 102 gene products, 60 of which have no known function (NKF). Based on gene content, NBJ is classified as a Cluster F, Subcluster F1 phage and shares 70% of its encoded gene phamilies with Girr, another F1 mycobacteriophage that was recently analyzed in a genome-wide overexpression screen and found to encode 29 diverse gene products capable of inhibiting growth of *M. smegmatis*. Similar functional screens in other mycobacteriophages have uncovered a growing repertoire of diverse, phage-encoded bacterial growth inhibitors, providing prime candidates for dissecting novel bacterial-phage interactions. An arrayed overexpression library encoding all 102 genes was constructed and systematically screened using a plate-based cytotoxicity assay, identifying 29 genes that inhibit mycobacterial growth. Because mycobacteriophage genomes are also known to encode systems involved in phage-phage competition, we conducted additional phage defense assays for a subset of NBJ genes in our library. This analysis confirmed homotypic immunity by the predicted immunity repressor and identified gp45 as a second defense factor that protects against Cluster F phages and promotes NBJ lysogen stability. Additional mechanistic analyses indicate that gp45 functions in a secretion-dependent manner, acting after phage adsorption to reduce productive infection and early phage transcript accumulation. Escape mutant and bacterial two-hybrid analyses implicate Cluster F tape measure proteins as the likely target of gp45, consistent with disruption of an early step in phage genome delivery. Finally, we extended our analysis to explore the essentiality of all identified host-modulating genes in the NBJ life cycle, using CRISPR-enhanced recombineering to generate phage deletion mutants, revealing two host modulators that are critical for lytic growth. Conducted as part of the SEA-GENES undergraduate research consortium, this study adds to a growing functional genomics resource and provides new insights into the complex interactions between phage gene products and the mycobacterial host cell.

## Introduction

The global bacteriophage population is vast and remarkably diverse. Nearly 5,500 genomes of phages infecting actinobacteria have been sequenced to date, largely through the efforts of the SEA-PHAGES undergraduate research consortium, exposing a dynamic genetic landscape shaped by rampant horizontal exchange and mosaicism (Russell and Hatfull 2016; Hatfull 2020; Heller et al. 2024). Collectively, these genomes encode more than 32,000 different gene phamilies, or phams, composed of related sequences with significant amino acid similarity (Cresawn et al. 2011; Gauthier et al. 2022) , yet only a fraction can be assigned a predicted function based on synteny and sequence homology (Hatfull 2020). The remainder represent an enormous reservoir of uncharacterized proteins whose roles in phage infection and modulation of bacterial physiology are largely unexplored.

Recently, functional genomics efforts in multiple phage systems have begun to bridge this gap, accelerating the identification of phage and host genes that are important for phage infection. On the phage side, deletion and knockout studies have shown that many phage genes are dispensable for lytic growth, forming a large and variable accessory gene pool that is likely beneficial under specific conditions (Dedrick et al. 2013; Chen et al. 2025). More recently, CRISPRi knockdown screens in coliphages have demonstrated that many genes of unknown function (NKF) contribute measurably to phage fitness (Adler et al. 2025). Systematic overexpression screens in several phages that infect *Mycobacterium smegmatis* have been employed to identify those genes that perturb mycobacterial physiology, uncovering numerous phage-encoded proteins, largely NKF, that influence bacterial growth (Ko and Hatfull 2020; Heller et al. 2022; Amaya et al. 2023; Freeman et al. 2024; Pollenz et al. 2024; Tafoya et al. 2024; Cagang et al. 2026). Functional genomics studies of mycobacteriophages have also revealed that many temperate phages encode diverse systems that mediate phage-phage competition, including canonical immunity repressors and a variety of other defense modules (Dedrick et al. 2017; Gentile et al. 2019; Montgomery et al. 2019) . On the host side, there has been a recent explosion in the number of systems found to be employed by bacteria to counter phage infection through abortive or immunity-like mechanisms (LeRoux and Laub 2022; Georjon and Bernheim 2023). Notably, many of these systems are encoded within prophages, reinforcing phages as a repository of important host-modulatory functions (Dedrick et al. 2021). Thus, defining the molecular functions of phage gene products and the mechanisms by which they alter bacterial physiology remains an important challenge.

Building on this functional genomics foundation, we carried out a genome-wide analysis of the Cluster F1 mycobacteriophage NormanBulbieJr (NBJ), a temperate siphovirus isolated on *M. smegmatis* mc^2^155 that encodes for 102 gene products (Russell and Hatfull 2016; Gauthier and Hatfull 2023), 60 of which have no known function (Figure 1). Using an arrayed whole-genome overexpression library, we identified NBJ genes that influence bacterial growth and susceptibility to phage and employed CRISPR-enhanced recombineering to assess their roles in the NBJ life cycle. Conducted as part of the SEA-GENES undergraduate research consortium (Heller et al. 2024), this study extends the SEA-GENES functional genomics platform to provide new insights into how mycobacteriophage genes shape cellular processes and to define promising targets for future mechanistic studies.

**Figure 1:**
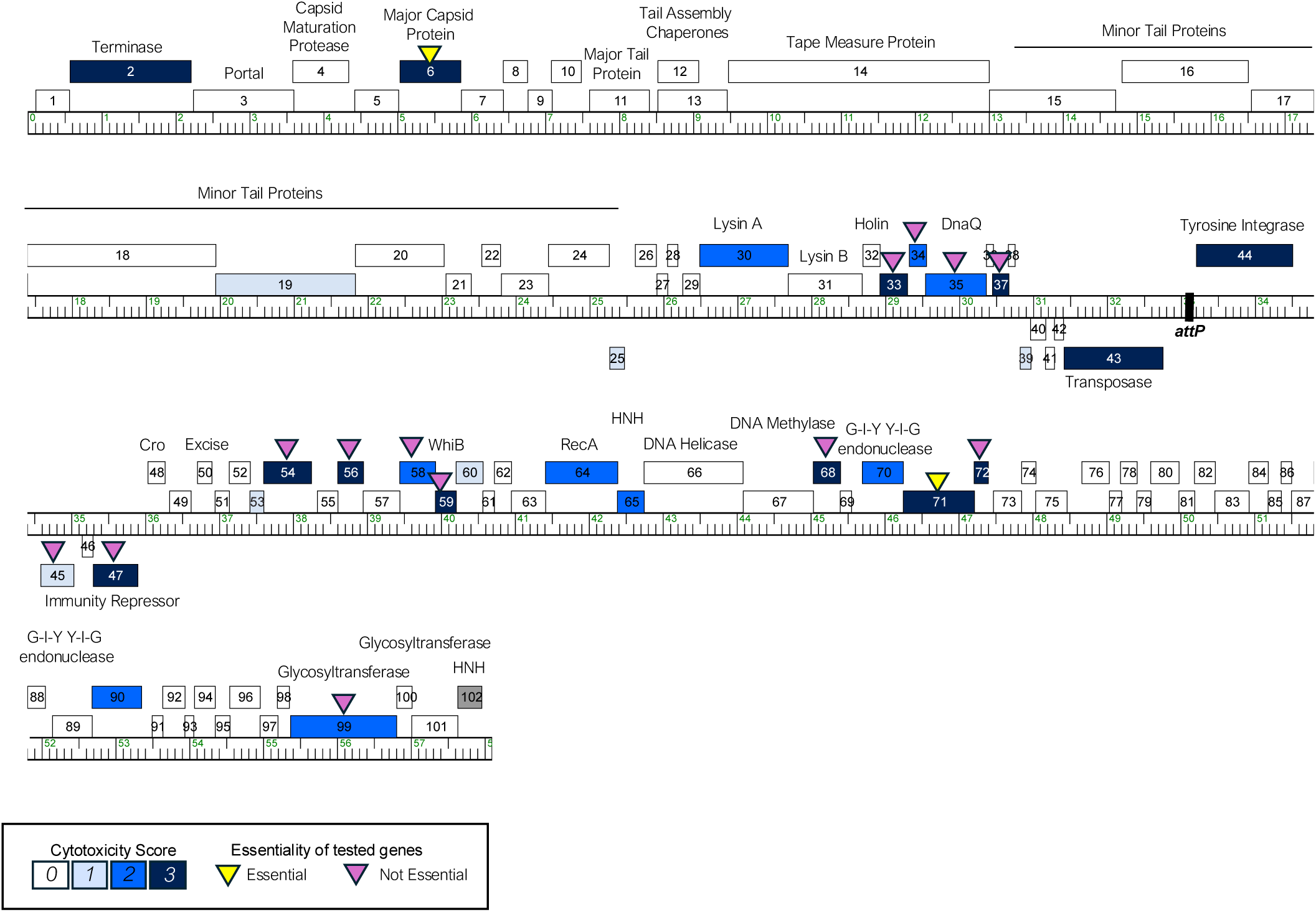
The genome of phage NormanBulbieJr. The NormanBulbieJr (NBJ) genome is shown as a line with kbp markers and genes represented by boxes—those above the line are transcribed rightwards and those below are transcribed leftwards. Numbers inside the box correspond to gene numbers and predicted functions are indicated above or below each gene. Box shading corresponds to cytotoxicity scoring, with white boxes designating genes found to have no effect on *M. smegmatis growth* (cytotoxicity score 0), gray box indicating gene *102* for which pExTra transformants could not be isolated, and blue representing observed toxicity in our assay. The saturation of blue boxes corresponds to the severity of growth inhibition using the following scores: light blue (score 1; reduction in colony size; genes *19, 25, 39, 45, 53*, and *60)*, medium blue (score 2; 1–3 log reduction in viability; genes *30, 34, 35, 58, 64, 65, 70, 90*, and *99*), and dark blue (score 3; >3-log reduction in viability; genes *92, 6, 33, 37, 43, 44, 47, 54, 56, 59, 68, 71*, and *72)*. Purple triangles positioned above the gene boxes indicate which genes were evaluated for essentiality by CRISPY-BRED, with yellow triangles signifying the two genes (*6* and *71*) found to be essential for lytic growth.

## Materials and Methods

### Growth of mycobacteria and mycobacteriophages

*M. smegmatis* mc^2^155 was grown in Middlebrook 7H9 (Difco) broth supplemented with 10% AD (2% w/v Dextrose, 145 mM NaCl, 5% w/v Albumin Fraction V), 0.05% Tween80, and 10 µg/ml cycloheximide (CHX) or on Middlebrook 7H10 (Difco) or 7H11 (Remel) agar supplemented with 10% AD, 10 µg/ml CHX, and kanamycin 10 µg/ml (GoldBio) as needed for selection of pExTra plasmids. To transform *M. smegmatis* mc^2^155, electrocompetent cells were electroporated with ∼100 ng of pExTra plasmid DNA, recovered in 7H9 broth for 2 h at 37 °C with shaking, and transformants selected on 7H11 agar supplemented with kanamycin. After 4 days of incubation at 37 °C, transformants were used directly in plate-based cytotoxicity assays or to inoculate liquid cultures for defense assays. All mycobacteriophages were propagated on *M. smegmatis* mc^2^155 grown at 37 °C in the presence of 1 mM CaCl_2_ and no Tween in Middlebrook media and top agar.

### Construction of plasmids

In this study, phage genes were cloned into the pExTra shuttle vector (Heller et al. 2022) downstream of an anhydrotetracycline-inducible promoter, *pTet* (Ehrt et al. 2005), to control gene expression, and upstream of linked *mcherry* transcriptional reporter (Figure 1A). Genes were PCR-amplified (New England Biolabs Q5 HotStart 2X Master Mix) from a high-titer lysate using a forward primer complementary to the first 15-25 bp of each gene sequence (Integrated DNA Technologies), introducing a uniform ATG start codon, and a reverse primer complementary to the last 15-25 bp of the gene sequence, including a uniform TGA stop codon (Supplemental Table 1). All forward primers contained a uniform, RBS-containing 5**’** 21 bp sequence and all reverse primers contained a separate 5’ 25 bp sequence; these added sequences have identity to the pExTra plasmid flanking the site of insertion. Linearized pExTra plasmid was prepared via PCR (NEB Q5 HotStart 2X Master Mix) of pExTra01 (Heller et al. 2022) using divergent primers pExTra_F and pExTra_R and assembled with each gene insert by isothermal assembly (NEB HiFi 2X Master Mix).

Recombinant plasmids were recovered by transformation of *E. coli NEB5α F’I^Q^* (New England Biolabs) and selection on LB agar supplemented with 50 µg/ml kanamycin. Truncated or mutated plasmid variants were constructed using a similar process, with inserts generated by PCR with mutagenic primers (Supplemental Table 1). The inserted genes for all recovered pExTra plasmids were sequence-verified by Sanger sequencing (Azenta) using sequencing primers pExTra_universalR and pExTra_seqF; longer genes were also sequenced with internal sequencing primers listed in Supplemental Table 1. All plasmid inserts were found to match the published genome sequence (Genbank accession MH399784).

To clone plasmids for assaying in the bacterial two-hybrid (B2H) system, we used the compatible vectors p2Hα and pCIMSLib, which are derivatives of the published pBRα and pAC-λCI plasmids, respectively, from the transcription-based B2H system developed by Dove and Hochschild (Dove et al. 1997; Fu et al. 2004; Heller et al. 2017). Truncated NBJ *45* was amplified from the pExTra-NBJ45ΔN plasmid using pExTra_universal_F and pExTra_universal_R primers. This product was digested with NdeI and BamHI (NEB) and cloned by restriction digest and ligation cloning (T4 ligase, NEB) into plasmid p2Hα, placing the insert under the control of the inducible *lacUV5* promoter as a translational fusion to the N-terminal domain of *E. coli* RpoA. The NBJ gp14 fragment (residues 144-345) was cloned as a translational fusion to the full-length λCI DNA-binding protein in the compatible pCIMSLib plasmid. This sequence was amplified from NBJ lysate using primers oDH458 and oDH459, digested with NdeI and SpeI-HF (NEB) and ligated into the pCIMSLib digested backbone. The R326S mutation was introduced by two-step mutagenic PCR using mutagenic primers oDH462 and oDH463. B2H plasmid sequences were verified by whole-plasmid sequencing (Plasmidsaurus).

### Overexpression screening and phenotype scoring

To assess cytotoxicity, pExTra-transformed colonies were resuspended and serially diluted in 7H9 broth then spotted on 7H11 plates supplemented with 10 µg/ml kanamycin and 0, 10, or 100 ng/ml anhydrotetracycline (aTc; Alfa Aesar). Each strain was tested in triplicate alongside the pExTra02 positive control plasmid, encoding cytotoxic gene Fruitloop *52,* and the pExTra03 negative control plasmid, encoding a non-toxic mutant allele of Fruitloop *52* (I70S) (Ko and Hatfull 2018; Heller et al. 2022). With incubation at 37 °C, spot growth was typically visible after 2 days, with effects on colony size and appearance more apparent after 3-5 days of incubation. Cytotoxic phenotypes were scored by comparing the spot dilution out to which cells grew in the presence versus the absence of aTc inducer and classified as either having no effect (score *0*), being moderately cytotoxic with a 1-3 log reduction in cell viability (score *2*), or being highly cytotoxic, causing complete or near complete (>3-log) inhibition of growth (score *3*). Strains were also evaluated for aTc-dependent size reduction in individual colonies (score *1*) as compared to the Fruitloop *52-I70S* negative control strain on the same aTc plate and the same strain on plates without inducer. Pink colony color from *mcherry* gene expression provided a visual indicator of gene expression through the *pTet* operon.

Reported cytotoxic genes were found to cause growth inhibition in at least two independent experiments with strong agreement between triplicate samples within each experiment. The magnitude of cytotoxicity for some genes was observed to vary slightly between experiments, likely due to minor variations in media or growth conditions; variability was more pronounced in those strains observed to have milder toxic effects, which are potentially at the detection limit of our assay. If differences in magnitude were observed between experiments, genes were scored based on the more conservative result. The inclusion of the Fruitloop *52* control strains on each screening plate aided in the evaluation of relative gene-mediated effects, and results were ultimately scored based on observations on 100 ng/ml aTc.

For plate-based defense assays, pExTra transformants were grown at 37 °C for 48 h in Middlebrook 7H9 broth with 10 µg/ml kanamycin and 1 mM CaCl_2_. Cells were pelleted and resuspended in 7H9 Middlebrook broth lacking Tween80, and 0.5 ml of culture was mixed with 4 ml Middlebrook top agar to generate bacterial lawns on 7H11 agar supplemented with 10 µg/ml kanamycin, 1 mM CaCl_2_, with or without 100 ng/ml aTc.

Lysates of tested phages were serially diluted in phage buffer (10 mM Tris pH 7.5, 10 mM MgSO_4_, 68 mM NaCl, 1 mM CaCl_2_) and spotted on top agar lawns. Plates were incubated at 37 °C for 48 h. Defense was defined as a >10-fold reduction in plaquing efficiency on lawns with aTc-100 compared to lawns without aTc. Positive defense phenotypes were tested in at least two independent experiments.

### Phage recombineering

NBJ gene deletion mutants were generated using the previously described Bacteriophage Recombineering of Electroporated DNA, or BRED, method (Marinelli et al. 2008) with the addition of CRISPR-Cas9 to facilitate efficient selection of recombinant phage (Wetzel et al. 2021). Briefly, to make an unmarked gene deletion in the NBJ genome, substrates were designed and synthesized as dsDNA gblocks (Integrated DNA Technologies) containing the first six codons and last six codons of the gene to be deleted, flanked by 250 bp of genomic sequence upstream and 250 bp of genomic sequence downstream of the gene. Substrates were amplified by PCR using primers designed to anneal at the 5’ and 3’ ends of the synthesized gblock (Supplemental Table 2).

CRISPR-Cas9 selection strains were generated according to the parameters outlined by Rock et al. (Rock et al. 2017) and as described by Wetzel et al. (Wetzel et al. 2021). The gene to be deleted was screened for one of 15 PAM sequences previously shown to allow for >25-fold repression of target sequences by *Streptococcus thermophilus* dCas9 (Rock et al. 2017). Complementary oligonucleotides containing the target sequence adjacent to the selected PAM (sequences shown in Supplementary Table 3) were annealed to form the sgRNA insert containing compatible sticky ends for subsequent ligation (T4 ligase, NEB) into plasmid pIRL53 (Rock et al. 2017) digested with BsmBI-v2 (NEB). Plasmids were recovered by transformation of *E. coli NEB5α F’I^Q^* and selection on LB agar supplemented with kanamycin 50 µg/ml and sequence-verified using primer oBW47.

Verified plasmids were used to transform *M. smegmatis mc^2^155*, and transformants were cultured in Middlebrook 7H9 broth with 10 µg/ml kanamycin and 1 mM CaCl_2_. Cells were pelleted and resuspended in 7H9 Middlebrook broth lacking Tween80 before use in phage assays. sgRNA-mediated selection was first tested by spotting a dilution series of wild-type NormanBulbieJr lysate onto Middlebrook top agar lawns of each selection strain (Supplemental Figure 3) with >100-fold reduction in plaquing indicative of efficient Cas9 targeting.

BRED or CRISPY-BRED were performed as previously described (Marinelli et al. 2008; Wetzel et al. 2021). For the latter, *M. smegmatis mc^2^155/*pJV138 was electroporated with 100-200 ng of NBJ genomic DNA and 100-400 ng of amplified deletion substrate. Cells were recovered in 7H9 broth with no Tween and 1 mM CaCl_2_ at 37 °C for 4 h to allow for lysis and phage release, combined with the appropriate *M. smegmatis* mc^2^155/psgRNA selection strain culture, then split in half and plated with top agar onto 7H11 agar supplemented with kanamycin 10 µg/ml with or without 100 ng/ml aTc (Supplemental Figure 4).

Well-isolated plaques were picked from the plate containing aTc into 100 µL phage buffer. One microliter of each plaque suspension was then used as template for PCR-screening with primers flanking the intended deletion site (NEB Q5 2X Master Mix) (Supplemental Figure 5). Plaques that yielded a band of the expected size were subjected to an additional round of plaque purification before a high titer lysate of mutant phage was generated. For genes *6* and *71*, CRISPY-BRED was independently repeated to confirm the inability to isolate any viable plaques on plates with aTc. Deletion mutants were sequenced by short-read Illumina sequencing using the NextSeq1000 sequencer (P1 reagents) following library preparation using NEBNext Ultra II FS Kit; genomes were assembled as described previously (Jirsa et al. 2026).

### Evaluation of phage gene deletion mutant phenotypes

Plaque size was measured for wild-type and mutant NBJ phages, with each lysate plated with *M. smegmatis* mc^2^155 in top agar on 7H10 agar with 1 mM CaCl_2_ to generate ∼100 well-isolated plaques per plate. After incubation at 37 °C for 48 h, plaque diameter was measured for ∼ 50 plaques using Fiji (Schindelin et al. 2012) and ObjectJ (https://sils.fnwi.uva.nl/bcb/objectj/index.html). Plaque sizes were measured in at least two independent experiments.

For one-step growth curves, wild-type or mutant NBJ was used to infect fresh, log-phase cultures (O.D.600 ∼0.2) of *M. smegmatis mc^2^155* at a multiplicity of infection (MOI) of 0.01, ∼2 x 10^5^ pfu ml^-1^, in experimental triplicate. After a 30 min infection without shaking at room temperature, cells were centrifuged for 10 min at 6,000 *x g*, supernatant containing any unadsorbed phage was removed, and the cell pellet was resuspended in 7H9 broth with 1 mM CaCl_2_. This resuspension was then diluted 100-fold into 25 ml of the same medium, an aliquot removed for time 0, and then, incubated at 37 °C shaking at 220 rpm.

Plaque-forming units were measured over time, with culture aliquots taken at specific times, spun down to pellet any cells, and the supernatant serially diluted and plated with *M. smegmatis mc^2^155* in plaque assays.

To measure frequency of lysogeny, 7H10 agar plates with 1 mM CaCl_2_ were seeded with ∼10^9^ plaque-forming units (pfu) of wild-type or mutant phage, and a dilution series of fresh, saturated *M. smegmatis mc^2^155* culture plated on top. The same dilution series was plated onto 7H10 agar plates not seeded with phage. Frequency of lysogeny for a given phage was calculated as the number of colony-forming units (cfu) on seeded plates divided by the number of cfu on non-seeded plates. Frequency of lysogeny was measured in three independent experiments, with complete results from one representative experiment reported here.

Isolated colonies on seeded plates were subjected to two additional rounds of streak purification, and lysogeny was confirmed by measuring phage release in culture supernatants after 24 h of growth at 37 °C with shaking. Whole-genome sequencing of lysogens was performed by Plasmidsaurus using Oxford Nanopore Technology with custom analysis and annotation.

### NBJ adsorption, post-adsorption infection, and transcript accumulation analysis

To evaluate the step of infection inhibited by gp45, duplicate cultures of *M. smegmatis* mc^2^155 transformed with pExTra03, pExTra-NBJ45, or pExTra-NBJ45ΔN were first grown to high cell density in Middlebrook 7H9 broth containing 0.05% Tween80, 10 µg/ml kanamycin, and 1 mM CaCl_2_. Cultures were then subcultured into 7H9 broth lacking Tween80 and grown again to high density before being back-diluted a second time into fresh 7H9 lacking Tween80 and grown until they reached an O.D.600 of 0.6. At this point, cultures were induced with 100 ng/ml aTc for 4 h at 37 °C with shaking. Induced cultures were then infected with NBJ at a multiplicity of infection (MOI) of 0.1 and incubated at 37 °C without shaking. To measure adsorption, aliquots were removed at 0, 20, and 45 min post-infection, cells were pelleted by centrifugation, and free phage remaining in the supernatant was quantified by plaque assay on *M. smegmatis* mc^2^155.

To measure productive infection after adsorption, duplicate infected cultures were allowed to adsorb for 45 min, then pelleted to remove unbound phage and washed once in fresh 7H9 broth lacking Tween80. Cell pellets were resuspended in fresh medium either lacking or containing 100 ng/ml aTc and incubated at 37 °C with shaking for 180 min.

Culture supernatants were then collected and phage titers measured by spot assay on lawns of *M. smegmatis* mc^2^155.

For transcript analysis, infected cultures prepared as described above were collected at 0, 30, 60, and 180 min after washing and resuspension in fresh medium containing 100 ng/ml aTc. For each sample, 4 ml of culture was pelleted and resuspended in DNA/RNA Shield (Zymo Research) for stabilization prior to RNA extraction. Total RNA was isolated using the Zymo Quick-RNA Fungal/Bacterial Miniprep Kit according to the manufacturer’s instructions, and RNA concentrations were determined using the Qubit RNA Broad Range Kit (Thermo Scientific). Samples were normalized such that 500 ng of total RNA was used as input for each reverse transcription reaction.

RNA was subjected to combined DNase treatment and reverse transcription using the SuperScript IV First-Strand Synthesis System (Thermo Scientific), with matched no-RT control reactions prepared in parallel for each sample. Endpoint PCR was performed using NEB Q5 2X Master Mix and primers targeting NBJ *54*, an early lytic gene selected based on transcriptomics data from the homologous locus in the Cluster F1 phage Fruitloop (Ko and Hatfull 2018). For each +RT sample, undiluted cDNA as well as 1:5 and 1:25 dilutions were used as template. PCR was performed under non-saturating amplification conditions using 24 cycles. DNA isolated from NBJ lysate and *M. smegmatis* mc^2^155 genomic DNA were used as positive and negative controls, respectively, for the NBJ *54* primer set.

### Escape mutant and bacterial two-hybrid analysis

To isolate mutants capable of overcoming gp45-mediated defense, *M. smegmatis* mc^2^155 harboring pExTra-NBJ45 was grown in Middlebrook 7H9 broth with 10 µg/ml kanamycin. Cultures were infected with phages Girr, NBJ, or Tchen at an MOI of 10 and plated in Middlebrook top agar on 7H11 agar supplemented with 10 µg/ml kanamycin and 100 ng/ml aTc to induce gp45 expression during plaquing. Plates were incubated at 37 °C and monitored for plaque formation. Independent escape plaques were picked and subjected to an additional round of plaquing on the same selective host strain. DNA from purified escape mutants and the original Tchen lysate was prepared for whole-genome short-read Illumina sequencing as described above, and assembled genomes (Jirsa et al. 2026)were compared to the parental Tchen genome sequence to identify variants unique to the escape mutants.

Protein-protein interactions were assayed using a transcription-based bacterial two-hybrid system (Dove et al. 1997; Fu et al. 2004; Heller et al. 2017). For this analysis, the gp45ΔN variant lacking the predicted N-terminal signal peptide was used to assay only the mature exported region of gp45. The p2Hα plasmid encoding NBJ gp45ΔN as a translational fusion to the N-terminal domain of *E. coli* RpoA was co-transformed with the compatible pCIMSLib plasmid encoding the wild-type or R326S NBJ gp14 fragment (residues 144–345) as a translational fusion to full-length λCI. Vector controls used in the B2H assay consisted of the corresponding p2Hα plasmid expressing full-length *rpoA* and the compatible pCIMSLib expressing full-length λCI. Plasmid pairs were co-transformed into the *E. coli* FW102 OL2-62 *lacZ* reporter strain and assayed for interaction by measurement of β-galactosidase activity. β-galactosidase activity was reported in Miller units from three independent replicates, with error bars representing standard deviation.

### NBJ genomic analysis

The NBJ genome map was created using the web-based tool Phamerator (phamerator.org). Reported gene functions are based on those available in the NBJ GenBank record (Accession MH399784). In several cases, these GenBank annotations were supplemented by comparison of functional assignments for other homologous genes belonging to the same phamily on PhagesDB (Russell and Hatfull 2016); these designations were confirmed using HHPRED (PDB_mmCIF70, SCOPe70_2.08, Pfam-A_v35, NCBI_Conserved_Domains(CD)_v3.19) (Gabler et al. 2020), NPS Helix-Turn-Helix predictor (https://npsa-prabi.ibcp.fr/), and DeepTMHMM (Hallgren et al. 2022). Gene content comparison between NBJ and Girr genomes was performed using the gene content comparison tool on PhagesDB (https://phagesdb.org/genecontent/), with phamily designations downloaded from the database on February 19, 2025. Reported protein similarities were determined by multiple sequence alignment with Clustal Omega (Madeira et al. 2022). tRNA analysis was performed using tRNAscan-SE2.0 (Chan et al. 2021). Structural predictions were performed using AlphaFold3 (Abramson et al. 2024) on the Google DeepMind server (https://alphafoldserver.com/). Foldseek searches (Kempen et al. 2024) were performed in TM-align mode against the Big Fantastic Viral Database (BFVD) (Kim et al. 2024), AlphaFold/UniProt50 v6 Database (AFDB50) (Fleming et al. 2025), and Protein Databank (PDB100 20240101) to identify structural homologs with TM-score >0.5. NBJ gp14 was modeled in AlphaFold3 as a homotrimer, with trimer stoichiometry chosen based on the tape measure protein organization reported for the mycobacteriophage Bxb1 tail-tip structure (PDB 9D93) (Freeman et al. 2025).

## Results

### Systematic genome-wide screen evaluating impacts of gene overexpression on host growth

To explore the impacts of NBJ gene expression on the mycobacterial host cell, each gene was inserted into the inducible mycobacterial expression plasmid, pExTra (Heller et al. 2022), generating an arrayed whole-genome library of 102 NBJ genes. In this library, each gene is upstream of a transcriptionally linked *mcherry* fluorescent reporter gene and downstream of the inducible *pTet* promoter, which drives expression of this two-gene operon in an aTc-dependent manner (Figure 2A). The host bacterium *M. smegmatis mc^2^155* was transformed with each plasmid in this library, and the impacts of gene overexpression were systematically evaluated in a semi-quantitative plate-based bacterial growth assay. Colonies of transformed strains were suspended, serially diluted, and spotted in triplicate on Middlebrook 7H11 agar supplemented with increasing concentrations of the inducer anhydrotetracycline (aTc) alongside control strains expressing known cytotoxic gene Fruitloop *52* or a nontoxic mutant allele (Fruitloop *52 I70S)* (Ko and Hatfull 2018). Gene-mediated impacts on host growth were scored using the following system: genes for which no change in growth was observed in the presence of aTc were scored as 0, genes that caused an aTc-dependent reduction in bacterial colony size were scored as 1, genes that caused a 1-3-log reduction in bacterial viability in the presence of aTc were scored as 2, and genes that fully abolished growth in the presence of aTc were scored as 3. Figure 2B provides a representative example of the cytotoxicity assay results for each of these scores.

**Figure 2:**
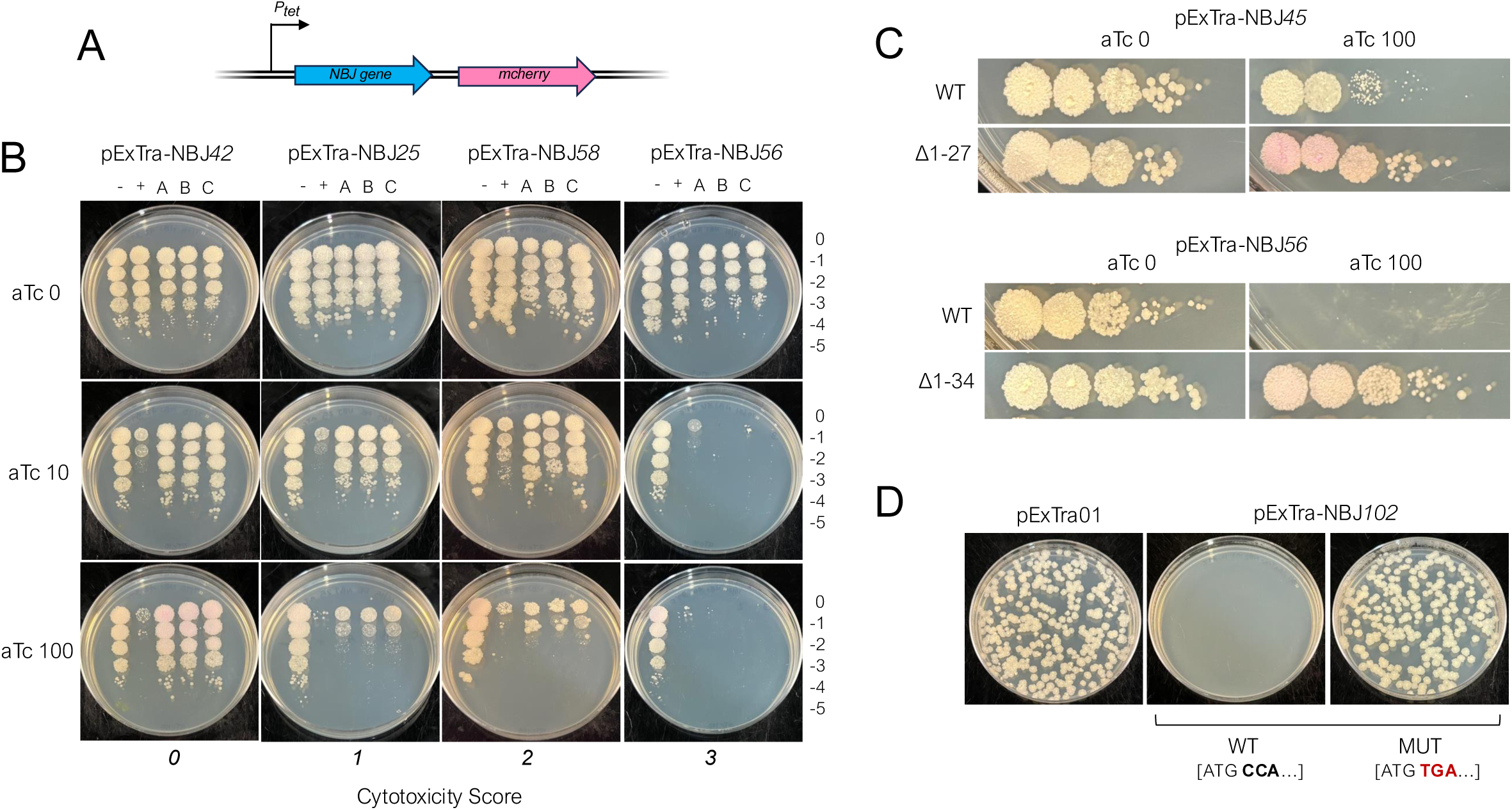
Identification of mycobacterial growth inhibitors encoded by NBJ. A) Recombinant pExTra plasmids constructed in this study encode NBJ gene sequences downstream of the *pTet* promoter and upstream of *mcherry*. The two genes in this pExTra operon are transcriptionally linked, each bearing their own translational signals. B) Results of representative cytotoxicity assays are shown to demonstrate the range of observed growth defects. In each assay, colonies *of M. smegmatis* mc^2^155 transformed with the specified pExTra plasmid were resuspended, serially diluted, and spotted on 7H11 Kan media containing 0, 10, or 100 ng/ml aTc. Triplicate colonies (a, b, c) were tested for each gene alongside a positive control strain (+) transformed with pExTra02 (expressing wild-type Fruitloop *52*) and a negative control strain (−) transformed with pExTra03 (expressing Fruitloop *52 I70S*). C) In a similar cytotoxicity assay, gp45 (top) and gp56 (bottom) variants lacking their predicted N-terminal signal peptides (the first 27 residues of gp45 and 34 residues of gp56) were overproduced from pExTra, showing a loss of cytotoxic phenotype for both. D) *M. smegmatis* mc^2^155 was transformed with 100 ng of pExTra01 or plasmids encoding wild-type or a nonsense allele of NBJ *102* and efficiency of transformant recovery compared by plating serial dilutions on 7H11 Kan.

### Systematic overexpression screen identifies many cytotoxic NBJ genes

Our screen identified 28 genes whose aTc-dependent overexpression caused some observable reduction in host growth (Table 1, Figure 1). Of these 28 genes, 6 were found to cause only a mild reduction in colony size (score 1), 9 caused moderate reduction in viable cell count (score 2), and 13 caused almost complete abolition of growth (score 3) (Table 1). The majority of NBJ genes had no measurable impacts on host growth, and a majority of these displayed some pink coloration consistent with expression through the *pTet* operon (Supplemental Figure 1); however, we cannot exclude the possibility that some genes may be poorly expressed in our system.

**Table 1:**
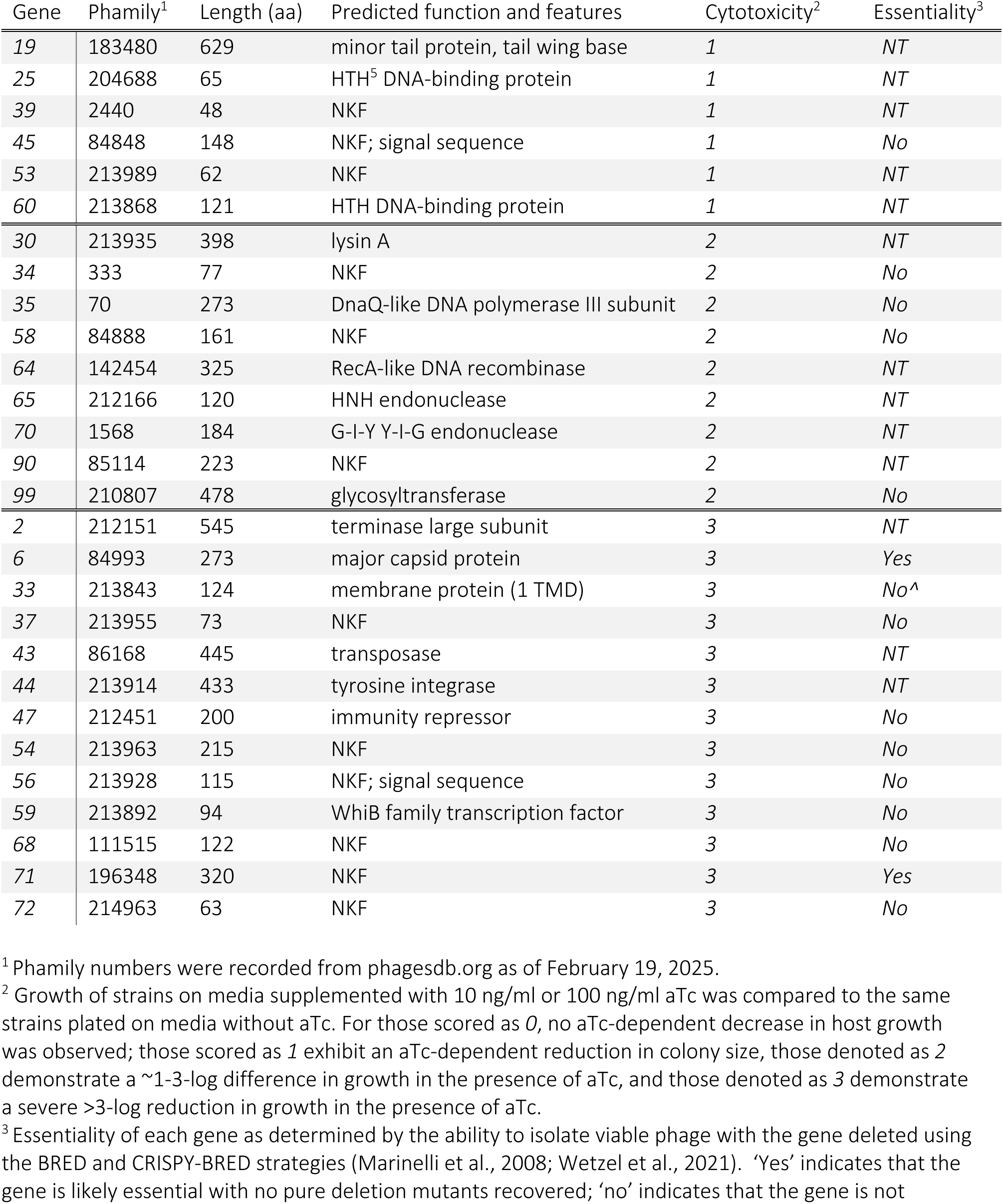
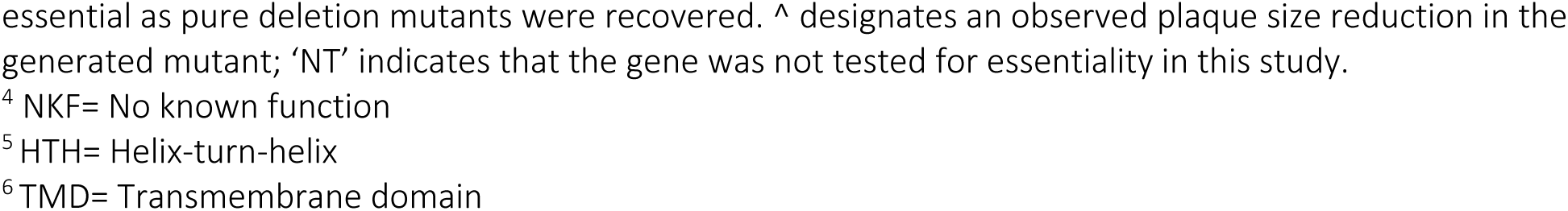
NormanBulbieJr genes observed to inhibit *Mycobacterium smegmatis* growth upon overexpression.

Only 15 of the 28 growth inhibitors identified in our cytotoxicity screen have predicted functions based on synteny and sequence homology (Table 1, Figure 3B). These include three genes involved in virion structure and assembly: the major capsid protein (gp6) and terminase large subunit (gp2), both of which caused severe growth inhibition upon overproduction (score 3), and the NBJ minor tail protein gp19, which caused only mild toxicity (score 1). Comparison to the recent cryo-EM structure of the mycobacteriophage Bxb1 tail tip indicates that NBJ gp19 is likely the tail wing base containing both a predicted glycan-binding FN3 domain and D-ala-D-ala carboxypeptidase domain (Freeman et al. 2025).

**Figure 3:**
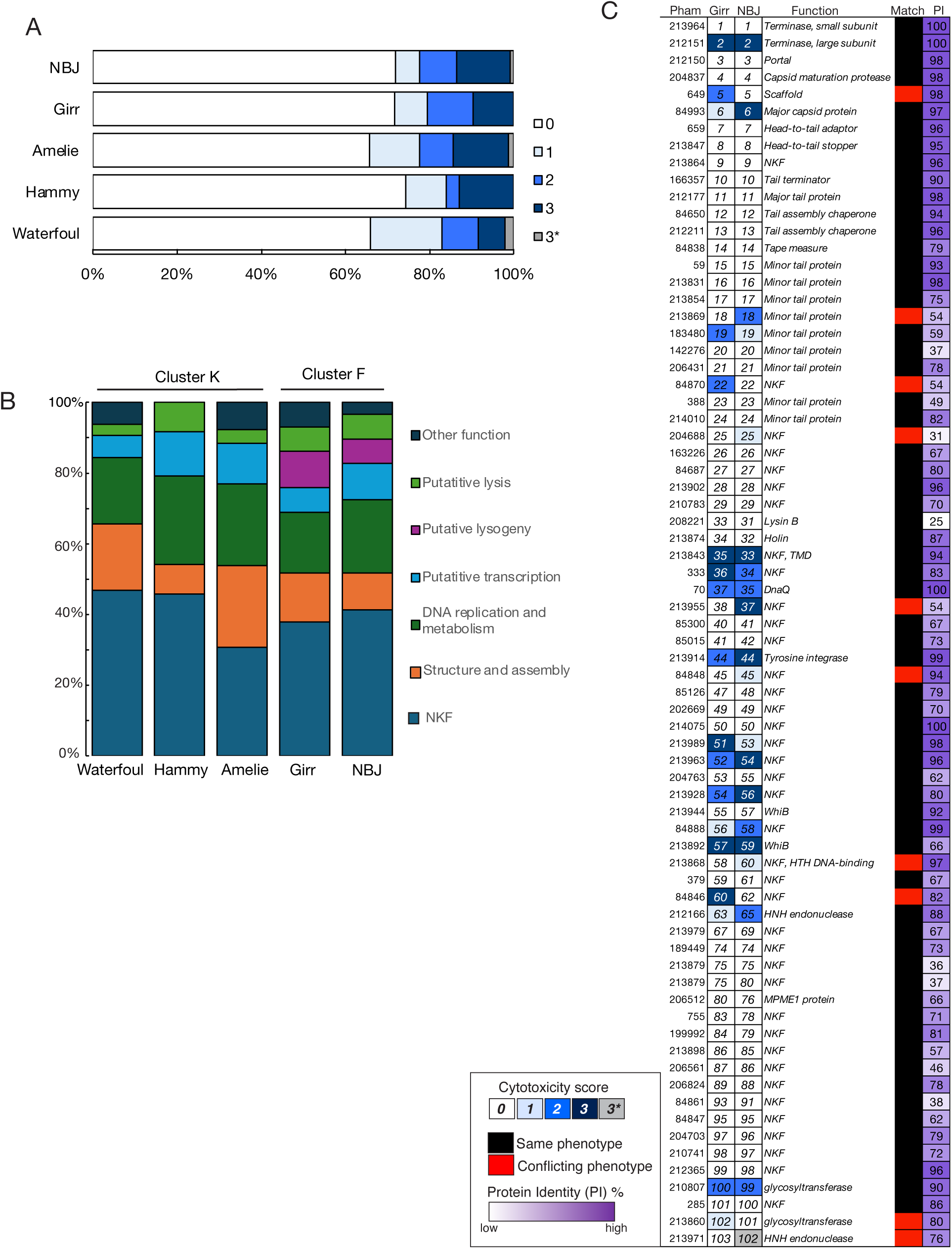
Conservation of observed cytotoxicity patterns. A) The proportion of genes from NBJ (this study), Girr (Pollenz et al. 2024), Amelie (Tafoya et al. 2024) , Hammy (Amaya et al. 2023), and Waterfoul (Heller et al. 2022) that were assigned each score (0–3 or 3*) is represented as a stacked bar chart. B) The proportion cytotoxic genes (score 1–3) that are NKF or that fall into various functional classes is represented as a stacked bar chart with different colors indicating functional class as described in the key. C) Shown is a chart listing the 72 gene phamilies shared by NBJ and Girr that were tested in similar cytotoxicity screens. Phamily number designations and functions are listed (NKF, no known function; TMD, transmembrane domain) next to representative homologous genes from NBJ and Girr, with boxes shaded by cytotoxicity score. A binary indicator of whether homologous genes were both classified as toxic or nontoxic is illustrated by black or red shading, and % protein identity is indicated by the purple gradient boxes.

Several gene products with predicted functions in DNA replication or DNA metabolism were also seen to cause a notable reduction in host viability upon overproduction, including the DnaQ-like protein gp35, RecA protein gp64, the tyrosine integrase gp44, HNH endonuclease gp65, and G-I-Y Y-I-G endonuclease gp70 (all score 2), as well as an encoded transposase gp43 (score 3). Four gene products with putative roles in gene regulation were also identified as cytotoxic: the mild growth inhibitors gp25 and gp60, which harbor predicted helix-turn-helix DNA-binding domains, the predicted immunity repressor protein gp47, and the WhiB-like protein, gp59. Overexpression of lysin A gene *30,* caused a moderate reduction in growth, adding to a growing list of mycobacteriophage Lysin A proteins that have been observed to mediate holin-independent toxicity when overexpressed in *M. smegmatis* (Payne and Hatfull 2012; Amaya et al. 2023).

Of the remaining 13 hits without annotated functions, DeepTMHMM analysis predicts that cytotoxic gene *33* (score *2*), located directly downstream of the annotated holin, encodes for a protein with a single pass transmembrane domain, whereas gp45 (score *1*) and gp56 (score *3*) are predicted to contain N-terminal signal sequences for export by the Sec system. As shown in Figure 2C, removal of the N-terminal signal sequence eliminated the cytotoxic phenotype of gp45 and gp56, suggesting that toxicity is linked to Sec-mediated export, either because these proteins act in the periplasm or cell envelope, or because overproduction disrupts the secretion pathway.

Despite multiple attempts, transformants could not be recovered for the pExTra plasmid encoding the predicted HNH endonuclease NBJ *102.* A nonsense mutation was introduced in the second codon position of the gene *102* sequence, and transformation of *M. smegmatis* with this mutated plasmid yielded a comparable number of transformants as the pExTra01 empty vector plasmid (Figure 2D). This result indicates that leaky expression of NBJ *102* from the pExTra plasmid prevents recovery of transformants, suggesting that 102 is likely highly cytotoxic (classified as score 3* in Figure 2). Therefore, in total, we identified 29 NBJ genes capable of inhibiting mycobacterial growth.

### Conservation of observed NBJ patterns of cytotoxicity in other mycobacteriophages

Our analysis indicates that 27% of the genes encoded in the NBJ genome can inhibit host growth, a comparable figure to the 28% of Girr genes found to inhibit host growth (Pollenz et al. 2024) as well as the 26-34% of genes found to be cytotoxic in similar whole-genome screens of Cluster K mycobacteriophages Waterfoul, Hammy, and Amelie (Heller et al. 2022; Amaya et al. 2023; Tafoya et al. 2024) (Figure 3A). As shown in Figure 3B, the functional profiles of the cytotoxic genes identified in these five genome-wide screens are also similar, with comparable representation of NKF genes as well as genes involved in structure and assembly, DNA replication and metabolism, transcription, and lysis functions. For example, the major capsid proteins from all five phages, which are classified based on amino acid similarity into two different phamilies, were all found to be cytotoxic upon overproduction, an intriguing observation given recent reports of major capsid proteins acting as triggers for bacterial abortive infection systems (Huiting et al. 2022; Zhang et al. 2022; Stokar-Avihail et al. 2023). Likewise, all five screens have identified a cytotoxic transmembrane protein encoded adjacent to the holin gene. Other conserved functions identified as being toxic in both Cluster F and Cluster K screens include putative replication proteins such as DnaQ (e.g., NBJ gp35 and Amelie gp49) and WhiB-like transcription factors (e.g., NBJ gp59 and Hammy gp53).

Genome-wide comparison of F1 phages NBJ and Girr reveals that phenotypic profiles are well-conserved across these closely related genomes (Figure 3C). In both phages, cytotoxic genes are encoded throughout the genome, with clusters of cytotoxic genes concentrated within the predicted early lytic region downstream of predicted *cro* genes (Ko and Hatfull 2018) and near the lysis cassettes. Additionally, whereas Cluster K genes involved in lysogeny were not observed to be cytotoxic in our system, for both F1 phages, the genes encoding the predicted tyrosine integrases and immunity repressors were found to cause severe growth defects when overexpressed. Downstream of the immunity repressors, both NBJ and Girr encode homologous proteins with signal peptides (gp45); however, only overproduction of NBJ gp45 was observed to cause mild toxicity.

Overall, as shown in Figure 3C, NBJ and Girr share 70% of their gene content, encoding 72 homologous phams (phagesdb.org). Of these, 62 phams (86%) exhibited consistent cytotoxicity phenotypes across the two screens, supporting their functional conservation. The remaining ten phams with discordant cytotoxicity results vary widely in amino acid identity, ranging from 31% (i.e., NBJ and Girr gp25) to 98% (i.e., NBJ and Girr gp5), leaving open whether these differences reflect true functional divergence or variation in expression levels within our system.

Finally, our NBJ screen identified eight cytotoxic gene phamilies not previously tested in other similar screens, including NBJ gp39 and gp43, encoded by reverse-strand genes near the tyrosine integrase, the RecA-like recombinase gp64, the orpham gp68, and gp70, gp71, and gp90 encoded within the right arm of the genome. These newly defined inhibitors highlight that even closely related phages can encode unique host-modulating factors, underscoring the value of extending systematic functional screens to phages across the spectrum of mycobacteriophage sequence diversity (Pope et al. 2015; Hatfull 2018) .

### Gene-mediated defense against phage infection

To further probe the ability of NBJ gene products to modulate host behaviors, we tested a subset of genes within our pExTra-NBJ library in an efficiency of plating assay to identify potential defense genes that can protect the host cell from phage infection. First, we sought to evaluate whether the predicted immunity repressor encoded by cytotoxic gene *47* can confer homotypic superinfection immunity upon overexpression as has been shown for other mycobacteriophage immunity repressors (Petrova et al. 2015) . *M. smegmatis* transformed with the pExTra-NBJ47 plasmid was challenged with NBJ and 9 other Cluster F phages in an efficiency of plating assay alongside a control strain harboring the pExTra03 control plasmid. Given that overexpression of *47* causes a severe growth defect in the host, we evaluated immunity only on plates without inducer, relying on leaky expression from the plasmid.

As shown in Figure 4A, basal expression of gene *47* was sufficient to cause a reduction in plaquing efficiency for three of the four F1 phages tested: NBJ and Akhila had ≥2-log reduction in efficiency of plaquing on the M. *smegmatis*/pExTra-NBJ47 lawn as compared to the *M. smegmatis*/pExTra03 control lawn, whereas infection by Girr was completely blocked. By contrast, F1 phage, TootsiePop, which encodes a divergent immunity repressor with only 39-44% sequence identity to the immunity repressors encoded by NBJ, Girr, and Akhila, was not impacted by basal expression of NBJ *47*.

**Figure 4:**
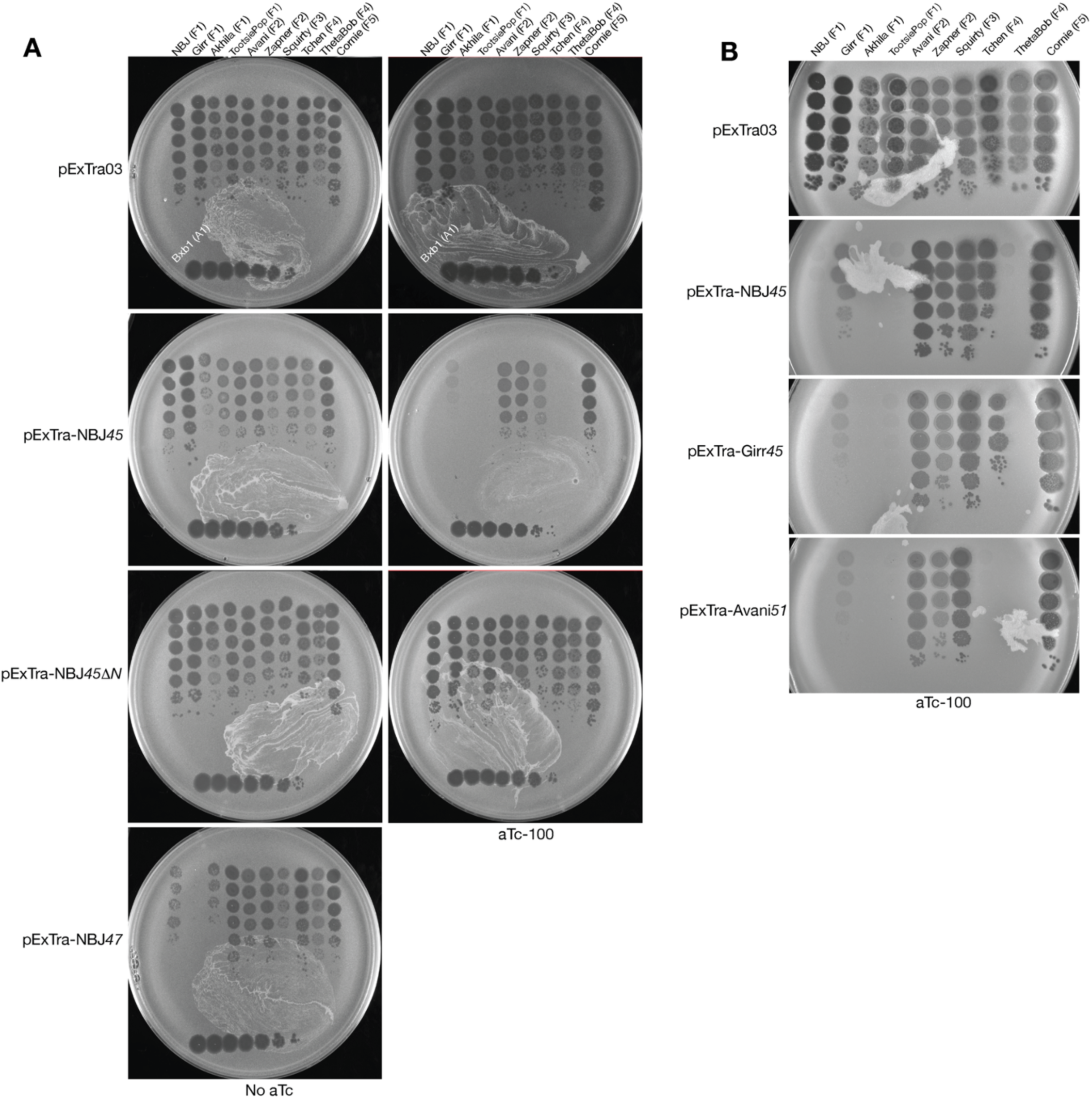
Gene-mediated defense against Cluster F phages. Strains of *M. smegmatis* mc^2^155 transformed with the indicated pExTra plasmids were combined with Middlebrook top agar and plated on 7H11 Kan with or without aTc. Ten-fold serial dilution series of lysates of ten Cluster F mycobacteriophages were spotted on these lawns to measure effects on the efficiency of plaquing with expression of NBJ genes (A) or homologous genes from Cluster F phages Girr and Avani (B).

Similarly, phages from other subclusters showed no notable difference in their ability to infect the host expressing NBJ *47,* though it is possible that broader immunity may be seen at higher expression levels.

In addition to traditional immunity repressor functions, temperate mycobacteriophages have been found to encode diverse phage-defense systems, with defense genes frequently clustered near other lysogeny genes in the genome center and expressed during lysogeny (Dedrick et al. 2017; Gentile et al. 2019; Montgomery et al. 2019). With this in mind, we next evaluated potential defense phenotypes for the subset of genes (*39, 40, 41, 42, 43, 45, 46*) flanking the NBJ integrase gene *44*. Cultures of *M. smegmatis* transformed with each pExTra plasmid or the pExTra03 control plasmid were used to form lawns on solid media with and without the aTc inducer. These strains were then challenged with a phage panel comprised of phages from Cluster F subclusters F1-F5 and five divergent clusters— A1/Bxb1, K2/ZoeJ, L1/LeBron, N/Charlie, P2/Tortellini, Y/Typha (Supplemental Figure 2).

Only gene *45* was observed to confer defense, with an aTc-dependent reduction in plaquing observed for the four F1 phages as well as two F4 phages, Tchen and ThetaBob. Plaquing efficiencies for F2 phages Avani and Zapner, F3 phage Squirty, and F5 phage Cornie were not impacted by expression of *45* (Figure 4A), nor were the plaquing efficiencies of the divergent phages (Supplemental Figure 2). Overproduction of the gp45 truncation variant missing the N-terminal signal peptide had no effect on phage plaquing, indicating that export to the cell surface is necessary for the observed defense phenotype.

Homologs of NBJ gp45 belonging to the same phamily are encoded by most sequenced Cluster F phages (phagesdb.org), and transcriptomic analysis of a lysogen of F1 phage Fruitloop demonstrated that NBJ *45* homologs are likely expressed in an operon with the immunity repressor during lysogeny (Ko and Hatfull 2018). In the study by Pollenz *et al*., overexpression of Girr *45* was not observed to cause cytotoxicity suggesting potential functional divergence within the pham (Figure 3C) (2024). Therefore, we sought to evaluate whether other pham members were able to confer a similar defense phenotype as NBJ gp45. Overexpression of homologous genes Girr *45* and Avani *51*, which encode products with >92% amino acid identity, were found to confer similar defense profiles against Cluster F phages, with aTc-dependent reduction of plaquing seen for F1 and F4 phages (Figure 4B). Together, these data indicate that NBJ encodes two conserved phage-defense modalities—homotypic immunity mediated by the immunity repressor gp47 and a secondary defense by secreted protein gp45 to block infection by both F1 and F4 phages.

### Structural predictions of host-modulating proteins

Because many cytotoxic NBJ proteins lacked sequence-level homologs, we turned to structure prediction by AlphaFold3 to identify remote functional relationships (Abramson et al. 2024). Figure 5A shows the predicted template modeling (pTM) scores for 15 cytotoxic NBJ gene products, including the 13 products with no known function and the two predicted HTH DNA-binding proteins, gp25 and gp60. pTM scores ranged from 0.28 (gp54) to 0.77 (gp25), with nine of the 15 proteins scoring greater than 0.5, indicating a high-confidence prediction. The best ranked model for each product was then queried using Foldseek (TM-align mode) to identify any informative structural homologs (TM-score >0.5) within the Protein Data Bank (PDB), Big Fantastic Viral Database (BFVD), and AlphaFold predicted structure database (AFDB) (Kempen et al. 2024; Kim et al. 2024). Gp39, the smallest cytotoxic gene product (48 amino acids) identified in our screen, had no structural matches in these three structural databases; similarly, gp54, the product with the lowest confidence prediction, did not yield any matches with TM-score >0.5. For all other gene products, not surprisingly, the highest scoring match was to another phage protein in the BFVD, in most cases, an uncharacterized hypothetical protein (Table 2). In the case of gp71, the top hit was to a predicted replisome organizer from an unclassified tailed phage, whereas the top hits for gp58, gp72, and gp90 were other uncharacterized actinobacteriophage proteins annotated as having some DNA-binding motifs (Table 2). gp25 and gp60 both aligned with DNA-binding proteins in the PDB or AFDB, supporting their putative roles in transcription. The highest scoring hit for gp25 was the DNA-binding domain of the *Bacillus subtilis* DeoR regulator, with significant alignment to region 4 domains of sigma factors from *Mycobacterium tuberculosis* (PDB: 2O8X, TM-score 0.539) and *Streptomyces coelicolor* (PDB: 5FGM, TM-score 0.558). gp60 aligned to DNA-binding domains from various prokaryotic, eukaryotic, and synthetic proteins as well as the AFDB predicted structure for a sigma 70 region 4-domain containing protein from *Gordonia terrae* (A0AAD0K934, TM-score 0.564).

**Figure 5:**
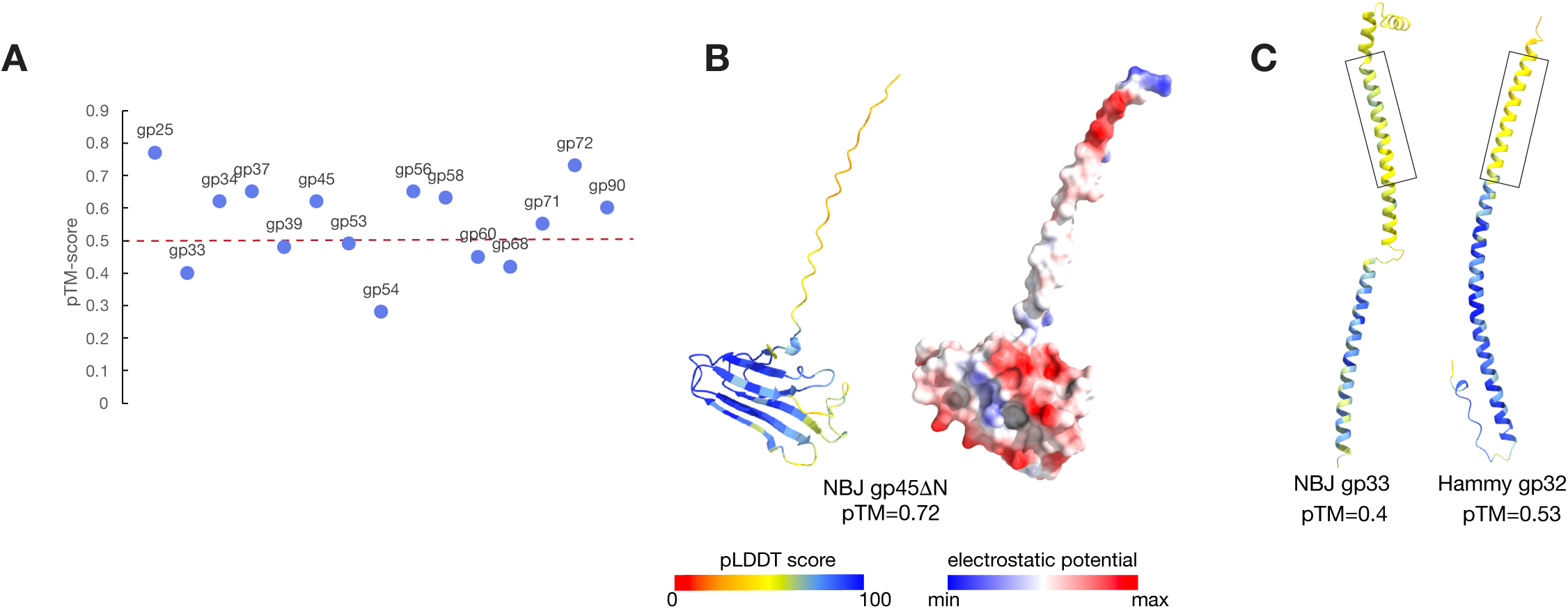
Structural predictions of NBJ cytotoxic NKF proteins. AlphaFold3 was used to predict monomeric structures of 15 cytotoxic NKF proteins. A) Shown is a plot of pTM scores for each structure with pTM >0.5 considered a high confidence structure. B) The predicted structure for NBJ gp45 lacking the N-terminal signal peptide is shown colored by pLDDT value (left) and electrostatic potential (right). C) Predicted structures for NBJ gp33 and Hammy gp32 are shown colored by pLDDT value with predicted transmembrane domains highlighted with black boxes.

**Table 2:** Structural homology of cytotoxic NBJ proteins of unknown function.

|  |  | Top Foldseek Prediction |  |  |  |  |  |  |  |  |
| --- | --- | --- | --- | --- | --- | --- | --- | --- | --- | --- |
| Gene Product | pTM | BFVD |  |  | AFDB-50 |  |  | PDB100 |  |  |
|  |  | Target | Prob. | TM-score | Target | Prob. | TM-score | Target | Prob. | TM-score |
| gp25 | 0.77 | A0A6J7WUJ2; Uncharacterized protein, uncultured Caudovirales phage | 0.99 | 0.827 | A0A1A1YEV0; Helix-turn- helix DNA binding protein from *Mycolicibacterium conceptionense* | 0.99 | 0.887 | 7bhy-assembly1_A; DNA-binding domain of DeoR from *Bacillus subtilis* | 0.8 | 0.650 |
| gp33 | 0.40 | No hit with TM-score >0.5 |  |  | No hit with TM-score >0.5 |  |  | No hit with TM-score >0.5 |  |  |
| gp34 | 0.62 | A5YK02; Uncharacterized protein, Mycobacterium phage Tweety | 0.85 | 0.642 | No hit with TM-score >0.5 |  |  | No hit with TM-score >0.5 |  |  |
| gp37 | 0.65 | A0A6G9LFF7; Uncharacterized protein, Mycobacterium phage Veteran | 0.97 | 0.682 | A0A100WIG6; Uncharacterized protein, gp38 *Mycolicibacterium canariasense* | 0.92 | 0.68 | No hit with TM-score >0.5 |  |  |
| gp39 | 0.48 | No hit with TM-score >0.5 |  |  | No hit with TM-score >0.5 |  |  | No hit with TM-score >0.5 |  |  |
| gp45 | 0.62 | Q19YL4; Uncharacterized protein, Mycobacterium virus PMC | 0.47 | 0.664 | A0AAU5EVU9; Uncharacterized protein, Streptomyces sp. NBC_00224 | 0.41 | 0.645 | 7wim-assembly3_N; *Arabidopsis thaliana* FKBP43 N-terminal domain | 0.25 | 0.509 |
| gp53 | 0.49 | A0A0F7LAN3; Uncharacterized protein, uncultured marine virus | 0.96 | 0.721 | A0A0C3Q0H7; Uncharacterized protein, *Tulasnella calospora* | 0.63 | 0.557 | No hit with TM-score >0.5 |  |  |
| gp54 | 0.28 | No hit with TM-score >0.5 |  |  | No hit with TM-score >0.5 |  |  | No hit with TM-score >0.5 |  |  |
| gp56 | 0.65 | A0A5Q2WIE3; DUF732 domain-containing protein, Streptomyces phage Daubenski | 0.85 | 0.668 | A0A0U1CXD5; DUF732 domain-containing protein, *Mycobacterium europaeum* | 0.77 | 0.746 | No hit with TM-score >0.5 |  |  |
| gp58 | 0.63 | S5XXU9; Helix-turn-helix DNA binding domain protein, Mycobacterium phage Hamulus | 0.87 | 0.646 | No hit with TM-score >0.5 |  |  | No hit with TM-score >0.5 |  |  |
| gp60 | 0.45 | A0A3G3LZ32; DNA binding protein, Mycobacterium phage Filuzino | 0.1 | 0.565 | A0AAD0K934; RNA polymerase sigma-70 region 4 domain protein, *Gordonia terrae* | 0.72 | 0.564 | 1tc3-assembly1_C; Transposase, *Caenorhabditis elegans* | 0.13 | 0.548 |
| gp68 | 0.42 | A0A068F2G8; Uncharacterized protein, Mycobacterium phage Gaia | 0.14 | 0.557 | A0A7C4VGC0; DUF1508 domain-containing protein, Pseudomonadota bacterium | 0.15 | 0.552 | No hit with TM-score >0.5 |  |  |
| gp71 | 0.55 | A0A8S5QGH6; Replisome organizer, unclassified Caudoviricetes | 0.10 | 0.534 | A0A931HR74; Uncharacterized protein, Rhodococcus sp. | 0.47 | 0.735 | No hit with TM-score >0.5 |  |  |
| gp72 | 0.73 | A0A8T8JGA1; HNH endonuclease, Mycobacterium phage Akhila | 1.00 | 0.782 | A0A1T8PBL3; DNA-binding phage zinc finger domain-containing protein, Mycobacteroides abscessus subsp. massiliense | 0.98 | 0.729 | No hit with TM-score >0.5 |  |  |
| gp90 | 0.60 | A0A8F2DAE6; Ribbon-helix-helix DNA binding domain protein, Gordonia phage VanLee | 0.60 | 0.558 | A0A5C5RQR7; DUF4254 domain-containing protein, *Tsukamurella sputi* | 0.98 | 0.793 | No hit with TM-score >0.5 |  |  |

For gp45, the predicted structure shows an unstructured tail of 24 residues at the new N-terminus created after cleavage of the predicted signal peptide followed by a globular C-terminal domain with a higher confidence fold (Figure 5B). Foldseek identified structural similarity between this folded C-terminal domain and nucleoplasmin domains from eukaryotic histone chaperones (PDB: 7WIM, TM-score 0.539). Nucleoplasmins are widely conserved proteins with a beta-barrel core that function as pentamers or decamers with negatively charged surfaces that interact with highly basic proteins like histones (Dutta et al. 2001). gp45 exhibits the hallmark nucleoplasmin eight strand beta-barrel core with extended acidic patches (Figure 5B), though AlphaFold3 predicts a pentameric structure of NBJ gp45 with relatively low confidence (ipTM 0.13, pTM 0.23).

Structural predictions for two of the conserved cytotoxic transmembrane proteins encoded adjacent to holin genes, NBJ gp33 and Hammy gp32, reveal extended alpha-helical structures with a single transmembrane domain near their N-termini (Figure 5C). Their genomic location and predicted structures are consistent with a potential relationship to the LysZ proteins recently described in phages of *Corynebacterium glutamicum* (McKitterick et al. 2025). LysZ proteins from corynebacteriophages are likewise encoded adjacent to annotated holin genes and predicted to form extended, membrane-associated helical structures (McKitterick et al. 2025).

### Most cytotoxic genes of unknown function are not necessary for lytic growth of NBJ

To assess whether cytotoxic NBJ genes were required for lytic growth, we used BRED and CRISPR-enhanced BRED (CRISPY-BRED) to generate targeted gene deletions and enrich for recombinant plaques (Wetzel et al. 2021). Recombinant plaques were identified by flanking PCR and confirmed by whole-genome sequencing (Supplemental Table 4). In total, single gene deletions were attempted for 16 cytotoxic NBJ proteins (Table 1, Supplemental Table 4). As expected, no plaques were recovered on CRISPR selection plates for the attempted deletion of major capsid gene *6* (Figure 6A); similarly, no plaques were recovered for deletion of gene *71* despite multiple attempts, suggesting that *71* is likely essential for NBJ lytic infection. Deletion of essential phage genes can sometimes be achieved when the gene is expressed from a plasmid in the host; however, in this case, we were unable to test complementation as plasmid-based expression of NBJ *71* completely abolishes host growth (Supplemental Figure 1).

**Figure 6:**
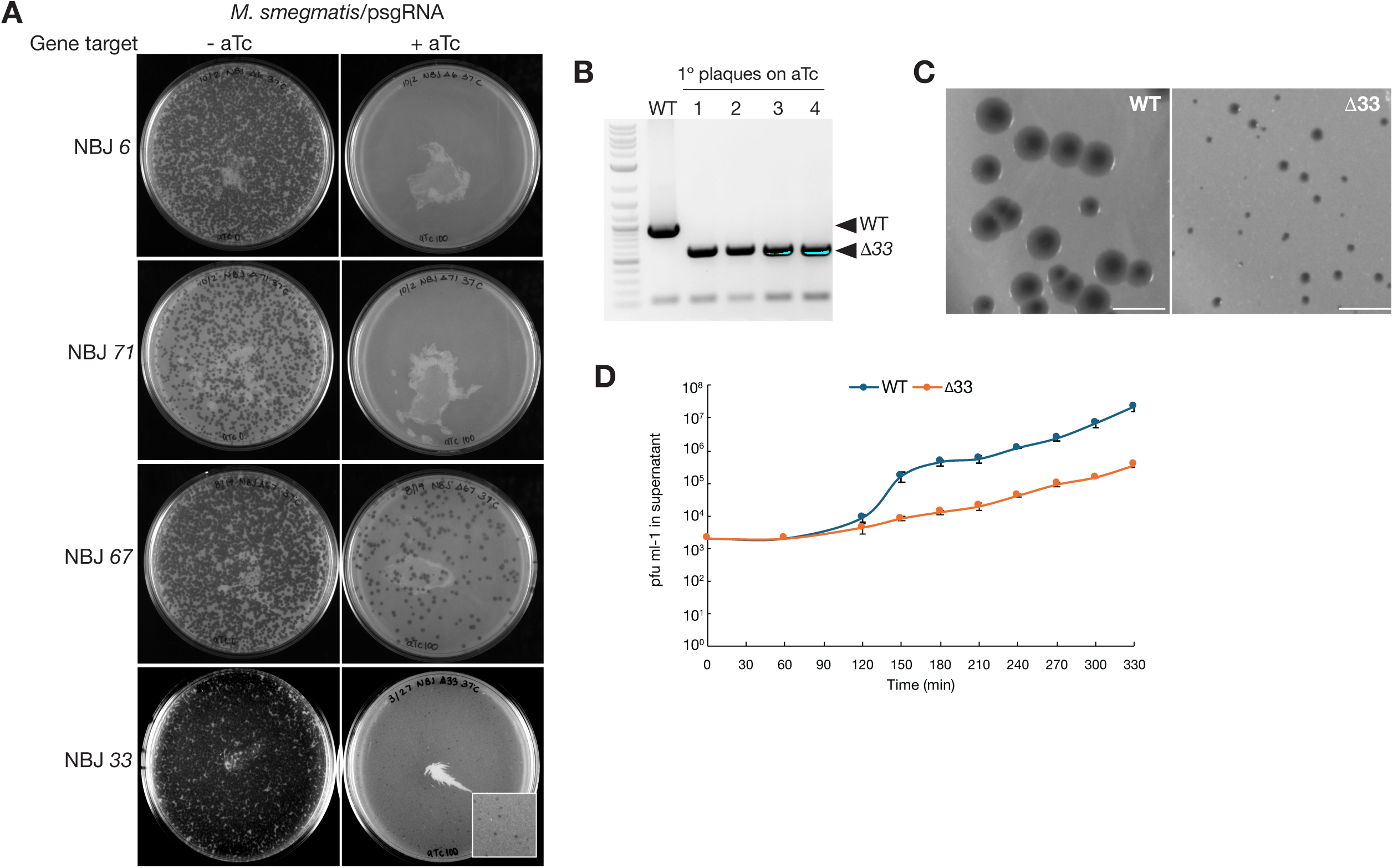
Gene deletion analysis of NBJ cytotoxic effectors. A) Recombineering strai*n M. smegmatis* mc^2^155/pJV138 (Wetzel et al. 2021) was electroporated with NBJ gDNA and the indicated deletion substrate, recovered for 4 h at 37 °C with shaking, and then combined with *M. smegmatis* bearing the relevant sgRNA-expressing plasmid. This mixture was split with half plated with top agar on 7H11 Kan plates without aTc inducer and the other plated with top agar on 7H11 Kan plates supplemented with 100 ng/ml aTc. Shown are the outcomes for a subset of NBJ genes tested. B) Plaques isolated on aTc-100 plates were picked and subjected to PCR with primers flanking the deletion site. Shown is an example PCR for deletion of gene *33,* where all four picked plaques display a single band corresponding to the expected size for the NBJΔ33 deletion mutant. C) Plaque size comparisons for wild-type NBJ and NBJΔ33. Plaques were generated from purified, high-titer lysates on wild-type *M. smegmatis* mc^2^155 on 7H10 plates and Middlebrook top agar and measured after 48 h of growth at 37 °C. D) One-step growth curve analysis comparing infection dynamics of wild-type NBJ and NBJΔ33 over 330 min. Each data point represents the average pfu/ml measured from triplicate cultures, and error bars represent standard deviation.

The remaining 14 cytotoxic genes tested were found to be non-essential for NBJ lytic growth (Table 1, Supplemental Figures 3 and 4), with mutant plaques readily isolated and the deletions confirmed by whole genome sequencing. Plaque size comparisons between the wild-type NBJ and deletion mutants showed no obvious difference in plaque size for 13 of the 14 mutants (Supplemental Table 4). This includes the immunity repressor gene *4*7 which is not predicted to be required for lysis based on its putative role in lysogeny, as well as the genes encoding for DnaQ-like protein gp35 and WhiB-like protein gp59, indicating that their putative roles in replication and transcription are not necessary for lytic growth under these conditions. This result is not surprising given similar findings from other studies showing that phages have a robust accessory gene pool that likely offers advantages under specific environmental conditions (Dedrick et al. 2013; Adler et al. 2025).

Notably, phage with the putative *lysZ-*like gene *33* deleted exhibited a significant reduction in plaque size (Figure 6C): wild-type NBJ plaques have an average diameter of 2.4 ± 0.5 mm, whereas *Δ33* plaques have an average diameter of 0.68 ± 0.2 mm. One-step growth curves of wild-type NBJ and the *Δ33* mutant show that at an MOI of 0.01, the wild-type phage exhibits an infection cycle with a latent period of ∼120 minutes and a burst size of ∼80 pfu (measured between 60 min and 150 min) (Figure 6D). The mutant on the other hand, shows an obvious lysis defect, with a gradual increase of pfu released into the supernatant over time rather than in a rapid orchestrated burst and >100-fold less pfu than the wild-type overall by 330 min post-infection (Figure 6D). A similar reduction in lysis efficiency was reported for *lysZ* deletion mutants in corynebacteriophages (McKitterick et al. 2025), and genetic analyses further determined they are necessary for efficient disruption of the LM/LAM layer of the *C. glutamicum* cell envelope during lysis. Together, these data are consistent with NBJ gp33 contributing to efficient lysis in Cluster F1 phages, a role that has since been confirmed and examined in greater detail in a follow-up study of the F1 lysis cassette (Pollenz et al. 2026).

### NBJ lysogen formation requires immunity repressor gp47

Like other Cluster F phages, NBJ is predicted to be temperate due to the presence of genes encoding for putative integrase (gp44) and immunity repressor functions within its genome. BLASTn alignment of the NBJ genome and the *M. smegmatis* mc^2^155 host identified the putative *attP* site, a 22 bp sequence (5’ gacccgctgattaagagtcagc) between NBJ genes *43* and *44* (Figure 1) that matches to one of two tRNA-Lys genes in the host genome; integration within bacterial tRNA genes is common for mycobacteriophages encoding tyrosine integrases (Hatfull 2018).

We next sought to confirm whether NBJ is a temperate phage capable of forming stable lysogens and to explore the potential roles of some of the cytotoxic genes identified in our screen. Lysogens were isolated by plating wild-type *M. smegmatis* on plates seeded with ∼10^9^ pfu of either wild-type NBJ or a subset of NBJ deletion mutants (Supplemental Table 4). Wild-type NBJ forms stable lysogens in the *M. smegmatis* host, with a frequency of lysogeny measured as 18-32% (average 22%) in five independent experiments. No lysogen formation was observed on the NBJ mutant lacking the predicted immunity repressor (NBJΔ*47*), confirming its essentiality in establishing lysogeny (Figure 7A).

**Figure 7:**
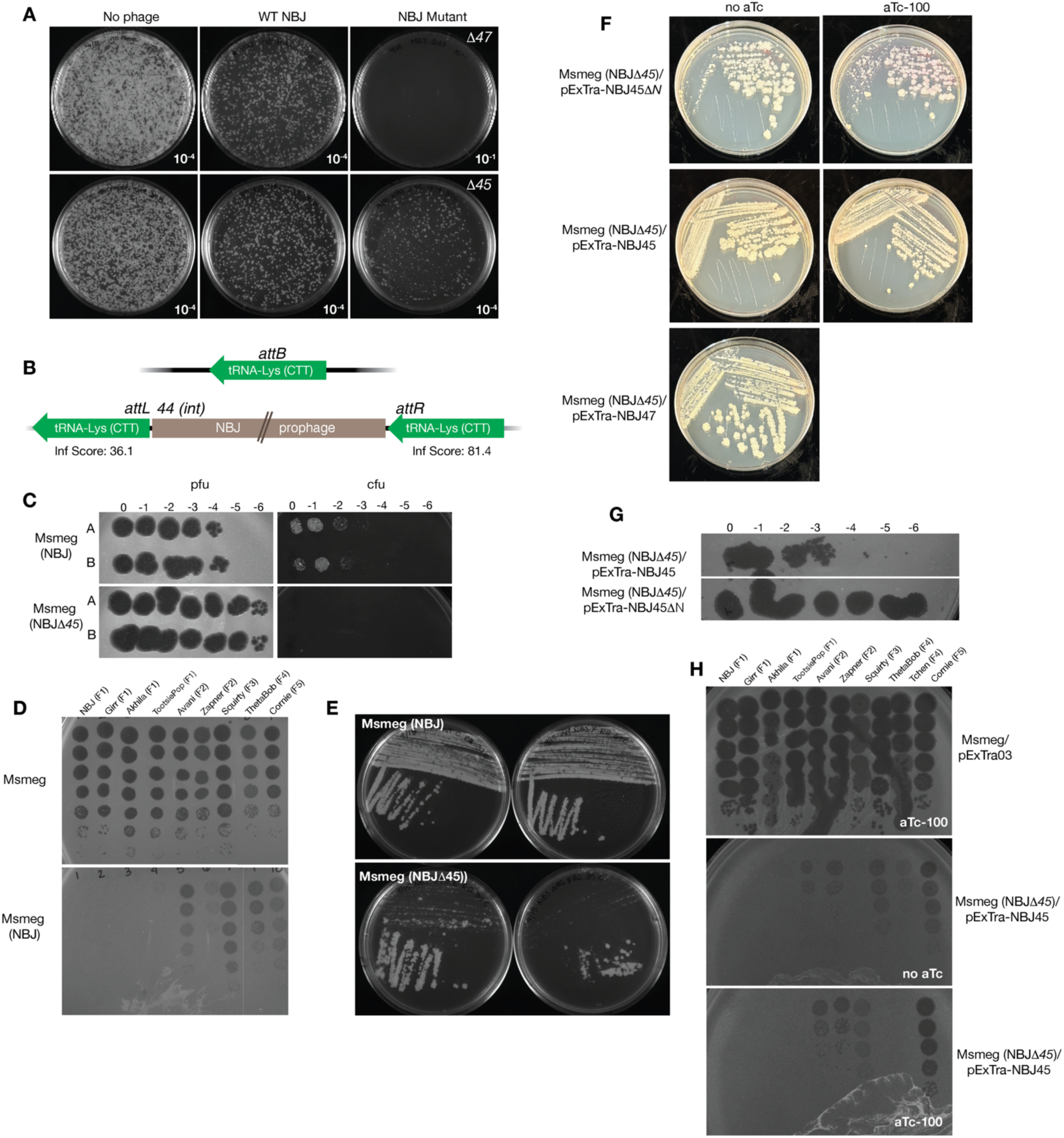
NBJ gp47 and gp45 are critical for lysogeny. A) Serial dilutions of an *M. smegmatis* mc^2^155 culture (O.D. 600 ∼2) were plated on 7H10 agar or on 7H10 agar seeded with ∼10^9^ plaque-forming units (pfu) of wild-type NBJ, NBJΔ45, or NBJΔ47. Colonies were apparent after 3 days incubation at 37 °C and frequency of lysogeny calculated as the number of colony-forming units (cfu) on seeded plates divided by the number of cfu on non-seeded plates. B) Integration of the NBJ genome into the *M. smegmatis* chromosome occurs within the tRNA-Lys gene. Integration of the prophage results in apparent duplication of the tRNA-Lys gene, with tRNAScanSE assigning the indicated inference (inf) scores for each. C) Cultures of the wild-type NBJ lysogen and NBJΔ45 lysogens were grown at 37 °C with shaking for 24 h. Cultures were serially diluted and plated on 7H11 plates to measure cell viability (right) and culture supernatants were titered on a wild-type *M. smegmatis* lawn to measure phage release by spontaneous induction (left). D) *M. smegmatis* or the wild-type NBJ lysogen strain were challenged with a panel of Cluster F phages to measure prophage-mediated immunity. E and F) Shown are streak plates of the indicated lysogen strains illustrating plaque formation and lyse-out effects seen in regions of high cell density. G) Cultures of the NBJΔ45 lysogen complemented with pExTra-NBJ45 or pExTra-NBJ45ΔN were grown at 37 °C with shaking for 24 h and culture supernatants were titered on a wild-type *M. smegmatis* lawn to measure phage release. H) The NBJΔ45 lysogen harboring the pExTra-NBJ45 plasmid was challenged with a panel of Cluster F phages in the presence or absence of aTc, with *M. smegmatis/*pExTra03 serving as a positive control for phage infection.

Based on gene expression data from lysogens of the Cluster F1 phage Fruitloop (Ko and Hatfull 2018), we also tested frequency of lysogeny for the subset of NBJ mutant phages with deletion of genes in regions predicted to be expressed during lysogeny; this includes gene *45*, which is expressed in a potential operon with the immunity repressor, and genes *59, 68, 72*, and *99* in the right arm of the genome where some expression was seen in the Fruitloop lysogen (Supplemental Table 4). A mild reduction in lysogen formation was observed for the NBJΔ45 mutant, which over three independent experiments was found to form lysogens at an average rate of 12%, ranging from a 9 to 19% decrease as compared to the wild-type (Supplemental Table 4, Figure 7A). No other mutants showed a consistent reduction in their frequency of lysogen formation.

Lysogens formed from wild-type NBJ and the NBJ*Δ45* mutant were carried through three rounds of streak purification, and the supernatants of pure lysogen cultures were found to contain high titers of phage particles, indicating spontaneous prophage induction and confirming lysogen isolation (Figure 7C). Long-read sequencing of the wild-type NBJ lysogen confirmed prophage integration within the within the host tRNA-Lys (Accession: CP00949.1: coordinates: 4,847,954- 4,848,029), with the prophage splitting the tRNA-Lys gene. tRNAscan-SE predicts tRNA-Lys genes at both *attL* (Inf score: 36.1) and *attR* (Inf score: 81.4) sites, indicating duplication of the host tRNA at the prophage boundaries (Figure 7B). When challenged with NBJ and other Cluster F phages in a plate-based immunity assay, the wild-type NBJ lysogen exhibited nearly complete immunity against NBJ and other homotypic F1 phages, including TootsiePop. It also had partial immunity against tested F2, F3, F4, and F5 phages, with a reduction in plaque size or moderate reduction in efficiency of plating observed for all (Figure 7D). The broader immunity observed for the wild-type NBJ lysogen than was seen with leaky plasmid expression could be due to a higher but sub-lethal level of *47* expression in the lysogen or a combinatorial effect caused by co-expression of *45, 47* and potentially other unidentified factors.

### NBJ gp45 is necessary for lysogen stability at high cell density

We next sought to evaluate the immunity profile of the lysogens formed by the NBJΔ45 mutant; however, as we purified lysogens raised on the NBJΔ45 mutant, an interesting phenomenon was observed. On streak plates for the wild-type lysogen, plaques were often observed within regions of high cell density, even after multiple rounds of purification away from any exogenous phage on the original selection plates. This phenomenon was far more pronounced for the NBJΔ45 mutant: streak plates exhibited complete clearing in the high density primary and secondary streaks, but viable individual colonies were still isolated in the least dense final streaks (Figure 7E). Furthermore, when grown in liquid culture for 24 h, both the wild-type and NBJΔ45 mutant showed signs of significant phage release, with the NBJΔ45 showing ∼100-fold more particle release than the wild-type lysogen (Figure 7C). Consistent with this higher level of phage release, cell viability counts from these cultures revealed that the wild-type lysogen maintained some viability whereas no viable colony forming units were detected for the NBJΔ45 lysogen.

These observations were surprising as lysogen cells should be immune to superinfection by phages released by spontaneous induction of neighboring cells. However, with significant phage release, local multiplicity of infection could be high enough to overwhelm the immunity repressor and lead to activation of lysis and productive rounds of phage release (Wiesmeyer 1966). The exacerbation of this phenomenon in the absence of gp45 production suggests it is necessary to maintain lysogen integrity in the presence of high levels of homotypic phage.

We next tested whether plasmid-based expression of *47* or *45* was sufficient to restore stability of the NBJΔ45 lysogen, with strains of *M. smegmatis* transformed with either the pExTra-NBJ47, pExTra-NBJ45 or pExTra-NBJ45ΔN plasmid used to raise lysogens of NBJΔ45. After two rounds of purification, we observed that on plates without aTc, NBJΔ45 lysogens harboring the pExTra-NBJ47 plasmid had far less pronounced signs of lysis than previously seen (Figure 7F) and could be grown to a high optical density in liquid, indicating that increased levels of the immunity repressor can reduce induction and prevent runaway lysis of lysogens. Similarly, NBJΔ45 lysogens overproducing the full-length gp45 from pExTra showed increased stability at high cell density whereas those lacking the N-terminal signal peptide showed significant lysis (Figure 7F) and had higher titers of phage in culture supernatants (Figure 7G). The NBJΔ45 lysogen was confirmed by long-read sequencing to have the prophage integrated into the same tRNA-Lys site as the wild-type. The immunity pattern of the NBJΔ45 lysogen complemented with pExTra-NBJ45 mimicked that of the wild-type lysogen on plates lacking aTc; only with addition of aTc was a considerable reduction in plaquing observed for F4 phages Tchen and Thetabob (Figure 7H), suggesting that defense against F4 phages requires higher levels of gp45 and may not be relevant under native lysogenic conditions.

Thus, our results indicate that although gp45 is not strictly required for the establishment of lysogeny, it is important for protecting the NBJ lysogen from secondary homotypic infection, a role that becomes especially critical when the lysogen is grown at high cell density. Gp45-mediated protection appears to be dependent on proper localization to the cell periplasm, consistent with a potential mechanism of surface-acting superinfection exclusion.

### Gp45 inhibits phage infection after adsorption and reduces early transcript accumulation

To determine which step of Cluster F phage infection is inhibited by gp45, we compared adsorption of NBJ to *M. smegmatis* carrying control gene plasmid pExTra03, pExTra-NBJ45, or pExTra-NBJ45ΔN. Cultures were grown in the absence of Tween and induced with 100 ng/ml aTc for 4 h prior to infection with NBJ at an MOI of 0.1. All three strains exhibited similar adsorption kinetics at 37 °C without shaking, with free phage titers decreasing over 45 min, from ∼10^8^ pfu/ml at time 0 to ∼10^7^ pfu/ml, corresponding to ∼80% adsorption (Figure 8A).

**Figure 8:**
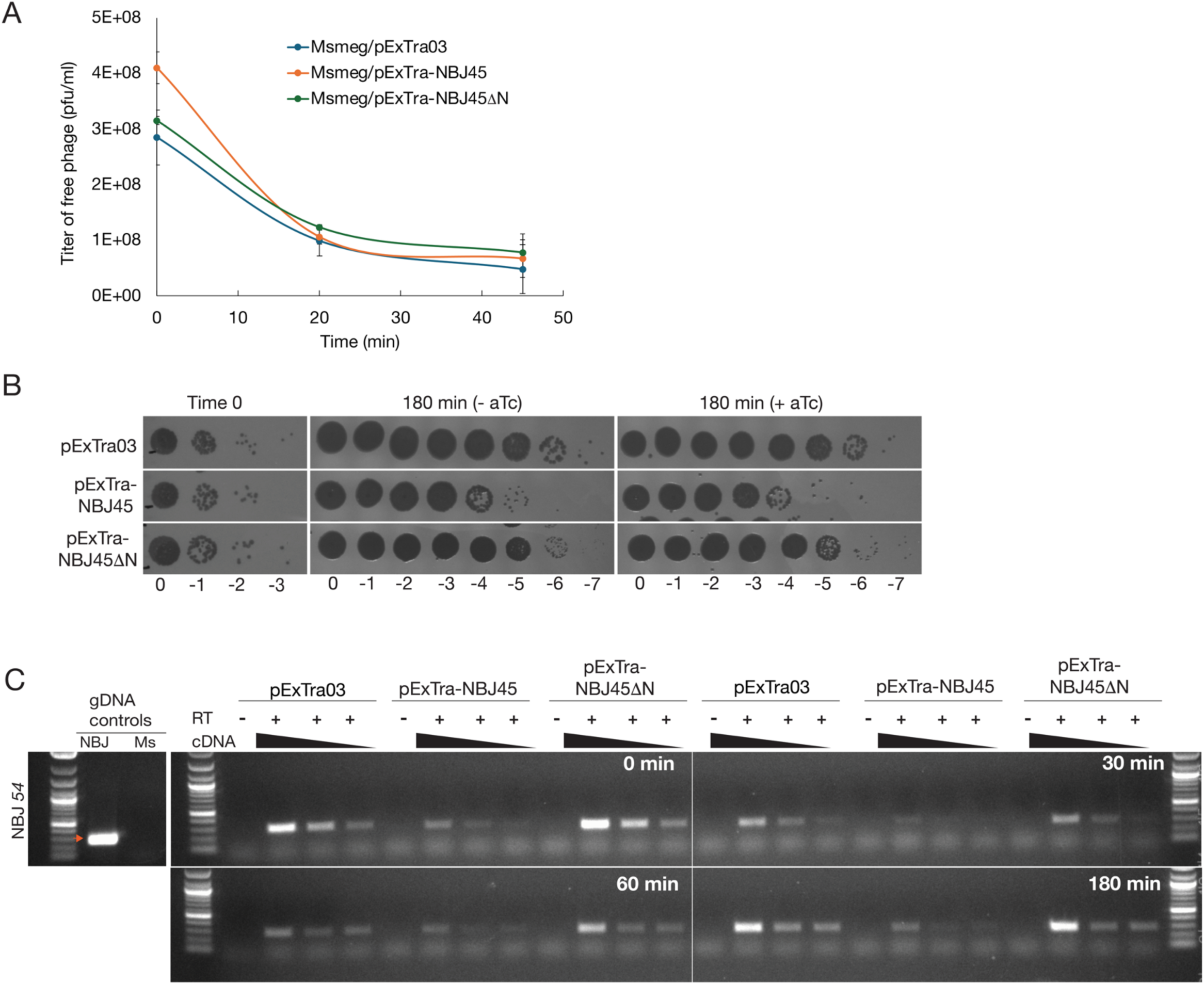
Gp45 inhibits NBJ infection post-adsorption and reduces early transcript accumulation. A) Adsorption of NBJ to *M. smegmatis* carrying control plasmid pExTra03, or pExTra expressing full-length NBJ *45* or NBJ *45ΔN.* Duplicate cultures were grown to an O.D.600 of 0.6 then induced with 100 ng/ml aTc for 4 h prior to infection. Induced cultures were mixed with NBJ at an MOI of 0.1, then incubated without shaking at 37 °C. Titers of free phage remaining in the supernatant were quantified by plaque assay over 45 min, with comparable adsorption kinetics observed across all three strains. B) After 45 min adsorption, cells were washed to remove unbound phage, and infection was allowed to proceed for 180 min in fresh medium either lacking or containing 100 ng/ml aTc. Representative spot assays of supernatants show reduced recovery of phage from the strain producing gp45 as compared to the pExTra03 or gp45ΔN control strains with a comparable defect observed whether or not aTc was present after washing. C) Semi-quantitative endpoint RT-PCR detecting an early NBJ transcript. Following adsorption and washing as described in A and B, cells from the same infected cultures were resuspended in medium containing 100 ng/ml aTc and samples collected at 0, 30, 60, and 180 min post-washing for RNA isolation. Total RNA was extracted, quantified, and normalized such that 500 ng of each sample was added to a combined DNase treatment and reverse transcription (RT) reaction or a matched DNase and no-RT control reaction. For each +RT sample, cDNA was assayed undiluted and serially diluted (1:5 and 1:25) by PCR using primers targeting early gene NBJ *54.* Under these non-saturating amplification conditions, the early NBJ *54* transcript remained detectable in all infected samples but was reduced in the gp45-producing strain relative to the pExTra03 and gp45ΔN controls.

To confirm whether gp45 inhibits infection after adsorption, infected cultures were washed after 45 min to remove unbound phage and resuspended in fresh medium either lacking or containing aTc. Cultures were then incubated at 37 °C with shaking for 180 min to allow infection to proceed, after which culture supernatants were titered to quantify productive infection. As shown in Figure 8B, strains producing full-length gp45 had a >2-log reduction in phage titer relative to either the pExTra03 or pExTra-NBJ45ΔN controls, indicating that gp45 inhibits NBJ infection after adsorption. A comparable defect was seen whether or not aTc was included after washing, indicating that continued gp45 production is not required and supporting a model in which gp45 acts at an early post-adsorption step in infection.

We next assessed accumulation of an early NBJ transcript in the same infected cultures using semi-quantitative endpoint RT-PCR on samples collected at 0, 30, 60, and 180 min post-wash. Using primers targeting NBJ *54*, a gene predicted to be expressed early in lytic growth based on the homologous locus in Cluster F1 mycobacteriophage Fruitloop (Ko and Hatfull 2018), we detected the expected amplicon in NBJ lysate DNA but not in *M. smegmatis* genomic DNA or matched no-RT controls, supporting that the observed signal was RNA-derived (Figure 8C). Under these non-saturating amplification conditions, the NBJ *54* transcript remained detectable in all infected samples but was consistently reduced in the gp45-producing strain relative to the pExTra03 and pExTra-NBJ45ΔN controls. This reduction in NBJ *54* transcript accumulation is consistent with the reduced, but not fully abolished, infection observed in liquid culture (Figure 8B) and indicates that gp45-mediated inhibition under these conditions is strong but incomplete, in contrast to the more complete defense against NBJ seen in plate-based assays. Together, these results indicate that gp45 acts after adsorption and before robust early transcript accumulation and productive infection.

### Escape genetics and bacterial two-hybrid analysis link gp45-mediated defense to a tape measure protein region

We next attempted to isolate phage defense escape mutants by infecting *M. smegmatis*/pExTra-NBJ45 with phage at an MOI of 10 and selecting for plaques in the presence of aTc. Despite repeated attempts, escape mutant plaques were not recovered for the F1 phages NBJ or Girr, but two escape plaques of Tchen were isolated. Whole-genome sequencing alongside the original Tchen lysate revealed that both escape mutants contained a single deviation from the wild-type sequence, encoding an arginine-to-serine substitution at position 310 in the tape measure protein gp15. The presence of this substitution was sufficient to overcome gp45 defense, restoring plaquing on a lawn expressing full-length *45* (Figure 9A). Mutant and wild-type Tchen showed comparable plaquing on a lawn of *M. smegmatis* expressing *45ΔN*, suggesting that the R310S substitution is not detectably detrimental to Tchen infection under these conditions (Figure 9A).

**Figure 9:**
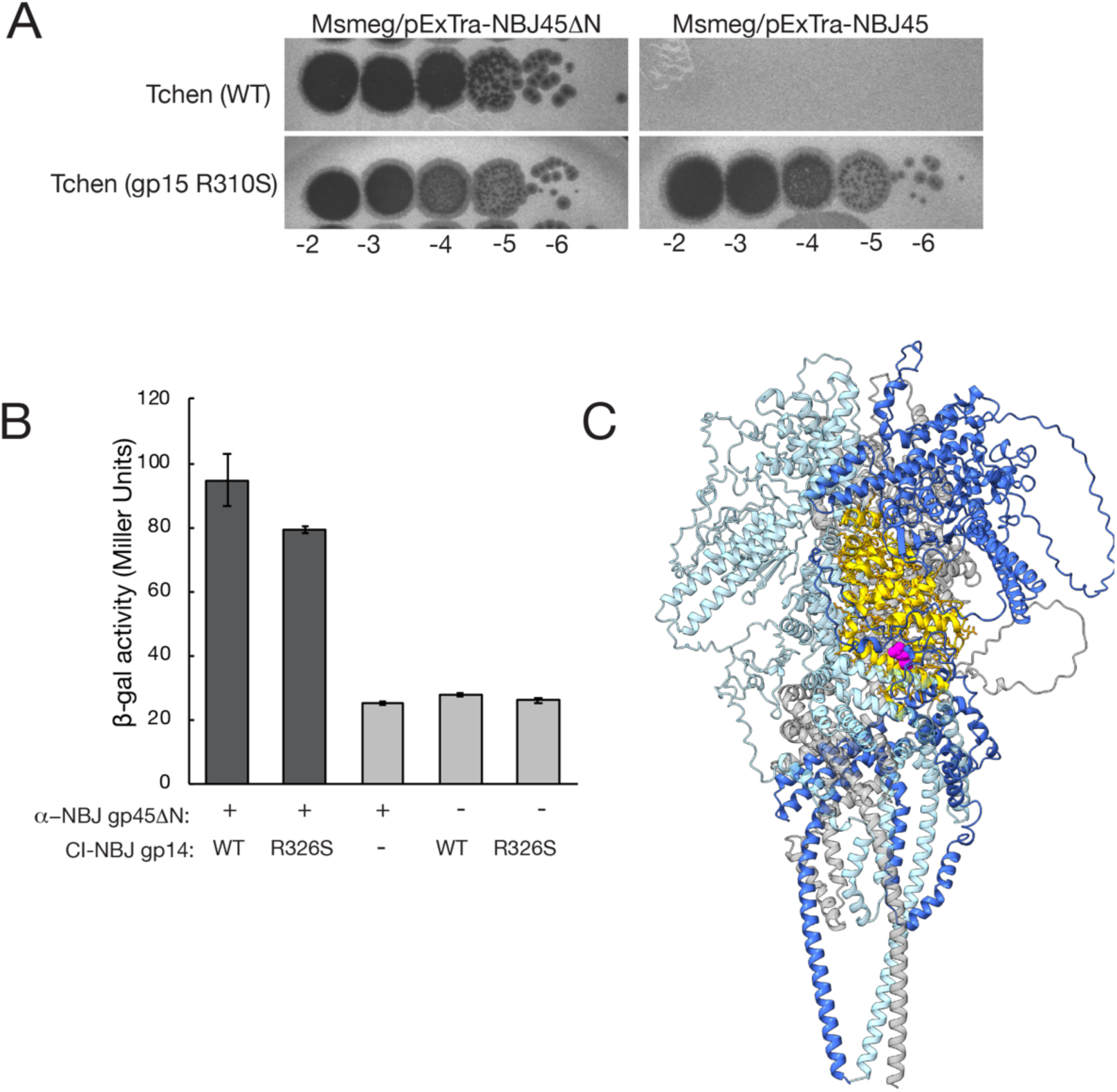
Escape genetics and bacterial two-hybrid analysis link gp45-mediated defense to a region of the Cluster F tape measure proteins. (A) Representative spot assays showing plaquing of wild-type Tchen and the isolated escape mutant carrying the gp15 R310S substitution on lawns of *M. smegmatis* expressing NBJ *45ΔN* or full-length NBJ *45*. (B) Bacterial two-hybrid assay measuring interaction between NBJ gp45ΔN and an NBJ gp14 tape measure protein fragment spanning residues 144–345. Bars show average β-galactosidase activity for three independent replicates of the wild-type gp14 fragment, the corresponding gp14 R326S mutant fragment, and negative controls. Error bars represent the standard deviation. (C) AlphaFold3 prediction of the NBJ gp14 tape measure protein trimer. The gp14 region tested in the bacterial two-hybrid assay (residues 144–345, gold) and the residue corresponding to the Tchen escape substitution (R326, magenta) are highlighted on one protomer.

Tape measure proteins are critical for the assembly of the phage tail and are involved in DNA transfer during infection (Cumby et al. 2015). They have also been previously identified as targets of phage-encoded superinfection exclusion systems, including HK97 gp15 and the lactococcal Ltp lipoproteins (Bebeacua et al. 2013; Cumby et al. 2015). We therefore next tested whether gp45 interacts with the tape measure region implicated by the Tchen escape mutation. Using a transcription-based bacterial two-hybrid system (Dove et al. 1997; Fu et al. 2004) , we measured interaction between NBJ gp45ΔN and an NBJ gp14 tape measure protein fragment spanning residues 144–345, which includes the residue corresponding to the Tchen escape substitution (NBJ gp14 R326). As shown in Figure 9B, the wild-type gp14 fragment interacted with gp45ΔN, resulting in a ∼3-fold increase in *lacZ* expression relative to the vector controls as measured by β-galactosidase assay. This result indicates that the escape-associated substitution lies within a region of the tape measure protein that can bind gp45. However, the corresponding R326S substitution caused only a modest reduction in this interaction (Figure 9B), suggesting that gp45-escape is not simply explained by a loss of interaction.

## Discussion

Here we present a systematic functional analysis of Cluster F1 mycobacteriophage NBJ to define the gene products that can influence *M. smegmatis* physiology over the course of phage infection. Using an arrayed, inducible whole-genome library of NBJ genes, we identified 29 genes, representing 27% of the genome, that inhibit host growth when overexpressed. This dataset contributes to the expanding functional genomics resource developed through the SEA-GENES undergraduate research project (Heller et al. 2024), reinforcing the patterns of growth inhibition seen in similar genome-wide screens of Cluster F and Cluster K mycobacteriophages. The consistent cytotoxicity caused by various DNA metabolism proteins, transcriptional regulators, and structural proteins suggests that many phage genes can disrupt essential host processes when expressed outside their native infection context, though it remains to be determined whether such perturbations reflect biologically relevant interactions or are simply a consequence of dysregulated protein expression. At the same time, the discovery of eight novel cytotoxic phamilies in NBJ, including the three potent growth inhibitors gp68, gp71, and gp72, underscores the value of extending these systematic screens to additional phages of varying evolutionary relatedness to discover novel interactions with the host cell.

Many of the modulatory proteins identified in our screen are proteins whose functions cannot be predicted using sequence-homology based methods, including orpham gp68, which has no close sequence homologs identified to date. As has been seen in other mycobacteriophages, many of these cytotoxic NKF genes are clustered near the lysis cassettes and within the early lytic region of the genome, making them especially good candidates for factors involved in the reprogramming and disruption of essential host machinery during lytic infection. To further explore these cytotoxic products and their potential roles in NBJ infection, we took two approaches: we performed a structure-based homology search to detect more distant evolutionary relationships missed by sequenced-based comparisons and used phage recombineering to evaluate gene essentiality.

As was seen by Guo et al., in their recent application of structural homology-based annotation of mycobacteriophage genomes (Guo and He 2025), structural modeling followed by Foldseek analysis yielded some informative hits that may improve functional annotations, though most of these were to other hypothetical viral proteins rather than experimentally characterized homologs. For example, the top hit for NBJ gp71 was an annotated replisome organizer from an unclassified siphovirus. Although the match probability was low, gp71 is encoded in the early lytic region of the genome near genes involved in DNA replication and DNA metabolism as are its homologous pham members encoded in more than half of Cluster F phages (phagesdb.org). This, together with our inability to recover *Δ71* mutants, supports a putative role in replication, a common functional class identified in mycobacteriophage cytotoxicity screens, though more experimental evidence is required.

Our structural homology search yielded no hits with a TM-score >0.5 for NBJ gp33, though the extended helical structure modeled by AlphaFold3, together with its synteny near the holin protein suggests that gp33 belongs to a broader class of small membrane-associated lysis proteins in actinobacteriophages. Overproduction of the corresponding LysZ proteins from corynebacteriophages Cog and CL31 was shown to inhibit growth of *C. glutamicum* while deletion of these genes caused a reduction in lysis efficiency on both solid and liquid media (McKitterick et al. 2025) . Structural homologs of gp33 with divergent sequences have been consistently identified in mycobacteriophage cytotoxicity screens, and the severe lysis defect seen for the NBJΔ33 mutant is consistent with a role in promoting efficient lysis of the mycobacterial cell. Indeed, since our initial report, subsequent analysis of the Cluster F1 lysis cassette has demonstrated that NBJ gp33 (renamed LysF1b) is one of two distinct holin-like proteins encoded in this region, functioning in concert with the upstream holin-like protein NBJ gp32 (renamed LysF1a) to mediate lysis. Importantly, in this follow-up study, infection of an *M. smegmatis* strain lacking lipomannan and lipoarabinomannan only modestly improved plaque size and did not fully rescue the *lysF1b* deletion defect, arguing that the F1 proteins may not act through the same LM/LAM-targeting mechanism proposed for corynebacteriophages (Pollenz et al. 2026). Further work will be needed to determine how broadly conserved the mechanisms of these actinobacteriophage lysis proteins are.

Our gene deletion analysis indicates the rest of these putative NKF host modulators are not required for lysis and plaque formation under standard laboratory conditions, nor are genes with established functions in transcription (WhiB gp59) and replication (DnaQ gp35). Therefore, it remains an open question what specific roles many of these cytotoxic genes play in the phage life cycle. The WhiB-like protein from Cluster K phage TM4 was previously shown to be toxic when overexpressed in *M. smegmatis* due to interference with the host WhiB2 protein involved in regulation of cell division; the phage encoded WhiB was found to bind within the promoter region upstream of the *whiB2* gene, leading to a downregulation of its expression (Rybniker et al. 2010). It is plausible that these phage-encoded WhiB proteins and other growth inhibitors may not be strictly required for phage lysis but interfere with host pathways to improve phage outcomes under specific conditions.

NBJ is a temperate phage confirmed to form lysogens, and two of the growth inhibitors identified in our screen were found to be critical for lysogeny—immunity repressor gp47 and predicted secreted protein gp45. In our cytotoxicity assay, overproduction of gp47 fully abolished *M. smegmatis* growth, even at the lower induction level, suggesting that precise regulation of its expression is critical for lysogen viability. A comparable phenotype was previously observed for the homologous repressor Girr gp46 (97% amino acid identity), where cytotoxicity persisted even when the predicted helix–turn–helix DNA-binding domain was deleted (Pollenz et al. 2024). This suggests that repressor-mediated toxicity is not likely due to DNA binding or repression of host transcription but rather some other interference with host processes, reminiscent of the lambda CII transcriptional regulator which inhibits DNA replication and is also highly toxic when overproduced (̦Ke dzierska et al. 2003). These observations emphasize that temperate repressors must be precisely regulated to balance immunity and host viability.

In our functional assays, basal production of gp47 from the pExTra plasmid was sufficient for homotypic immunity against F1 phages with highly similar immunity repressors, though the plasmid-based immunity observed was more limited than that displayed by the lysogen. In contrast, aTc-dependent overexpression of neighboring gene *45* conferred a distinct, secretion-dependent phenotype that extended protection to both F1 and F4 phages. Notably, here too the immunity pattern observed in the NBJ lysogen did not fully match the defense profile produced by gp45 overproduction, suggesting that its expression may also be tightly regulated in concert with operon partner *47* (Ko and Hatfull 2018), and that the observed lysogen immunity emerges from the combined action of gp47, gp45, and potentially other factors. Consistent with this model, deletion of *45* results in pronounced lysogen instability and killing, indicating gp45 is necessary for lysogen stability under high phage load. Interestingly, the F2 phages Avani and Zapner as well as the F3 phage Squirty, all of which were not inhibited by NBJ gp45 in our defense assay, encode their own gp45 homologs (phagesdb.org). Overexpression of Avani gp51 in *M. smegmatis* conferred defense against both F1 and F4 phages but not homotypic defense, suggesting that while NBJ gp45 functions to stabilize the NBJ lysogen, related homologs in other clusters may function primarily to provide broader defense against heterotypic infection.

Gp45 and its homologs harbor an N-terminal secretion signal, and our data indicate that gp45-mediated protection depends on export to the cell periplasm, consistent with a superinfection exclusion mechanism. Taken together, our adsorption, post-adsorption infection, and transcript analyses place gp45 action at an early stage of infection after phage binding has occurred. Rather than fully blocking infection under all conditions, gp45 appears to mediate a strong but incomplete defect that limits the development of productive infection and reduces accumulation of early phage transcripts. Our genetic and interaction data further connect this phenotype to the Cluster F tape measure proteins.

Defense escape by the F4 phage Tchen mapped to a single amino acid substitution (R310S) in tape measure protein gp15, while the corresponding region of the NBJ tape measure protein was sufficient to interact with gp45 in a bacterial two-hybrid assay. An AlphaFold3 model of the NBJ gp14 trimer, generated based on the trimeric organization reported for the Bxb1 tape measure protein (Freeman et al. 2025), provides a predicted structural context for the tape measure region implicated by the escape and two-hybrid results (Figure 9C). The modest effect of the analogous R326S substitution on the measured two-hybrid interaction suggests that escape is not simply explained by loss of interaction, but instead likely reflects disruption of a gp45-sensitive state or activity of the tape measure protein during an early infection step.

Structural modeling and Foldseek analysis of gp45 yielded a moderate-confidence homology to nucleoplasmin-like β-barrel domains typically found in eukaryotic histone chaperones. Although this similarity is not sufficient to infer direct homology or function, it is consistent with a growing theme in bacterial immunity, where many recently discovered anti-phage defense systems exhibit distant structural relationships to eukaryotic counterparts, often innate immunity proteins (Ledvina and Whiteley 2024). Further work will be needed to determine whether gp45 adopts a nucleoplasmin-like oligomeric state and whether that organization is important for its defense activity.

Altogether, this work further highlights how temperate mycobacteriophages encode a diverse repertoire of gene products that influence host physiology during both lytic and lysogenic growth. It also expands the SEA-GENES functional genomics dataset, underscoring the value of integrating systematic experimental screens with structural and comparative genomics approaches to better define the diverse biological roles encoded within phage genomes.

### Data availability statement

All plasmids, plasmid sequences, and phage sequences reported in this study are available upon request. The authors affirm that all data necessary for confirming the conclusions of this article are represented fully within the article and its tables and figures. Extended data, including plasmid Sanger sequencing data and confirmatory cytotoxicity assay data can be found at the SEA-GENES project open access database GenesDB (https://genesdb.org). To access these data on genesDB.org, any user can register for a free account, and once logged in to this account, navigate from the home page to Cluster F1 phage NormanBulbieJr.

## Acknowledgements

We are grateful to the members of the Science Education Alliance for their invaluable research support, particularly Viknesh Sivanathan, Denise Monti, James Melton, Billy Biederman, Graham Hatfull, Deborah Jacobs-Sera, Steven Cresawn, and Dan Russell. We also thank New England Biolabs (NEB) and Integrated DNA Technologies (IDT) for generously providing reagents. We also thank all the University of South Florida SEA-GENES students from the 2021-2024 cohorts for their work on F1 phage Girr and F2 phage Avani and for providing plasmid reagents and preliminary data to support the work in this report.

