## Supplemental Materials for "Identification and characterization of host-modulating effectors encoded by the Cluster F1 mycobacteriophage NormanBulbieJr"

**Supplemental Figure 1: Cytotoxicity screening.** Shown are the results of representative cytotoxicity assays for the NBJ genes screened in this study. Each strain was spotted in triplicate alongside *M. smegmatis*/pExTra-Fruitloop52 (+) and pExTra-Fruitloop52I70S (-) control strains on 7H11 Kan supplemented with 0, 10, or 100 ng/ml aTc. In all experiments, $10^{-1}$ to $10^{-5}$ dilutions are shown; in some experiments the undiluted sample was also spotted. Plates were monitored over 3 or 4 days at 37 ºC, with results shown to best illustrate effects on colony color and size. Colony color was scored using the indicated key.

**Supplemental Figure 2: Defense screening.** Shown are the results of defense assays for a subset of NBJ genes. Cultures of transformed *M. smegmatis* were plated with top agar on 7H11 Kan supplemented with 0 or 100 ng/ml aTc. The indicated phage lysates were spotted on replicate experimental lawns as well as a control lawn containing pExTra03 to compare efficiency of plaquing with gene expression.

**Supplemental Figure 3: Testing efficiency of psgRNAs.** Shown are the results of NBJ (F1), ZoeJ (K2) and Adephagia (K1) lysates spotted on *M. smegmatis/*psgRNA lawns *to* evaluate the ability of the designed sgRNA to specifically target wildtype NBJ and reduce plaquing efficiency.

**Supplemental Figure 4: Using CRISPY-BRED to isolate mutant phages.** Shown are the results of plaque assays performed by mixing *M. smegmatis*/pJV138 electroporated with NBJ gDNA and the indicated deletion substrate after 4 h recovery with the selection strain *M. smegmatis* /psgRNA. Lawns were plated with Middlebrook top agar on 7H11 Kan supplemented with 0 or 100 ng/mL aTc.

**Supplemental Figure 5: PCR Verification of mutant phages.** Shown are the flanking PCR results for plaques picked from7H11 Kan aTc-100 plates after BRED and CRISPR selection. Once the presence of the mutant was confirmed (smaller band), at least one round of streak purification was performed and PCR results confirmed.

### Supplemental Figure 1

Images taken after 3 days at 37 ºC on 7H11 agar

Gene *1*; Score *0*

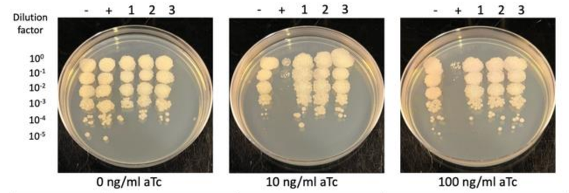


| Lane | Plasmid name | Gene name, replicate | Toxic/Non-toxic | Colony color on 100 ng/ml aTc plate* |
|---|---|---|---|---|
| - Non-toxic control | pExTra03 | Fruitloop *52* mutant | Non-toxic | + |
| + Toxic control | pExTra02 | Fruitloop *52* | Toxic | - |
| 1 | pExTra-NormanBulbieJr1 | NormanBulbieJr *1* replicate 1 | Non-toxic | + |
| 2 | pExTra-NormanBulbieJr1 | NormanBulbieJr *1* replicate 2 | Non-toxic | + |
| 3 | pExTra-NormanBulbieJr1 | NormanBulbieJr *1* replicate 3 | Non-toxic | + |

*Key: NG (no growth) - (no pink color) +(faint pink color) ++(obvious pink color) +++ (dark pink color)

Images taken after 3 days at 37 ºC on 7H11 agar

Gene *2*; Score *3*

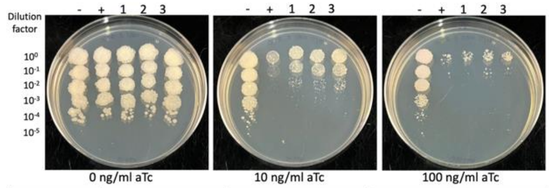


| Lane | Plasmid name | Gene name, replicate | Toxic/Non-toxic | Colony color on 100 ng/ml aTc plate* |
|---|---|---|---|---|
| - Non-toxic control | pExTra03 | Fruitloop *52* mutant | Non-toxic | + |
| + Toxic control | pExTra02 | Fruitloop *52* | Toxic | NG |
| 1 | pExTra-NormanBulbieJr2 | NormanBulbieJr *2* replicate 1 | Toxic | - |
| 2 | pExTra-NormanBulbieJr2 | NormanBulbieJr *2* replicate 2 | Toxic | - |
| 3 | pExTra-NormanBulbieJr2 | NormanBulbieJr *2* replicate 3 | Toxic | - |

*Key: NG (no growth) - (no pink color) +(faint pink color) ++(obvious pink color) +++ (dark pink color)

Images taken after 3 days at 37 ºC on 7H11 agar

Gene *3*; Score *0*

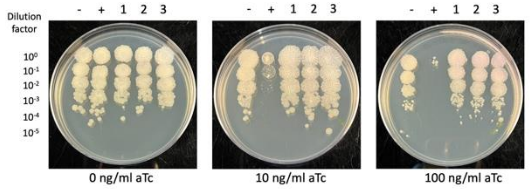


| Lane | Plasmid name | Gene name, replicate | Toxic/Non-toxic | Colony color on 100 ng/ml aTc plate* |
|---|---|---|---|---|
| - Non-toxic control | pExTra03 | Fruitloop *52* mutant | Non-toxic | - |
| + Toxic control | pExTra02 | Fruitloop *52* | Toxic | - |
| 1 | pExTra-NormanBulbieJr3 | NormanBulbieJr *3* replicate 1 | Non-toxic | + |
| 2 | pExTra-NormanBulbieJr3 | NormanBulbieJr *3* replicate 2 | Non-toxic | + |
| 3 | pExTra-NormanBulbieJr3 | NormanBulbieJr *3* replicate 3 | Non-toxic | + |

*Key: NG (no growth) - (no pink color) +(faint pink color) ++(obvious pink color) +++ (dark pink color)

Images taken after 3 days at 37 ºC on 7H11 agar

Gene *4*; Score *0*

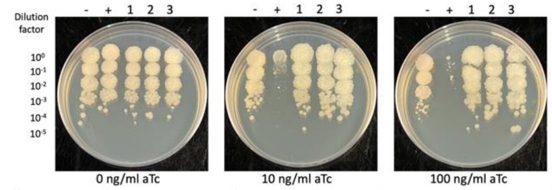


| Lane | Plasmid name | Gene name, replicate | Toxic/Non-toxic | Colony color on 100 ng/ml aTc plate* |
|---|---|---|---|---|
| - Non-toxic control | pExTra03 | Fruitloop *52* mutant | Non-toxic | + |
| + Toxic control | pExTra02 | Fruitloop *52* | Toxic | - |
| 1 | pExTra-NormanBulbieJr4 | NormanBulbieJr *4* replicate 1 | Non-toxic | - |
| 2 | pExTra-NormanBulbieJr4 | NormanBulbieJr *4* replicate 2 | Non-toxic | - |
| 3 | pExTra-NormanBulbieJr4 | NormanBulbieJr *4* replicate 3 | Non-toxic | - |

*Key: NG (no growth) - (no pink color) +(faint pink color) ++(obvious pink color) +++ (dark pink color)

Images taken after 3 days at 37 ºC on 7H11 agar

Gene *5*; Score *0*

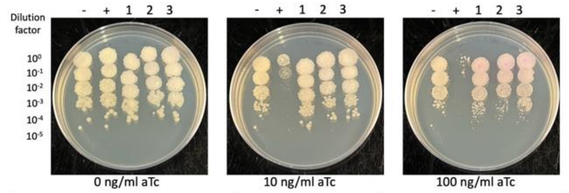


| Lane | Plasmid name | Gene name, replicate | Toxic/Non-toxic | Colony color on 100 ng/ml aTc plate* |
|---|---|---|---|---|
| - Non-toxic control | pExTra03 | Fruitloop *52* mutant | Non-toxic | - |
| + Toxic control | pExTra02 | Fruitloop *52* | Toxic | - |
| 1 | pExTra-NormanBulbieJr5 | NormanBulbieJr *5* replicate 1 | Non-toxic | + |
| 2 | pExTra-NormanBulbieJr5 | NormanBulbieJr *5* replicate 2 | Non-toxic | + |
| 3 | pExTra-NormanBulbieJr5 | NormanBulbieJr *5* replicate 3 | Non-toxic | + |

*Key: NG (no growth) - (no pink color) +(faint pink color) ++(obvious pink color) +++ (dark pink color)

Images taken after 3 days at 37 ºC on 7H11 agar

Gene *6*; Score *3*

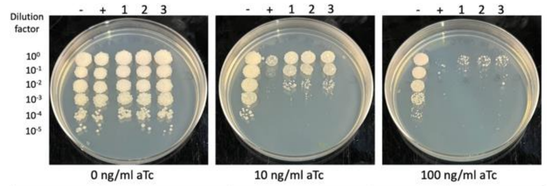


| Lane | Plasmid name | Gene name, replicate | Toxic/Non-toxic | Colony color on 100 ng/ml aTc plate* |
|---|---|---|---|---|
| - Non-toxic control | pExTra03 | Fruitloop *52* mutant | Non-toxic | + |
| + Toxic control | pExTra02 | Fruitloop *52* | Toxic | NG |
| 1 | pExTra-NormanBulbieJr6 | NormanBulbieJr *6* replicate 1 | Toxic | - |
| 2 | pExTra-NormanBulbieJr6 | NormanBulbieJr *6* replicate 2 | Toxic | - |
| 3 | pExTra-NormanBulbieJr6 | NormanBulbieJr *6* replicate 3 | Toxic | - |

*Key: NG (no growth) - (no pink color) +(faint pink color) ++(obvious pink color) +++ (dark pink color)

Images taken after 4 days at 37 ºC on 7H11 agar

Gene *7*; Score *0*

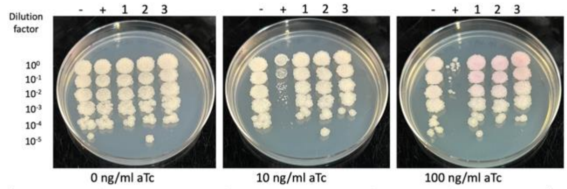


| Lane | Plasmid name | Gene name, replicate | Toxic/Non-toxic | Colony color on 100 ng/ml aTc plate* |
|---|---|---|---|---|
| - Non-toxic control | pExTra03 | Fruitloop *52* mutant | Non-toxic | + |
| + Toxic control | pExTra02 | Fruitloop *52* | Toxic | - |
| 1 | pExTra-NormanBulbieJr7 | NormanBulbieJr *7* replicate 1 | Non-toxic | + |
| 2 | pExTra-NormanBulbieJr7 | NormanBulbieJr *7* replicate 2 | Non-toxic | + |
| 3 | pExTra-NormanBulbieJr7 | NormanBulbieJr *7* replicate 3 | Non-toxic | + |

*Key: NG (no growth) - (no pink color) +(faint pink color) ++(obvious pink color) +++ (dark pink color)

Images taken after 3 days at 37 ºC on 7H11 agar

Gene *8*; Score *0*

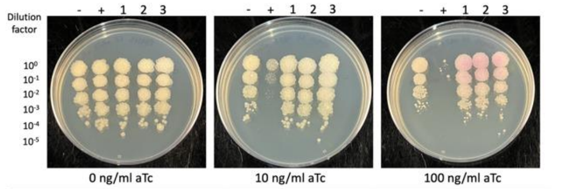


| Lane | Plasmid name | Gene name, replicate | Toxic/Non-toxic | Colony color on 100 ng/ml aTc plate* |
|---|---|---|---|---|
| - Non-toxic control | pExTra03 | Fruitloop *52* mutant | Non-toxic | - |
| + Toxic control | pExTra02 | Fruitloop *52* | Toxic | NG |
| 1 | pExTra-NormanBulbieJr8 | NormanBulbieJr *8* replicate 1 | Non-toxic | ++ |
| 2 | pExTra-NormanBulbieJr8 | NormanBulbieJr *8* replicate 2 | Non-toxic | ++ |
| 3 | pExTra-NormanBulbieJr8 | NormanBulbieJr *8* replicate 3 | Non-toxic | ++ |

*Key: NG (no growth) - (no pink color) +(faint pink color) ++(obvious pink color) +++ (dark pink color)

Figure S1

Gene *9*; Score *0*

Images taken after 3 days at 37 ºC on 7H11 agar

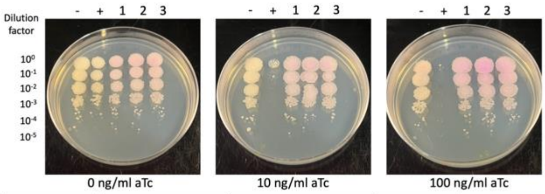


| Lane | Plasmid name | Gene name, replicate | Toxic/Non-toxic | Colony color on 100 ng/ml aTc plate* |
|---|---|---|---|---|
| - Non-toxic control | pExTra03 | Fruitloop *52* mutant | Non-toxic | + |
| + Toxic control | pExTra02 | Fruitloop *52* | Toxic | NG |
| 1 | pExTra-NormanBulbieJr9 | NormanBulbieJr *9* replicate 1 | Non-toxic | ++ |
| 2 | pExTra-NormanBulbieJr9 | NormanBulbieJr *9* replicate 2 | Non-toxic | ++ |
| 3 | pExTra-NormanBulbieJr9 | NormanBulbieJr *9* replicate 3 | Non-toxic | ++ |

*Key: NG (no growth) - (no pink color) +(faint pink color) ++(obvious pink color) +++ (dark pink color)

Gene *10*; Score *0*

Images taken after 3 days at 37 ºC on 7H11 agar

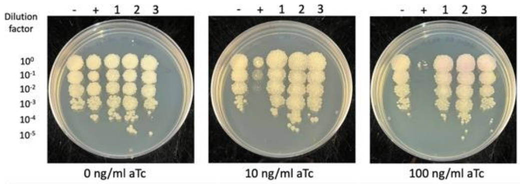


| Lane | Plasmid name | Gene name, replicate | Toxic/Non-toxic | Colony color on 100 ng/ml aTc plate* |
|---|---|---|---|---|
| - Non-toxic control | pExTra03 | Fruitloop *52* mutant | Non-toxic | - |
| + Toxic control | pExTra02 | Fruitloop *52* | Toxic | NG |
| 1 | pExTra-NormanBulbieJr10 | NormanBulbieJr *10* replicate 1 | Non-toxic | + |
| 2 | pExTra-NormanBulbieJr10 | NormanBulbieJr *10* replicate 2 | Non-toxic | + |
| 3 | pExTra-NormanBulbieJr10 | NormanBulbieJr *10* replicate 3 | Non-toxic | + |

*Key: NG (no growth) - (no pink color) +(faint pink color) ++(obvious pink color) +++ (dark pink color)

Gene *11*; Score *0*

Images taken after 4 days at 37 ºC on 7H11 agar

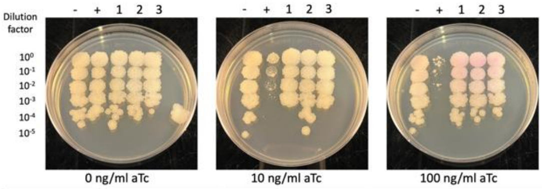


| Lane | Plasmid name | Gene name, replicate | Toxic/Non-toxic | Colony color on 100 ng/ml aTc plate* |
|---|---|---|---|---|
| - Non-toxic control | pExTra03 | Fruitloop *52* mutant | Non-toxic | + |
| + Toxic control | pExTra02 | Fruitloop *52* | Toxic | - |
| 1 | pExTra-NormanBulbieJr11 | NormanBulbieJr *11* replicate 1 | Non-toxic | + |
| 2 | pExTra-NormanBulbieJr11 | NormanBulbieJr *11* replicate 2 | Non-toxic | + |
| 3 | pExTra-NormanBulbieJr11 | NormanBulbieJr *11* replicate 3 | Non-toxic | + |

*Key: NG (no growth) - (no pink color) +(faint pink color) ++(obvious pink color) +++ (dark pink color)

Gene *12*; Score *0*

Images taken after 3 days at 37 ºC on 7H11 agar

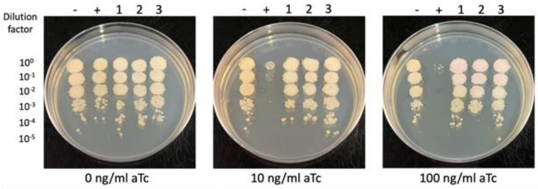


| Lane | Plasmid name | Gene name, replicate | Toxic/Non-toxic | Colony color on 100 ng/ml aTc plate* |
|---|---|---|---|---|
| - Non-toxic control | pExTra03 | Fruitloop *52* mutant | Non-toxic | - |
| + Toxic control | pExTra02 | Fruitloop *52* | Toxic | NG |
| 1 | pExTra-NormanBulbieJr12 | NormanBulbieJr *12* replicate 1 | Non-toxic | + |
| 2 | pExTra-NormanBulbieJr12 | NormanBulbieJr *12* replicate 2 | Non-toxic | + |
| 3 | pExTra-NormanBulbieJr12 | NormanBulbieJr *12* replicate 3 | Non-toxic | + |

*Key: NG (no growth) - (no pink color) +(faint pink color) ++(obvious pink color) +++ (dark pink color)

Gene *13*; Score *0*

Images taken after 4 days at 37 ºC on 7H11 agar

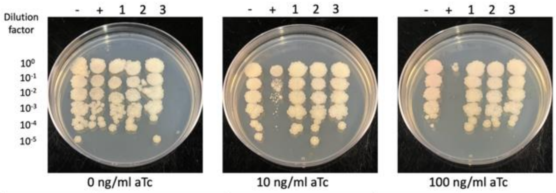


| Lane | Plasmid name | Gene name, replicate | Toxic/Non-toxic | Colony color on 100 ng/ml aTc plate* |
|---|---|---|---|---|
| - Non-toxic control | pExTra03 | Fruitloop *52* mutant | Non-toxic | + |
| + Toxic control | pExTra02 | Fruitloop *52* | Toxic | - |
| 1 | pExTra-NormanBulbieJr13 | NormanBulbieJr *13* replicate 1 | Non-toxic | - |
| 2 | pExTra-NormanBulbieJr13 | NormanBulbieJr *13* replicate 2 | Non-toxic | - |
| 3 | pExTra-NormanBulbieJr13 | NormanBulbieJr *13* replicate 3 | Non-toxic | - |

*Key: NG (no growth) - (no pink color) +(faint pink color) ++(obvious pink color) +++ (dark pink color)

Gene *14*; Score *0*

Images taken after 4 days at 37 ºC on 7H11 agar

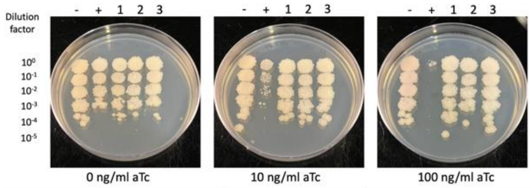


| Lane | Plasmid name | Gene name, replicate | Toxic/Non-toxic | Colony color on 100 ng/ml aTc plate* |
|---|---|---|---|---|
| - Non-toxic control | pExTra03 | Fruitloop *52* mutant | Non-toxic | + |
| + Toxic control | pExTra02 | Fruitloop *52* | Toxic | - |
| 1 | pExTra-NormanBulbieJr14 | NormanBulbieJr *14* replicate 1 | Non-toxic | - |
| 2 | pExTra-NormanBulbieJr14 | NormanBulbieJr *14* replicate 2 | Non-toxic | - |
| 3 | pExTra-NormanBulbieJr14 | NormanBulbieJr *14* replicate 3 | Non-toxic | - |

*Key: NG (no growth) - (no pink color) +(faint pink color) ++(obvious pink color) +++ (dark pink color)

Gene *15*; Score *0*

Images taken after 4 days at 37 ºC on 7H11 agar

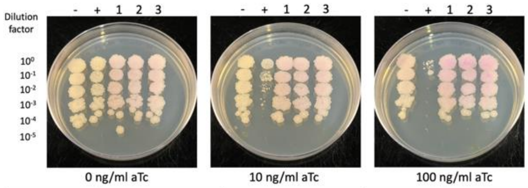


| Lane | Plasmid name | Gene name, replicate | Toxic/Non-toxic | Colony color on 100 ng/ml aTc plate* |
|---|---|---|---|---|
| - Non-toxic control | pExTra03 | Fruitloop *52* mutant | Non-toxic | + |
| + Toxic control | pExTra02 | Fruitloop *52* | Toxic | - |
| 1 | pExTra-NormanBulbieJr15 | NormanBulbieJr *15* replicate 1 | Non-toxic | ++ |
| 2 | pExTra-NormanBulbieJr15 | NormanBulbieJr *15* replicate 2 | Non-toxic | ++ |
| 3 | pExTra-NormanBulbieJr15 | NormanBulbieJr *15* replicate 3 | Non-toxic | ++ |

*Key: NG (no growth) - (no pink color) +(faint pink color) ++(obvious pink color) +++ (dark pink color)

Gene *16*; Score *0*

Images taken after 3 days at 37 ºC on 7H11 agar

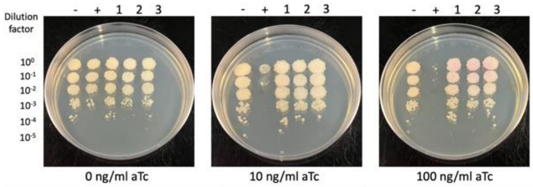


| Lane | Plasmid name | Gene name, replicate | Toxic/Non-toxic | Colony color on 100 ng/ml aTc plate* |
|---|---|---|---|---|
| - Non-toxic control | pExTra03 | Fruitloop *52* mutant | Non-toxic | - |
| + Toxic control | pExTra02 | Fruitloop *52* | Toxic | NG |
| 1 | pExTra-NormanBulbieJr16 | NormanBulbieJr *16* replicate 1 | Non-toxic | + |
| 2 | pExTra-NormanBulbieJr16 | NormanBulbieJr *16* replicate 2 | Non-toxic | + |
| 3 | pExTra-NormanBulbieJr16 | NormanBulbieJr *16* replicate 3 | Non-toxic | + |

*Key: NG (no growth) - (no pink color) +(faint pink color) ++(obvious pink color) +++ (dark pink color)

### Figure S1

Gene *17*; Score *0*

Images taken after 4 days at 37 ºC on 7H11 agar

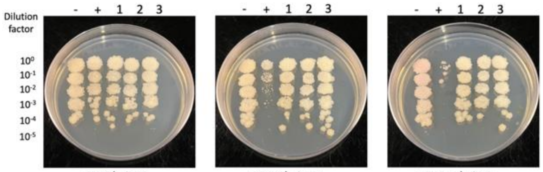


0 ng/ml aTc 10 ng/ml aTc 100 ng/ml aTc

| Lane | Plasmid name | Gene name, replicate | Toxic/Non-toxic | Colony color on 100 ng/ml aTc plate* |
|---|---|---|---|---|
| - Non-toxic control | pExTra03 | Fruitloop *52* mutant | Non-toxic | + |
| + Toxic control | pExTra02 | Fruitloop *52* | Toxic | - |
| 1 | pExTra-NormanBulbieJr17 | NormanBulbieJr *17* replicate 1 | Non-toxic | - |
| 2 | pExTra-NormanBulbieJr17 | NormanBulbieJr *17* replicate 2 | Non-toxic | - |
| 3 | pExTra-NormanBulbieJr17 | NormanBulbieJr *17* replicate 3 | Non-toxic | - |

*Key: NG (no growth) - (no pink color) +(faint pink color) ++(obvious pink color) +++ (dark pink color)

Gene *18*; Score *2*

Images taken after 4 days at 37 ºC on 7H11 agar

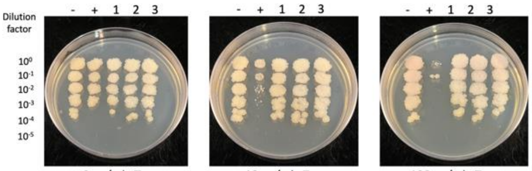


0 ng/ml aTc 10 ng/ml aTc 100 ng/ml aTc

| Lane | Plasmid name | Gene name, replicate | Toxic/Non-toxic | Colony color on 100 ng/ml aTc plate* |
|---|---|---|---|---|
| - Non-toxic control | pExTra03 | Fruitloop *52* mutant | Non-toxic | + |
| + Toxic control | pExTra02 | Fruitloop *52* | Toxic | NG |
| 1 | pExTra-NormanBulbieJr18 | NormanBulbieJr *18* replicate 1 | Non-toxic | + |
| 2 | pExTra-NormanBulbieJr18 | NormanBulbieJr *18* replicate 2 | Non-toxic | + |
| 3 | pExTra-NormanBulbieJr18 | NormanBulbieJr *18* replicate 3 | Non-toxic | + |

*Key: NG (no growth) - (no pink color) +(faint pink color) ++(obvious pink color) +++ (dark pink color)

Gene *19*; Score *1*

Images taken after 4 days at 37 ºC on 7H11 agar

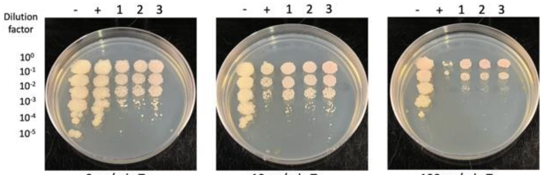


0 ng/ml aTc 10 ng/ml aTc 100 ng/ml aTc

| Lane | Plasmid name | Gene name, replicate | Toxic/Non-toxic | Colony color on 100 ng/ml aTc plate* |
|---|---|---|---|---|
| - Non-toxic control | pExTra03 | Fruitloop *52* mutant | Non-toxic | + |
| + Toxic control | pExTra02 | Fruitloop *52* | Toxic | - |
| 1 | pExTra-NormanBulbieJr19 | NormanBulbieJr *19* replicate 1 | Toxic | + |
| 2 | pExTra-NormanBulbieJr19 | NormanBulbieJr *19* replicate 2 | Toxic | + |
| 3 | pExTra-NormanBulbieJr19 | NormanBulbieJr *19* replicate 3 | Toxic | + |

*Key: NG (no growth) - (no pink color) +(faint pink color) ++(obvious pink color) +++ (dark pink color)

Gene *20*; Score *0*

Images taken after 3 days at 37 ºC on 7H11 agar

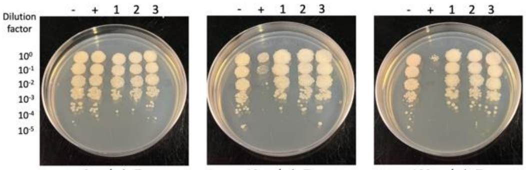


0 ng/ml aTc 10 ng/ml aTc 100 ng/ml aTc

| Lane | Plasmid name | Gene name, replicate | Toxic/Non-toxic | Colony color on 100 ng/ml aTc plate* |
|---|---|---|---|---|
| - Non-toxic control | pExTra03 | Fruitloop *52* mutant | Non-toxic | + |
| + Toxic control | pExTra02 | Fruitloop *52* | Toxic | - |
| 1 | pExTra-NormanBulbieJr20 | NormanBulbieJr *20* replicate 1 | Non-toxic | - |
| 2 | pExTra-NormanBulbieJr20 | NormanBulbieJr *20* replicate 2 | Non-toxic | - |
| 3 | pExTra-NormanBulbieJr20 | NormanBulbieJr *20* replicate 3 | Non-toxic | - |

*Key: NG (no growth) - (no pink color) +(faint pink color) ++(obvious pink color) +++ (dark pink color)

Gene *21*; Score *0*

Images taken after 3 days at 37 ºC on 7H11 agar

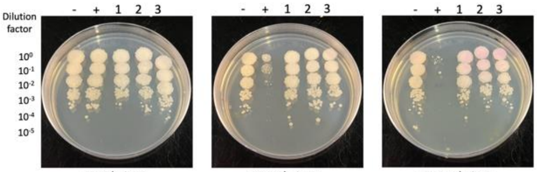


0 ng/ml aTc 10 ng/ml aTc 100 ng/ml aTc

| Lane | Plasmid name | Gene name, replicate | Toxic/Non-toxic | Colony color on 100 ng/ml aTc plate* |
|---|---|---|---|---|
| - Non-toxic control | pExTra03 | Fruitloop *52* mutant | Non-toxic | + |
| + Toxic control | pExTra02 | Fruitloop *52* | Toxic | - |
| 1 | pExTra-NormanBulbieJr21 | NormanBulbieJr *21* replicate 1 | Non-toxic | + |
| 2 | pExTra-NormanBulbieJr21 | NormanBulbieJr *21* replicate 2 | Non-toxic | + |
| 3 | pExTra-NormanBulbieJr21 | NormanBulbieJr *21* replicate 3 | Non-toxic | + |

*Key: NG (no growth) - (no pink color) +(faint pink color) ++(obvious pink color) +++ (dark pink color)

Gene *22*; Score *0*

Images taken after 4 days at 37 ºC on 7H11 agar

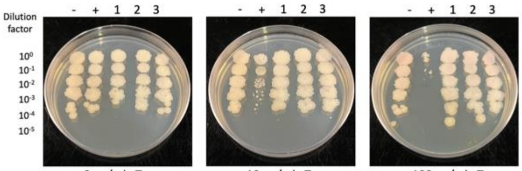


0 ng/ml aTc 10 ng/ml aTc 100 ng/ml aTc

| Lane | Plasmid name | Gene name, replicate | Toxic/Non-toxic | Colony color on 100 ng/ml aTc plate* |
|---|---|---|---|---|
| - Non-toxic control | pExTra03 | Fruitloop *52* mutant | Non-toxic | + |
| + Toxic control | pExTra02 | Fruitloop *52* | Toxic | - |
| 1 | pExTra-NormanBulbieJr22 | NormanBulbieJr *22* replicate 1 | Non-toxic | + |
| 2 | pExTra-NormanBulbieJr22 | NormanBulbieJr *22* replicate 2 | Non-toxic | + |
| 3 | pExTra-NormanBulbieJr22 | NormanBulbieJr *22* replicate 3 | Non-toxic | + |

*Key: NG (no growth) - (no pink color) +(faint pink color) ++(obvious pink color) +++ (dark pink color)

Gene *23*; Score *0*

Images taken after 3 days at 37 ºC on 7H11 agar

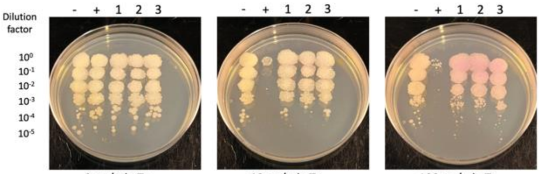


0 ng/ml aTc 10 ng/ml aTc 100 ng/ml aTc

| Lane | Plasmid name | Gene name, replicate | Toxic/Non-toxic | Colony color on 100 ng/ml aTc plate* |
|---|---|---|---|---|
| - Non-toxic control | pExTra03 | Fruitloop *52* mutant | Non-toxic | + |
| + Toxic control | pExTra02 | Fruitloop *52* | Toxic | - |
| 1 | pExTra-NormanBulbieJr23 | NormanBulbieJr *23* replicate 1 | Non-toxic | ++ |
| 2 | pExTra-NormanBulbieJr23 | NormanBulbieJr *23* replicate 2 | Non-toxic | ++ |
| 3 | pExTra-NormanBulbieJr23 | NormanBulbieJr *23* replicate 3 | Non-toxic | ++ |

*Key: NG (no growth) - (no pink color) +(faint pink color) ++(obvious pink color) +++ (dark pink color)

Gene *24*; Score *0*

Images taken after 4 days at 37 ºC on 7H11 agar

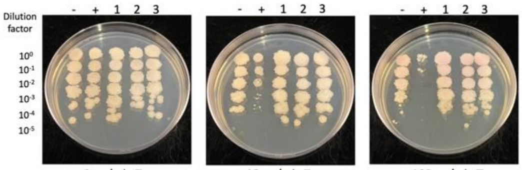


0 ng/ml aTc 10 ng/ml aTc 100 ng/ml aTc

| Lane | Plasmid name | Gene name, replicate | Toxic/Non-toxic | Colony color on 100 ng/ml aTc plate* |
|---|---|---|---|---|
| - Non-toxic control | pExTra03 | Fruitloop *52* mutant | Non-toxic | + |
| + Toxic control | pExTra02 | Fruitloop *52* | Toxic | - |
| 1 | pExTra-NormanBulbieJr24 | NormanBulbieJr *24* replicate 1 | Non-toxic | + |
| 2 | pExTra-NormanBulbieJr24 | NormanBulbieJr *24* replicate 2 | Non-toxic | + |
| 3 | pExTra-NormanBulbieJr24 | NormanBulbieJr *24* replicate 3 | Non-toxic | + |

*Key: NG (no growth) - (no pink color) +(faint pink color) ++(obvious pink color) +++ (dark pink color)

Figure S1

Images taken after 4 days at 37 ºC on 7H11 agar

Gene *25*; Score *1*

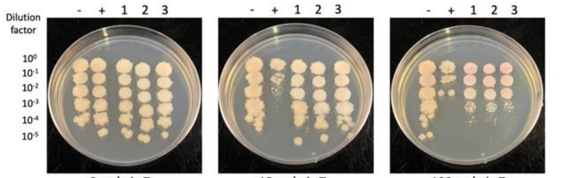


0 ng/ml aTc | 10 ng/ml aTc | 100 ng/ml aTc

| Lane | Plasmid name | Gene name, replicate | Toxic/Non-toxic | Colony color on 100 ng/ml aTc plate* |
|---|---|---|---|---|
| - Non-toxic control | pExTra03 | Fruitloop *52* mutant | Non-toxic | + |
| + Toxic control | pExTra02 | Fruitloop *52* | Toxic | - |
| 1 | pExTra-NormanBulbieJr25 | NormanBulbieJr *25* replicate 1 | Toxic | + |
| 2 | pExTra-NormanBulbieJr25 | NormanBulbieJr *25* replicate 2 | Toxic | + |
| 3 | pExTra-NormanBulbieJr25 | NormanBulbieJr *25* replicate 3 | Toxic | + |

*Key: NG (no growth) - (no pink color) +(faint pink color) ++(obvious pink color) +++ (dark pink color)

Images taken after 5 days at 37 ºC on 7H11 agar

Gene *26*; Score *0*

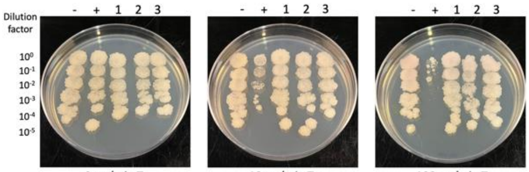


0 ng/ml aTc | 10 ng/ml aTc | 100 ng/ml aTc

| Lane | Plasmid name | Gene name, replicate | Toxic/Non-toxic | Colony color on 100 ng/ml aTc plate* |
|---|---|---|---|---|
| - Non-toxic control | pExTra03 | Fruitloop *52* mutant | Non-toxic | + |
| + Toxic control | pExTra02 | Fruitloop *52* | Toxic | - |
| 1 | pExTra-NormanBulbieJr26 | NormanBulbieJr *26* replicate 1 | Non-toxic | - |
| 2 | pExTra-NormanBulbieJr26 | NormanBulbieJr *26* replicate 2 | Non-toxic | - |
| 3 | pExTra-NormanBulbieJr26 | NormanBulbieJr *26* replicate 3 | Non-toxic | - |

*Key: NG (no growth) - (no pink color) +(faint pink color) ++(obvious pink color) +++ (dark pink color)

Images taken after 4 days at 37 ºC on 7H11 agar

Gene *27*; Score *0*

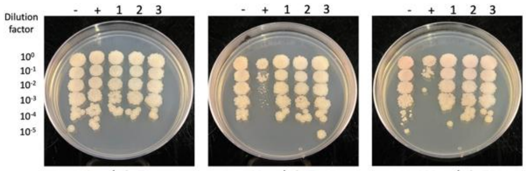


0 ng/ml aTc | 10 ng/ml aTc | 100 ng/ml aTc

| Lane | Plasmid name | Gene name, replicate | Toxic/Non-toxic | Colony color on 100 ng/ml aTc plate* |
|---|---|---|---|---|
| - Non-toxic control | pExTra03 | Fruitloop *52* mutant | Non-toxic | + |
| + Toxic control | pExTra02 | Fruitloop *52* | Toxic | - |
| 1 | pExTra-NormanBulbieJr27 | NormanBulbieJr *27* replicate 1 | Non-toxic | + |
| 2 | pExTra-NormanBulbieJr27 | NormanBulbieJr *27* replicate 2 | Non-toxic | + |
| 3 | pExTra-NormanBulbieJr27 | NormanBulbieJr *27* replicate 3 | Non-toxic | + |

*Key: NG (no growth) - (no pink color) +(faint pink color) ++(obvious pink color) +++ (dark pink color)

Images taken after 3 days at 37 ºC on 7H11 agar

Gene *28*; Score *0*

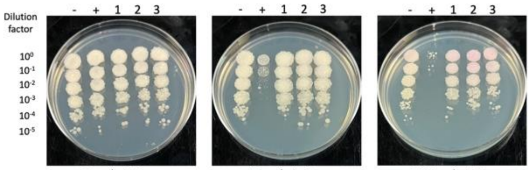


0 ng/ml aTc | 10 ng/ml aTc | 100 ng/ml aTc

| Lane | Plasmid name | Gene name, replicate | Toxic/Non-toxic | Colony color on 100 ng/ml aTc plate* |
|---|---|---|---|---|
| - Non-toxic control | pExTra03 | Fruitloop *52* mutant | Non-toxic | + |
| + Toxic control | pExTra02 | Fruitloop *52* | Toxic | - |
| 1 | pExTra-NormanBulbieJr28 | NormanBulbieJr *28* replicate 1 | Non-toxic | + |
| 2 | pExTra-NormanBulbieJr28 | NormanBulbieJr *28* replicate 2 | Non-toxic | + |
| 3 | pExTra-NormanBulbieJr28 | NormanBulbieJr *28* replicate 3 | Non-toxic | + |

*Key: NG (no growth) - (no pink color) +(faint pink color) ++(obvious pink color) +++ (dark pink color)

Images taken after 4 days at 37 ºC on 7H11 agar

Gene *29*; Score *0*

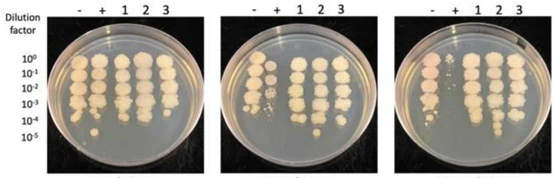


0 ng/ml aTc | 10 ng/ml aTc | 100 ng/ml aTc

| Lane | Plasmid name | Gene name, replicate | Toxic/Non-toxic | Colony color on 100 ng/ml aTc plate* |
|---|---|---|---|---|
| - Non-toxic control | pExTra03 | Fruitloop *52* mutant | Non-toxic | + |
| + Toxic control | pExTra02 | Fruitloop *52* | Toxic | - |
| 1 | pExTra-NormanBulbieJr29 | NormanBulbieJr *29* replicate 1 | Non-toxic | - |
| 2 | pExTra-NormanBulbieJr29 | NormanBulbieJr *29* replicate 2 | Non-toxic | - |
| 3 | pExTra-NormanBulbieJr29 | NormanBulbieJr *29* replicate 3 | Non-toxic | - |

*Key: NG (no growth) - (no pink color) +(faint pink color) ++(obvious pink color) +++ (dark pink color)

Images taken after 4 days at 37 ºC on 7H11 agar

Gene *30*; Score *2*

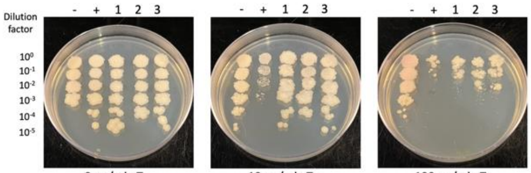


0 ng/ml aTc | 10 ng/ml aTc | 100 ng/ml aTc

| Lane | Plasmid name | Gene name, replicate | Toxic/Non-toxic | Colony color on 100 ng/ml aTc plate* |
|---|---|---|---|---|
| - Non-toxic control | pExTra03 | Fruitloop *52* mutant | Non-toxic | + |
| + Toxic control | pExTra02 | Fruitloop *52* | Toxic | - |
| 1 | pExTra-NormanBulbieJr30 | NormanBulbieJr *30* replicate 1 | Toxic | - |
| 2 | pExTra-NormanBulbieJr30 | NormanBulbieJr *30* replicate 2 | Toxic | - |
| 3 | pExTra-NormanBulbieJr30 | NormanBulbieJr *30* replicate 3 | Toxic | - |

*Key: NG (no growth) - (no pink color) +(faint pink color) ++(obvious pink color) +++ (dark pink color)

Images taken after 4 days at 37 ºC on 7H11 agar

Gene *31*; Score *0*

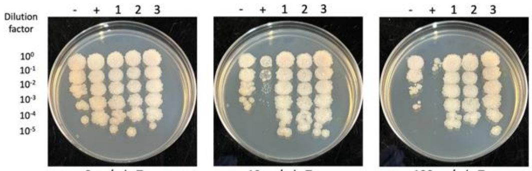


0 ng/ml aTc | 10 ng/ml aTc | 100 ng/ml aTc

| Lane | Plasmid name | Gene name, replicate | Toxic/Non-toxic | Colony color on 100 ng/ml aTc plate* |
|---|---|---|---|---|
| - Non-toxic control | pExTra03 | Fruitloop *52* mutant | Non-toxic | + |
| + Toxic control | pExTra02 | Fruitloop *52* | Toxic | - |
| 1 | pExTra-NormanBulbieJr31 | NormanBulbieJr *31* replicate 1 | Non-toxic | - |
| 2 | pExTra-NormanBulbieJr31 | NormanBulbieJr *31* replicate 2 | Non-toxic | - |
| 3 | pExTra-NormanBulbieJr31 | NormanBulbieJr *31* replicate 3 | Non-toxic | - |

*Key: NG (no growth) - (no pink color) +(faint pink color) ++(obvious pink color) +++ (dark pink color)

Images taken after 4 days at 37 ºC on 7H11 agar

Gene *32*; Score *0*

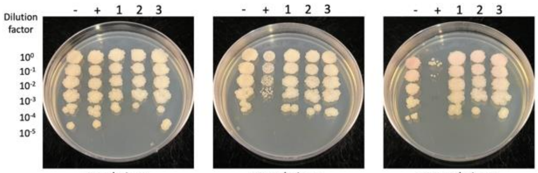


0 ng/ml aTc | 10 ng/ml aTc | 100 ng/ml aTc

| Lane | Plasmid name | Gene name, replicate | Toxic/Non-toxic | Colony color on 100 ng/ml aTc plate* |
|---|---|---|---|---|
| - Non-toxic control | pExTra03 | Fruitloop *52* mutant | Non-toxic | + |
| + Toxic control | pExTra02 | Fruitloop *52* | Toxic | - |
| 1 | pExTra-NormanBulbieJr32 | NormanBulbieJr *32* replicate 1 | Non-toxic | + |
| 2 | pExTra-NormanBulbieJr32 | NormanBulbieJr *32* replicate 2 | Non-toxic | + |
| 3 | pExTra-NormanBulbieJr32 | NormanBulbieJr *32* replicate 3 | Non-toxic | + |

*Key: NG (no growth) - (no pink color) +(faint pink color) ++(obvious pink color) +++ (dark pink color)

### Figure S1

Images taken after 3 days at 37 ºC on 7H11 agar

#### Gene *33*; Score *3*

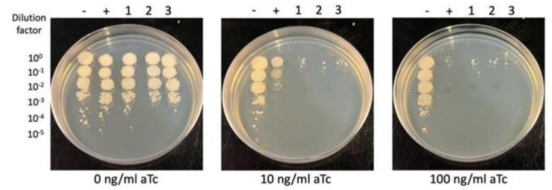


| Lane | Plasmid name | Gene name, replicate | Toxic/Non-toxic | Colony color on 100 ng/ml aTc plate* |
|---|---|---|---|---|
| - Non-toxic control | pExTra03 | Fruitloop *52* mutant | Non-toxic | + |
| + Toxic control | pExTra02 | Fruitloop *52* | Toxic | NG |
| 1 | pExTra-NormanBulbieJr33 | NormanBulbieJr *33* replicate 1 | Toxic | - |
| 2 | pExTra-NormanBulbieJr33 | NormanBulbieJr *33* replicate 2 | Toxic | - |
| 3 | pExTra-NormanBulbieJr33 | NormanBulbieJr *33* replicate 3 | Toxic | - |

*Key: NG (no growth) - (no pink color) +(faint pink color) ++(obvious pink color) +++ (dark pink color)

Images taken after 4 days at 37 ºC on 7H11 agar

#### Gene *34*; Score *2*

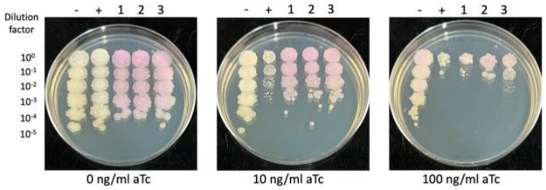


| Lane | Plasmid name | Gene name, replicate | Toxic/Non-toxic | Colony color on 100 ng/ml aTc plate* |
|---|---|---|---|---|
| - Non-toxic control | pExTra03 | Fruitloop *52* mutant | Non-toxic | + |
| + Toxic control | pExTra02 | Fruitloop *52* | Toxic | - |
| 1 | pExTra-NormanBulbieJr34 | NormanBulbieJr *34* replicate 1 | Toxic | + |
| 2 | pExTra-NormanBulbieJr34 | NormanBulbieJr *34* replicate 2 | Toxic | + |
| 3 | pExTra-NormanBulbieJr34 | NormanBulbieJr *34* replicate 3 | Toxic | + |

*Key: NG (no growth) - (no pink color) +(faint pink color) ++(obvious pink color) +++ (dark pink color)

Images taken after 4 days at 37 ºC on 7H11 agar

#### Gene *35*; Score *2*

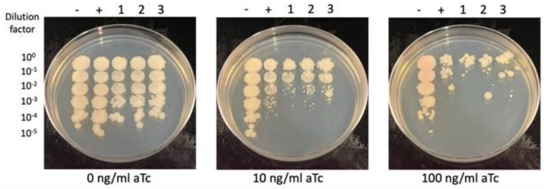


| Lane | Plasmid name | Gene name, replicate | Toxic/Non-toxic | Colony color on 100 ng/ml aTc plate* |
|---|---|---|---|---|
| - Non-toxic control | pExTra03 | Fruitloop *52* mutant | Non-toxic | + |
| + Toxic control | pExTra02 | Fruitloop *52* | Toxic | - |
| 1 | pExTra-NormanBulbieJr35 | NormanBulbieJr *35* replicate 1 | Toxic | - |
| 2 | pExTra-NormanBulbieJr35 | NormanBulbieJr *35* replicate 2 | Toxic | - |
| 3 | pExTra-NormanBulbieJr35 | NormanBulbieJr *35* replicate 3 | Toxic | - |

*Key: NG (no growth) - (no pink color) +(faint pink color) ++(obvious pink color) +++ (dark pink color)

Images taken after 3 days at 37 ºC on 7H11 agar

#### Gene *36*; Score *0*

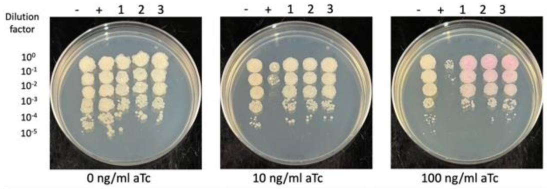


| Lane | Plasmid name | Gene name, replicate | Toxic/Non-toxic | Colony color on 100 ng/ml aTc plate* |
|---|---|---|---|---|
| - Non-toxic control | pExTra03 | Fruitloop *52* mutant | Non-toxic | + |
| + Toxic control | pExTra02 | Fruitloop *52* | Toxic | - |
| 1 | pExTra-NormanBulbieJr36 | NormanBulbieJr *36* replicate 1 | Non-toxic | ++ |
| 2 | pExTra-NormanBulbieJr36 | NormanBulbieJr *36* replicate 2 | Non-toxic | ++ |
| 3 | pExTra-NormanBulbieJr36 | NormanBulbieJr *36* replicate 3 | Non-toxic | ++ |

*Key: NG (no growth) - (no pink color) +(faint pink color) ++(obvious pink color) +++ (dark pink color)

Images taken after 3 days at 37 ºC on 7H11 agar

#### Gene *37*; Score *3*

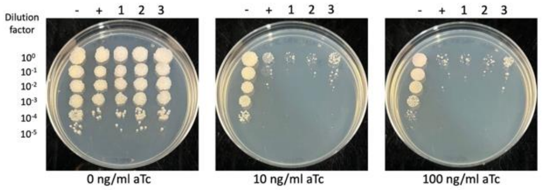


| Lane | Plasmid name | Gene name, replicate | Toxic/Non-toxic | Colony color on 100 ng/ml aTc plate* |
|---|---|---|---|---|
| - Non-toxic control | pExTra03 | Fruitloop *52* mutant | Non-toxic | + |
| + Toxic control | pExTra02 | Fruitloop *52* | Toxic | - |
| 1 | pExTra-NormanBulbieJr37 | NormanBulbieJr *37* replicate 1 | Toxic | - |
| 2 | pExTra-NormanBulbieJr37 | NormanBulbieJr *37* replicate 2 | Toxic | - |
| 3 | pExTra-NormanBulbieJr37 | NormanBulbieJr *37* replicate 3 | Toxic | - |

*Key: NG (no growth) - (no pink color) +(faint pink color) ++(obvious pink color) +++ (dark pink color)

Images taken after 4 days at 37 ºC on 7H11 agar

#### Gene *38*; Score *0*

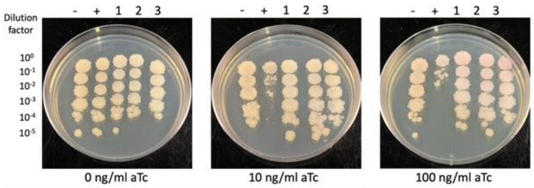


| Lane | Plasmid name | Gene name, replicate | Toxic/Non-toxic | Colony color on 100 ng/ml aTc plate* |
|---|---|---|---|---|
| - Non-toxic control | pExTra03 | Fruitloop *52* mutant | Non-toxic | + |
| + Toxic control | pExTra02 | Fruitloop *52* | Toxic | - |
| 1 | pExTra-NormanBulbieJr38 | NormanBulbieJr *38* replicate 1 | Non-toxic | + |
| 2 | pExTra-NormanBulbieJr38 | NormanBulbieJr *38* replicate 2 | Non-toxic | + |
| 3 | pExTra-NormanBulbieJr38 | NormanBulbieJr *38* replicate 3 | Non-toxic | + |

*Key: NG (no growth) - (no pink color) +(faint pink color) ++(obvious pink color) +++ (dark pink color)

Images taken after 3 days at 37 ºC on 7H11 agar

#### Gene *39*; Score *1*

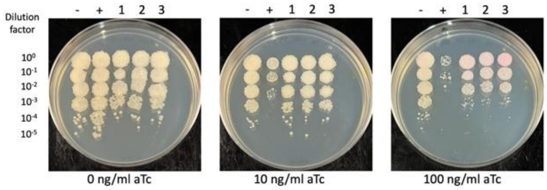


| Lane | Plasmid name | Gene name, replicate | Toxic/Non-toxic | Colony color on 100 ng/ml aTc plate* |
|---|---|---|---|---|
| - Non-toxic control | pExTra03 | Fruitloop *52* mutant | Non-toxic | + |
| + Toxic control | pExTra02 | Fruitloop *52* | Toxic | - |
| 1 | pExTra-NormanBulbieJr39 | NormanBulbieJr *39* replicate 1 | Toxic | + |
| 2 | pExTra-NormanBulbieJr39 | NormanBulbieJr *39* replicate 2 | Toxic | + |
| 3 | pExTra-NormanBulbieJr39 | NormanBulbieJr *39* replicate 3 | Toxic | + |

*Key: NG (no growth) - (no pink color) +(faint pink color) ++(obvious pink color) +++ (dark pink color)

Images taken after 3 days at 37 ºC on 7H11 agar

#### Gene *40*; Score *0*

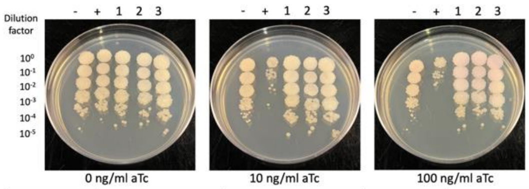


| Lane | Plasmid name | Gene name, replicate | Toxic/Non-toxic | Colony color on 100 ng/ml aTc plate* |
|---|---|---|---|---|
| - Non-toxic control | pExTra03 | Fruitloop *52* mutant | Non-toxic | + |
| + Toxic control | pExTra02 | Fruitloop *52* | Toxic | - |
| 1 | pExTra-NormanBulbieJr40 | NormanBulbieJr *40* replicate 1 | Non-toxic | + |
| 2 | pExTra-NormanBulbieJr40 | NormanBulbieJr *40* replicate 2 | Non-toxic | + |
| 3 | pExTra-NormanBulbieJr40 | NormanBulbieJr *40* replicate 3 | Non-toxic | + |

*Key: NG (no growth) - (no pink color) +(faint pink color) ++(obvious pink color) +++ (dark pink color)

Figure S1

##### *Gene 41; Score 0*

Images taken after 3 days at 37 ºC on 7H11 agar

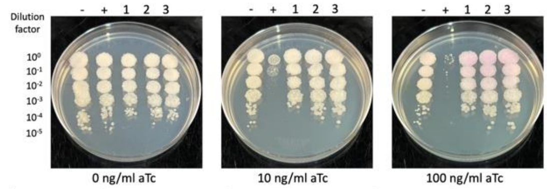


| Lane | Plasmid name | Gene name, replicate | Toxic/Non-toxic | Colony color on 100 ng/ml aTc plate* |
|---|---|---|---|---|
| - Non-toxic control | pExTra03 | Fruitloop *52* mutant | Non-toxic | + |
| + Toxic control | pExTra02 | Fruitloop *52* | Toxic | - |
| 1 | pExTra-NormanBulbieJr41 | NormanBulbieJr *41* replicate 1 | Non-toxic | ++ |
| 2 | pExTra-NormanBulbieJr41 | NormanBulbieJr *41* replicate 2 | Non-toxic | ++ |
| 3 | pExTra-NormanBulbieJr41 | NormanBulbieJr *41* replicate 3 | Non-toxic | ++ |

*Key: NG (no growth) - (no pink color) +(faint pink color) ++(obvious pink color) +++ (dark pink color)

##### Gene *45*; Score *1*

Images taken after 3 days at 37 ºC on 7H11 agar

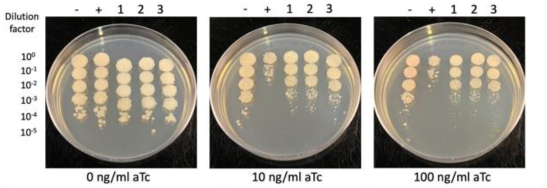


| Lane | Plasmid name | Gene name, replicate | Toxic/Non-toxic | Colony color on 100 ng/ml aTc plate* |
|---|---|---|---|---|
| - Non-toxic control | pExTra03 | Fruitloop *52* mutant | Non-toxic | + |
| + Toxic control | pExTra02 | Fruitloop *52* | Toxic | - |
| 1 | pExTra-NormanBulbieJr45 | NormanBulbieJr *45* replicate 1 | Toxic | - |
| 2 | pExTra-NormanBulbieJr45 | NormanBulbieJr *45* replicate 2 | Toxic | - |
| 3 | pExTra-NormanBulbieJr45 | NormanBulbieJr *45* replicate 3 | Toxic | - |

*Key: NG (no growth) - (no pink color) +(faint pink color) ++(obvious pink color) +++ (dark pink color)

##### *Gene 42; Score 0*

Images taken after 3 days at 37 ºC on 7H11 agar

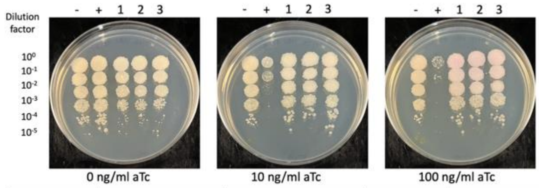


| Lane | Plasmid name | Gene name, replicate | Toxic/Non-toxic | Colony color on 100 ng/ml aTc plate* |
|---|---|---|---|---|
| - Non-toxic control | pExTra03 | Fruitloop *52* mutant | Non-toxic | + |
| + Toxic control | pExTra02 | Fruitloop *52* | Toxic | - |
| 1 | pExTra-NormanBulbieJr42 | NormanBulbieJr *42* replicate 1 | Non-toxic | + |
| 2 | pExTra-NormanBulbieJr42 | NormanBulbieJr *42* replicate 2 | Non-toxic | + |
| 3 | pExTra-NormanBulbieJr42 | NormanBulbieJr *42* replicate 3 | Non-toxic | + |

*Key: NG (no growth) - (no pink color) +(faint pink color) ++(obvious pink color) +++ (dark pink color)

##### Gene *46*; Score *0*

Images taken after 3 days at 37 ºC on 7H11 agar

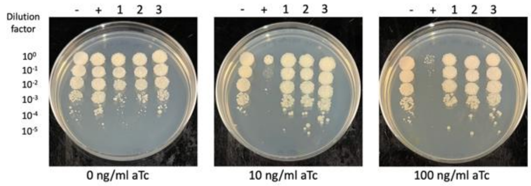


| Lane | Plasmid name | Gene name, replicate | Toxic/Non-toxic | Colony color on 100 ng/ml aTc plate* |
|---|---|---|---|---|
| - Non-toxic control | pExTra03 | Fruitloop *52* mutant | Non-toxic | + |
| + Toxic control | pExTra02 | Fruitloop *52* | Toxic | - |
| 1 | pExTra-NormanBulbieJr46 | NormanBulbieJr *46* replicate 1 | Non-toxic | + |
| 2 | pExTra-NormanBulbieJr46 | NormanBulbieJr *46* replicate 2 | Non-toxic | + |
| 3 | pExTra-NormanBulbieJr46 | NormanBulbieJr *46* replicate 3 | Non-toxic | + |

*Key: NG (no growth) - (no pink color) +(faint pink color) ++(obvious pink color) +++ (dark pink color)

##### *Gene 43; Score 3*

Images taken after 3 days at 37 ºC on 7H11 agar

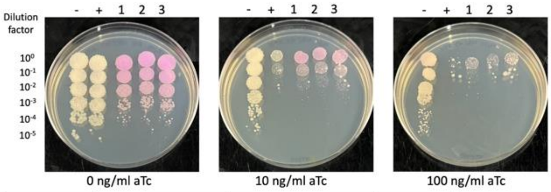


| Lane | Plasmid name | Gene name, replicate | Toxic/Non-toxic | Colony color on 100 ng/ml aTc plate* |
|---|---|---|---|---|
| - Non-toxic control | pExTra03 | Fruitloop *52* mutant | Non-toxic | + |
| + Toxic control | pExTra02 | Fruitloop *52* | Toxic | - |
| 1 | pExTra-NormanBulbieJr43 | NormanBulbieJr *43* replicate 1 | Toxic | + |
| 2 | pExTra-NormanBulbieJr43 | NormanBulbieJr *43* replicate 2 | Toxic | + |
| 3 | pExTra-NormanBulbieJr43 | NormanBulbieJr *43* replicate 3 | Toxic | + |

*Key: NG (no growth) - (no pink color) +(faint pink color) ++(obvious pink color) +++ (dark pink color)

##### Gene *47*; Score *3*

Images taken after 3 days at 37 ºC on 7H11 agar

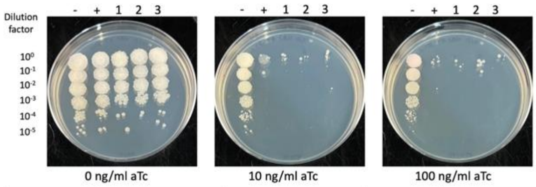


| Lane | Plasmid name | Gene name, replicate | Toxic/Non-toxic | Colony color on 100 ng/ml aTc plate* |
|---|---|---|---|---|
| - Non-toxic control | pExTra03 | Fruitloop *52* mutant | Non-toxic | + |
| + Toxic control | pExTra02 | Fruitloop *52* | Toxic | - |
| 1 | pExTra-NormanBulbieJr47 | NormanBulbieJr *47* replicate 1 | Toxic | - |
| 2 | pExTra-NormanBulbieJr47 | NormanBulbieJr *47* replicate 2 | Toxic | - |
| 3 | pExTra-NormanBulbieJr47 | NormanBulbieJr *47* replicate 3 | Toxic | - |

*Key: NG (no growth) - (no pink color) +(faint pink color) ++(obvious pink color) +++ (dark pink color)

##### *Gene 44; Score 3*

Images taken after 3 days at 37 ºC on 7H11 agar

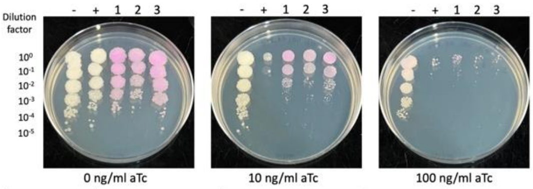


| Lane | Plasmid name | Gene name, replicate | Toxic/Non-toxic | Colony color on 100 ng/ml aTc plate* |
|---|---|---|---|---|
| - Non-toxic control | pExTra03 | Fruitloop *52* mutant | Non-toxic | + |
| + Toxic control | pExTra02 | Fruitloop *52* | Toxic | - |
| 1 | pExTra-NormanBulbieJr44 | NormanBulbieJr *44* replicate 1 | Toxic | + |
| 2 | pExTra-NormanBulbieJr44 | NormanBulbieJr *44* replicate 2 | Toxic | + |
| 3 | pExTra-NormanBulbieJr44 | NormanBulbieJr *44* replicate 3 | Toxic | + |

*Key: NG (no growth) - (no pink color) +(faint pink color) ++(obvious pink color) +++ (dark pink color)

##### Gene *48*; Score *0*

Images taken after 4 days at 37 ºC on 7H11 agar

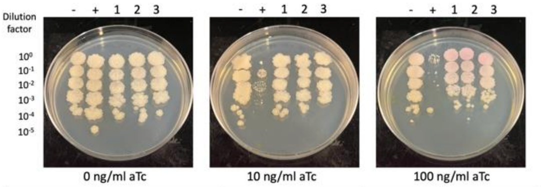


| Lane | Plasmid name | Gene name, replicate | Toxic/Non-toxic | Colony color on 100 ng/ml aTc plate* |
|---|---|---|---|---|
| - Non-toxic control | pExTra03 | Fruitloop *52* mutant | Non-toxic | + |
| + Toxic control | pExTra02 | Fruitloop *52* | Toxic | - |
| 1 | pExTra-NormanBulbieJr48 | NormanBulbieJr *48* replicate 1 | Non-toxic | ++ |
| 2 | pExTra-NormanBulbieJr48 | NormanBulbieJr *48* replicate 2 | Non-toxic | ++ |
| 3 | pExTra-NormanBulbieJr48 | NormanBulbieJr *48* replicate 3 | Non-toxic | ++ |

*Key: NG (no growth) - (no pink color) +(faint pink color) ++(obvious pink color) +++ (dark pink color)

Figure S1

Images taken after 4 days at 37 ºC on 7H11 agar

Gene *49*; Score *0*

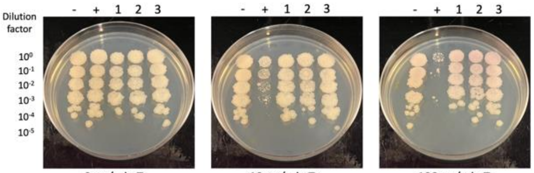


| Lane | Plasmid name | Gene name, replicate | Toxic/Non-toxic | Colony color on 100 ng/ml aTc plate* |
|---|---|---|---|---|
| - Non-toxic control | pExTra03 | Fruitloop *52* mutant | Non-toxic | + |
| + Toxic control | pExTra02 | Fruitloop *52* | Toxic | - |
| 1 | pExTra-NormanBulbieJr49 | NormanBulbieJr *49* replicate 1 | Non-toxic | + |
| 2 | pExTra-NormanBulbieJr49 | NormanBulbieJr *49* replicate 2 | Non-toxic | + |
| 3 | pExTra-NormanBulbieJr49 | NormanBulbieJr *49* replicate 3 | Non-toxic | + |

*Key: NG (no growth) - (no pink color) +(faint pink color) ++(obvious pink color) +++ (dark pink color)

Images taken after 3 days at 37 ºC on 7H11 agar

Gene *50*; Score *0*

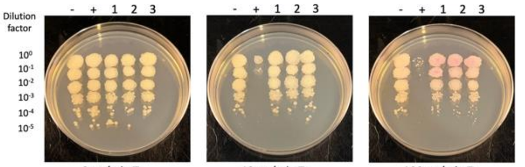


| Lane | Plasmid name | Gene name, replicate | Toxic/Non-toxic | Colony color on 100 ng/ml aTc plate* |
|---|---|---|---|---|
| - Non-toxic control | pExTra03 | Fruitloop *52* mutant | Non-toxic | - |
| + Toxic control | pExTra02 | Fruitloop *52* | Toxic | - |
| 1 | pExTra-NormanBulbieJr50 | NormanBulbieJr *50* replicate 1 | Non-toxic | ++ |
| 2 | pExTra-NormanBulbieJr50 | NormanBulbieJr *50* replicate 2 | Non-toxic | ++ |
| 3 | pExTra-NormanBulbieJr50 | NormanBulbieJr *50* replicate 3 | Non-toxic | ++ |

*Key: NG (no growth) - (no pink color) +(faint pink color) ++(obvious pink color) +++ (dark pink color)

Images taken after 4 days at 37 ºC on 7H11 agar

Gene *51*; Score *0*

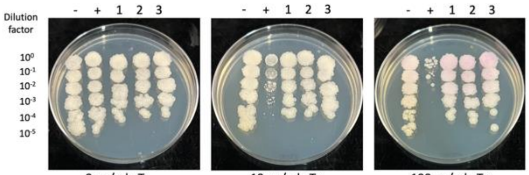


| Lane | Plasmid name | Gene name, replicate | Toxic/Non-toxic | Colony color on 100 ng/ml aTc plate* |
|---|---|---|---|---|
| - Non-toxic control | pExTra03 | Fruitloop *52* mutant | Non-toxic | + |
| + Toxic control | pExTra02 | Fruitloop *52* | Toxic | - |
| 1 | pExTra-NormanBulbieJr51 | NormanBulbieJr *51* replicate 1 | Non-toxic | + |
| 2 | pExTra-NormanBulbieJr51 | NormanBulbieJr *51* replicate 2 | Non-toxic | + |
| 3 | pExTra-NormanBulbieJr51 | NormanBulbieJr *51* replicate 3 | Non-toxic | + |

*Key: NG (no growth) - (no pink color) +(faint pink color) ++(obvious pink color) +++ (dark pink color)

Images taken after 4 days at 37 ºC on 7H11 agar

Gene *52*; Score *0*

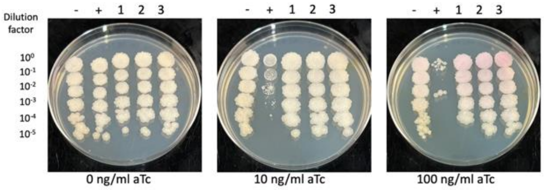


| Lane | Plasmid name | Gene name, replicate | Toxic/Non-toxic | Colony color on 100 ng/ml aTc plate* |
|---|---|---|---|---|
| - Non-toxic control | pExTra03 | Fruitloop *52* mutant | Non-toxic | + |
| + Toxic control | pExTra02 | Fruitloop *52* | Toxic | - |
| 1 | pExTra-NormanBulbieJr52 | NormanBulbieJr *52* replicate 1 | Non-toxic | + |
| 2 | pExTra-NormanBulbieJr52 | NormanBulbieJr *52* replicate 2 | Non-toxic | + |
| 3 | pExTra-NormanBulbieJr52 | NormanBulbieJr *52* replicate 3 | Non-toxic | + |

*Key: NG (no growth) - (no pink color) +(faint pink color) ++(obvious pink color) +++ (dark pink color)

Images taken after 4 days at 37 ºC on 7H11 agar

Gene *53*; Score *1*

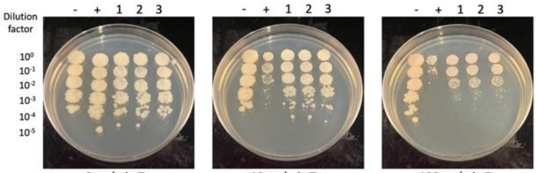


| Lane | Plasmid name | Gene name, replicate | Toxic/Non-toxic | Colony color on 100 ng/ml aTc plate* |
|---|---|---|---|---|
| - Non-toxic control | pExTra03 | Fruitloop *52* mutant | Non-toxic | + |
| + Toxic control | pExTra02 | Fruitloop *52* | Toxic | - |
| 1 | pExTra-NormanBulbieJr53 | NormanBulbieJr *53* replicate 1 | Toxic | + |
| 2 | pExTra-NormanBulbieJr53 | NormanBulbieJr *53* replicate 2 | Toxic | + |
| 3 | pExTra-NormanBulbieJr53 | NormanBulbieJr *53* replicate 3 | Toxic | + |

*Key: NG (no growth) - (no pink color) +(faint pink color) ++(obvious pink color) +++ (dark pink color)

Images taken after 3 days at 37 ºC on 7H11 agar

Gene *54*; Score *3*

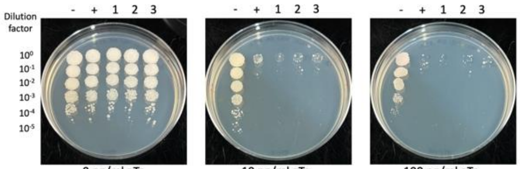


| Lane | Plasmid name | Gene name, replicate | Toxic/Non-toxic | Colony color on 100 ng/ml aTc plate* |
|---|---|---|---|---|
| - Non-toxic control | pExTra03 | Fruitloop *52* mutant | Non-toxic | + |
| + Toxic control | pExTra02 | Fruitloop *52* | Toxic | NG |
| 1 | pExTra-NormanBulbieJr54 | NormanBulbieJr *54* replicate 1 | Toxic | NG |
| 2 | pExTra-NormanBulbieJr54 | NormanBulbieJr *54* replicate 2 | Toxic | NG |
| 3 | pExTra-NormanBulbieJr54 | NormanBulbieJr *54* replicate 3 | Toxic | NG |

*Key: NG (no growth) - (no pink color) +(faint pink color) ++(obvious pink color) +++ (dark pink color)

Images taken after 3 days at 37 ºC on 7H11 agar

Gene *55*; Score *0*

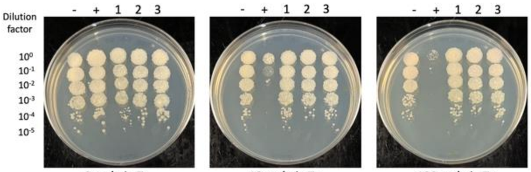


| Lane | Plasmid name | Gene name, replicate | Toxic/Non-toxic | Colony color on 100 ng/ml aTc plate* |
|---|---|---|---|---|
| - Non-toxic control | pExTra03 | Fruitloop *52* mutant | Non-toxic | + |
| + Toxic control | pExTra02 | Fruitloop *52* | Toxic | - |
| 1 | pExTra-NormanBulbieJr55 | NormanBulbieJr *55* replicate 1 | Non-toxic | - |
| 2 | pExTra-NormanBulbieJr55 | NormanBulbieJr *55* replicate 2 | Non-toxic | - |
| 3 | pExTra-NormanBulbieJr55 | NormanBulbieJr *55* replicate 3 | Non-toxic | - |

*Key: NG (no growth) - (no pink color) +(faint pink color) ++(obvious pink color) +++ (dark pink color)

Images taken after 3 days at 37 ºC on 7H11 agar

Gene *56*; Score *3*

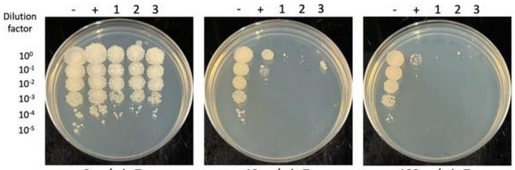


| Lane | Plasmid name | Gene name, replicate | Toxic/Non-toxic | Colony color on 100 ng/ml aTc plate* |
|---|---|---|---|---|
| - Non-toxic control | pExTra03 | Fruitloop *52* mutant | Non-toxic | + |
| + Toxic control | pExTra02 | Fruitloop *52* | Toxic | - |
| 1 | pExTra-NormanBulbieJr56 | NormanBulbieJr *56* replicate 1 | Toxic | NG |
| 2 | pExTra-NormanBulbieJr56 | NormanBulbieJr *56* replicate 2 | Toxic | NG |
| 3 | pExTra-NormanBulbieJr56 | NormanBulbieJr *56* replicate 3 | Toxic | NG |

*Key: NG (no growth) - (no pink color) +(faint pink color) ++(obvious pink color) +++ (dark pink color)

Figure S1

Gene *57*; Score *0*

Images taken after 4 days at 37 ºC on 7H11 agar

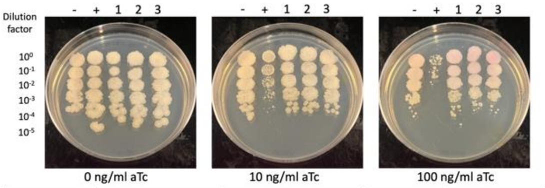


| Lane | Plasmid name | Gene name, replicate | Toxic/Non-toxic | Colony color on 100 ng/ml aTc plate* |
|---|---|---|---|---|
| - Non-toxic control | pExTra03 | Fruitloop *52* mutant | Non-toxic | + |
| + Toxic control | pExTra02 | Fruitloop *52* | Toxic | - |
| 1 | pExTra-NormanBulbieJr57 | NormanBulbieJr *57* replicate 1 | Non-toxic | + |
| 2 | pExTra-NormanBulbieJr57 | NormanBulbieJr *57* replicate 2 | Non-toxic | + |
| 3 | pExTra-NormanBulbieJr57 | NormanBulbieJr *57* replicate 3 | Non-toxic | + |

*Key: NG (no growth) - (no pink color) +(faint pink color) ++(obvious pink color) +++ (dark pink color)

Gene *61*; Score *0*

Images taken after 4 days at 37 ºC on 7H11 agar

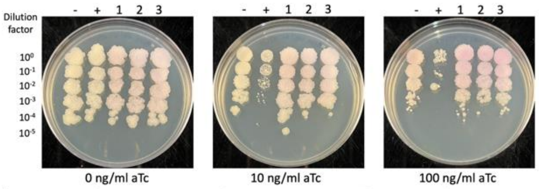


| Lane | Plasmid name | Gene name, replicate | Toxic/Non-toxic | Colony color on 100 ng/ml aTc plate* |
|---|---|---|---|---|
| - Non-toxic control | pExTra03 | Fruitloop *52* mutant | Non-toxic | + |
| + Toxic control | pExTra02 | Fruitloop *52* | Toxic | - |
| 1 | pExTra-NormanBulbieJr61 | NormanBulbieJr *61* replicate 1 | Non-toxic | ++ |
| 2 | pExTra-NormanBulbieJr61 | NormanBulbieJr *61* replicate 2 | Non-toxic | ++ |
| 3 | pExTra-NormanBulbieJr61 | NormanBulbieJr *61* replicate 3 | Non-toxic | ++ |

*Key: NG (no growth) - (no pink color) +(faint pink color) ++(obvious pink color) +++ (dark pink color)

Gene *58*; Score *2*

Images taken after 4 days at 37 ºC on 7H11 agar

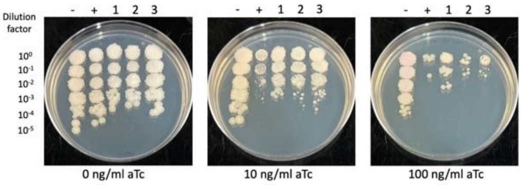


| Lane | Plasmid name | Gene name, replicate | Toxic/Non-toxic | Colony color on 100 ng/ml aTc plate* |
|---|---|---|---|---|
| - Non-toxic control | pExTra03 | Fruitloop *52* mutant | Non-toxic | + |
| + Toxic control | pExTra02 | Fruitloop *52* | Toxic | - |
| 1 | pExTra-NormanBulbieJr58 | NormanBulbieJr *58* replicate 1 | Toxic | - |
| 2 | pExTra-NormanBulbieJr58 | NormanBulbieJr *58* replicate 2 | Toxic | - |
| 3 | pExTra-NormanBulbieJr58 | NormanBulbieJr *58* replicate 3 | Toxic | - |

*Key: NG (no growth) - (no pink color) +(faint pink color) ++(obvious pink color) +++ (dark pink color)

Gene *62*; Score *0*

Images taken after 4 days at 37 ºC on 7H11 agar

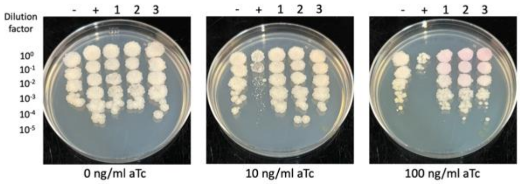


| Lane | Plasmid name | Gene name, replicate | Toxic/Non-toxic | Colony color on 100 ng/ml aTc plate* |
|---|---|---|---|---|
| - Non-toxic control | pExTra03 | Fruitloop *52* mutant | Non-toxic | - |
| + Toxic control | pExTra02 | Fruitloop *52* | Toxic | - |
| 1 | pExTra-NormanBulbieJr62 | NormanBulbieJr *62* replicate 1 | Non-toxic | + |
| 2 | pExTra-NormanBulbieJr62 | NormanBulbieJr *62* replicate 2 | Non-toxic | + |
| 3 | pExTra-NormanBulbieJr62 | NormanBulbieJr *62* replicate 3 | Non-toxic | + |

*Key: NG (no growth) - (no pink color) +(faint pink color) ++(obvious pink color) +++ (dark pink color)

Gene *59*; Score *3*

Images taken after 3 days at 37 ºC on 7H11 agar

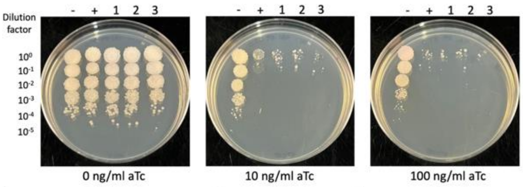


| Lane | Plasmid name | Gene name, replicate | Toxic/Non-toxic | Colony color on 100 ng/ml aTc plate* |
|---|---|---|---|---|
| - Non-toxic control | pExTra03 | Fruitloop *52* mutant | Non-toxic | + |
| + Toxic control | pExTra02 | Fruitloop *52* | Toxic | - |
| 1 | pExTra-NormanBulbieJr59 | NormanBulbieJr *59* replicate 1 | Toxic | - |
| 2 | pExTra-NormanBulbieJr59 | NormanBulbieJr *59* replicate 2 | Toxic | - |
| 3 | pExTra-NormanBulbieJr59 | NormanBulbieJr *59* replicate 3 | Toxic | - |

*Key: NG (no growth) - (no pink color) +(faint pink color) ++(obvious pink color) +++ (dark pink color)

Gene *63*; Score *0*

Images taken after 4 days at 37 ºC on 7H11 agar

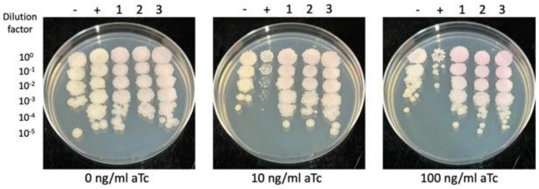


| Lane | Plasmid name | Gene name, replicate | Toxic/Non-toxic | Colony color on 100 ng/ml aTc plate* |
|---|---|---|---|---|
| - Non-toxic control | pExTra03 | Fruitloop *52* mutant | Non-toxic | + |
| + Toxic control | pExTra02 | Fruitloop *52* | Toxic | - |
| 1 | pExTra-NormanBulbieJr63 | NormanBulbieJr *63* replicate 1 | Non-toxic | ++ |
| 2 | pExTra-NormanBulbieJr63 | NormanBulbieJr *63* replicate 2 | Non-toxic | ++ |
| 3 | pExTra-NormanBulbieJr63 | NormanBulbieJr *63* replicate 3 | Non-toxic | ++ |

*Key: NG (no growth) - (no pink color) +(faint pink color) ++(obvious pink color) +++ (dark pink color)

Gene *60*; Score *1*

Images taken after 3 days at 37 ºC on 7H11 agar

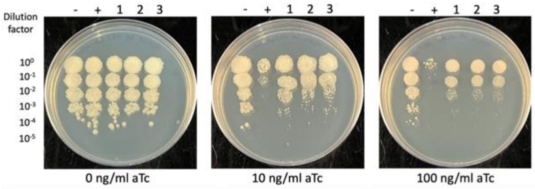


| Lane | Plasmid name | Gene name, replicate | Toxic/Non-toxic | Colony color on 100 ng/ml aTc plate* |
|---|---|---|---|---|
| - Non-toxic control | pExTra03 | Fruitloop *52* mutant | Non-toxic | + |
| + Toxic control | pExTra02 | Fruitloop *52* | Toxic | - |
| 1 | pExTra-NormanBulbieJr60 | NormanBulbieJr *60* replicate 1 | Toxic | + |
| 2 | pExTra-NormanBulbieJr60 | NormanBulbieJr *60* replicate 2 | Toxic | + |
| 3 | pExTra-NormanBulbieJr60 | NormanBulbieJr *60* replicate 3 | Toxic | + |

*Key: NG (no growth) - (no pink color) +(faint pink color) ++(obvious pink color) +++ (dark pink color)

Gene *64*; Score *2*

Images taken after 4 days at 37 ºC on 7H11 agar

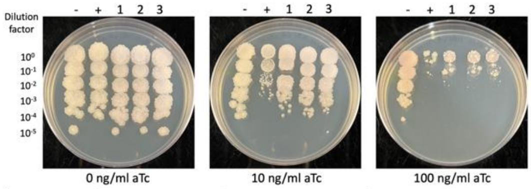


| Lane | Plasmid name | Gene name, replicate | Toxic/Non-toxic | Colony color on 100 ng/ml aTc plate* |
|---|---|---|---|---|
| - Non-toxic control | pExTra03 | Fruitloop *52* mutant | Non-toxic | + |
| + Toxic control | pExTra02 | Fruitloop *52* | Toxic | - |
| 1 | pExTra-NormanBulbieJr64 | NormanBulbieJr *64* replicate 1 | Toxic | + |
| 2 | pExTra-NormanBulbieJr64 | NormanBulbieJr *64* replicate 2 | Toxic | + |
| 3 | pExTra-NormanBulbieJr64 | NormanBulbieJr *64* replicate 3 | Toxic | + |

*Key: NG (no growth) - (no pink color) +(faint pink color) ++(obvious pink color) +++ (dark pink color)

### Figure S1

#### Gene *65*; Score *2*

Images taken after 5 days at 37 ºC on 7H11 agar

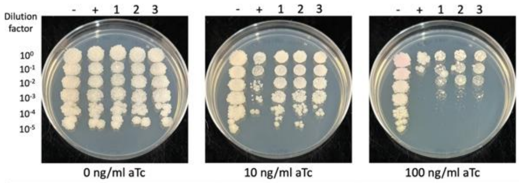


| Lane | Plasmid name | Gene name, replicate | Toxic/Non-toxic | Colony color on 100 ng/ml aTc plate* |
|---|---|---|---|---|
| - Non-toxic control | pExTra03 | Fruitloop *52* mutant | Non-toxic | + |
| + Toxic control | pExTra02 | Fruitloop *52* | Toxic | - |
| 1 | pExTra-NormanBulbieJr65 | NormanBulbieJr *65* replicate 1 | Toxic | - |
| 2 | pExTra-NormanBulbieJr65 | NormanBulbieJr *65* replicate 2 | Toxic | - |
| 3 | pExTra-NormanBulbieJr65 | NormanBulbieJr *65* replicate 3 | Toxic | - |

*Key: NG (no growth) - (no pink color) +(faint pink color) ++(obvious pink color) +++ (dark pink color)

#### Gene *69*; Score *0*

Images taken after 4 days at 37 ºC on 7H11 agar

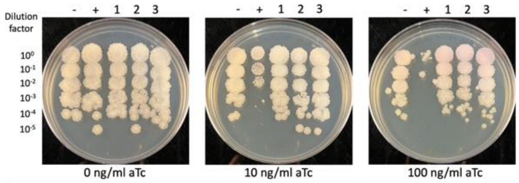


| Lane | Plasmid name | Gene name, replicate | Toxic/Non-toxic | Colony color on 100 ng/ml aTc plate* |
|---|---|---|---|---|
| - Non-toxic control | pExTra03 | Fruitloop *52* mutant | Non-toxic | + |
| + Toxic control | pExTra02 | Fruitloop *52* | Toxic | - |
| 1 | pExTra-NormanBulbieJr69 | NormanBulbieJr *69* replicate 1 | Non-toxic | + |
| 2 | pExTra-NormanBulbieJr69 | NormanBulbieJr *69* replicate 2 | Non-toxic | + |
| 3 | pExTra-NormanBulbieJr69 | NormanBulbieJr *69* replicate 3 | Non-toxic | + |

*Key: NG (no growth) - (no pink color) +(faint pink color) ++(obvious pink color) +++ (dark pink color)

#### Gene *66*; Score *0*

Images taken after 4 days at 37 ºC on 7H11 agar

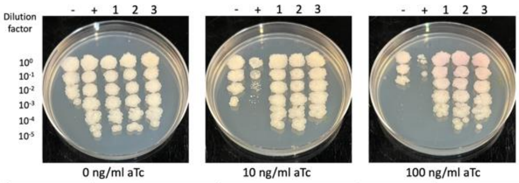


| Lane | Plasmid name | Gene name, replicate | Toxic/Non-toxic | Colony color on 100 ng/ml aTc plate* |
|---|---|---|---|---|
| - Non-toxic control | pExTra03 | Fruitloop *52* mutant | Non-toxic | + |
| + Toxic control | pExTra02 | Fruitloop *52* | Toxic | - |
| 1 | pExTra-NormanBulbieJr66 | NormanBulbieJr *66* replicate 1 | Non-toxic | + |
| 2 | pExTra-NormanBulbieJr66 | NormanBulbieJr *66* replicate 2 | Non-toxic | + |
| 3 | pExTra-NormanBulbieJr66 | NormanBulbieJr *66* replicate 3 | Non-toxic | + |

*Key: NG (no growth) - (no pink color) +(faint pink color) ++(obvious pink color) +++ (dark pink color)

#### Gene *70*; Score *2*

Images taken after 3 days at 37 ºC on 7H11 agar

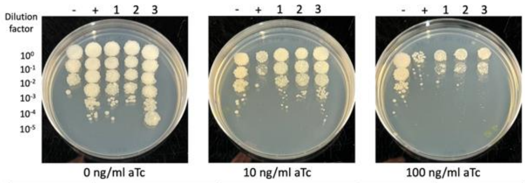


| Lane | Plasmid name | Gene name, replicate | Toxic/Non-toxic | Colony color on 100 ng/ml aTc plate* |
|---|---|---|---|---|
| - Non-toxic control | pExTra03 | Fruitloop *52* mutant | Non-toxic | - |
| + Toxic control | pExTra02 | Fruitloop *52* | Toxic | - |
| 1 | pExTra-NormanBulbieJr70 | NormanBulbieJr *70* replicate 1 | Toxic | - |
| 2 | pExTra-NormanBulbieJr70 | NormanBulbieJr *70* replicate 2 | Toxic | - |
| 3 | pExTra-NormanBulbieJr70 | NormanBulbieJr *70* replicate 3 | Toxic | - |

*Key: NG (no growth) - (no pink color) +(faint pink color) ++(obvious pink color) +++ (dark pink color)

#### Gene *67*; Score *0*

Images taken after 3 days at 37 ºC on 7H11 agar

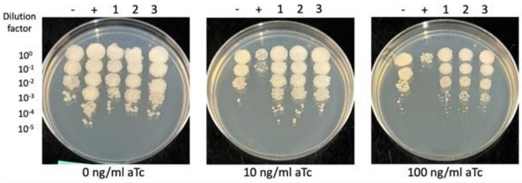


| Lane | Plasmid name | Gene name, replicate | Toxic/Non-toxic | Colony color on 100 ng/ml aTc plate* |
|---|---|---|---|---|
| - Non-toxic control | pExTra03 | Fruitloop *52* mutant | Non-toxic | - |
| + Toxic control | pExTra02 | Fruitloop *52* | Toxic | - |
| 1 | pExTra-NormanBulbieJr67 | NormanBulbieJr *67* replicate 1 | Non-toxic | + |
| 2 | pExTra-NormanBulbieJr67 | NormanBulbieJr *67* replicate 2 | Non-toxic | + |
| 3 | pExTra-NormanBulbieJr67 | NormanBulbieJr *67* replicate 3 | Non-toxic | + |

*Key: NG (no growth) - (no pink color) +(faint pink color) ++(obvious pink color) +++ (dark pink color)

#### Gene *71*; Score *3*

Images taken after 3 days at 37 ºC on 7H11 agar

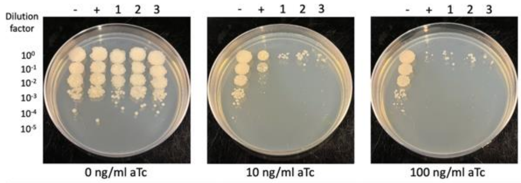


| Lane | Plasmid name | Gene name, replicate | Toxic/Non-toxic | Colony color on 100 ng/ml aTc plate* |
|---|---|---|---|---|
| - Non-toxic control | pExTra03 | Fruitloop *52* mutant | Non-toxic | + |
| + Toxic control | pExTra02 | Fruitloop *52* | Toxic | - |
| 1 | pExTra-NormanBulbieJr71 | NormanBulbieJr *71* replicate 1 | Toxic | - |
| 2 | pExTra-NormanBulbieJr71 | NormanBulbieJr *71* replicate 2 | Toxic | - |
| 3 | pExTra-NormanBulbieJr71 | NormanBulbieJr *71* replicate 3 | Toxic | - |

*Key: NG (no growth) - (no pink color) +(faint pink color) ++(obvious pink color) +++ (dark pink color)

#### Gene *68*; Score *3*

Images taken after 3 days at 37 ºC on 7H11 agar

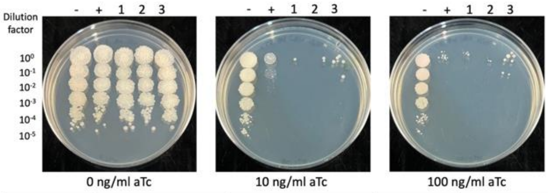


| Lane | Plasmid name | Gene name, replicate | Toxic/Non-toxic | Colony color on 100 ng/ml aTc plate* |
|---|---|---|---|---|
| - Non-toxic control | pExTra03 | Fruitloop *52* mutant | Non-toxic | + |
| + Toxic control | pExTra02 | Fruitloop *52* | Toxic | - |
| 1 | pExTra-NormanBulbieJr68 | NormanBulbieJr *68* replicate 1 | Toxic | - |
| 2 | pExTra-NormanBulbieJr68 | NormanBulbieJr *68* replicate 2 | Toxic | NG |
| 3 | pExTra-NormanBulbieJr68 | NormanBulbieJr *68* replicate 3 | Toxic | - |

*Key: NG (no growth) - (no pink color) +(faint pink color) ++(obvious pink color) +++ (dark pink color)

#### Gene *72*; Score *3*

Images taken after 3 days at 37 ºC on 7H11 agar

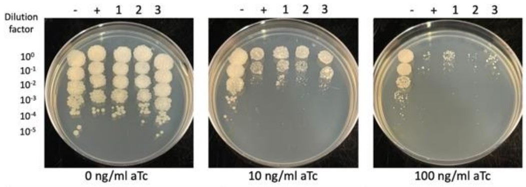


| Lane | Plasmid name | Gene name, replicate | Toxic/Non-toxic | Colony color on 100 ng/ml aTc plate* |
|---|---|---|---|---|
| - Non-toxic control | pExTra03 | Fruitloop *52* mutant | Non-toxic | - |
| + Toxic control | pExTra02 | Fruitloop *52* | Toxic | - |
| 1 | pExTra-NormanBulbieJr72 | NormanBulbieJr *72* replicate 1 | Toxic | - |
| 2 | pExTra-NormanBulbieJr72 | NormanBulbieJr *72* replicate 2 | Toxic | - |
| 3 | pExTra-NormanBulbieJr72 | NormanBulbieJr *72* replicate 3 | Toxic | - |

*Key: NG (no growth) - (no pink color) +(faint pink color) ++(obvious pink color) +++ (dark pink color)

Figure S1

Images taken after 3 days at 37 ºC on 7H11 agar

Gene *73*; Score *0*

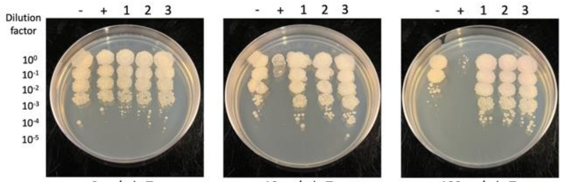


| Lane | Plasmid name | Gene name, replicate | Toxic/Non-toxic | Colony color on 100 ng/ml aTc plate* |
|---|---|---|---|---|
| - Non-toxic control | pExTra03 | Fruitloop *52* mutant | Non-toxic | + |
| + Toxic control | pExTra02 | Fruitloop *52* | Toxic | - |
| 1 | pExTra-NormanBulbieJr73 | NormanBulbieJr *73* replicate 1 | Non-toxic | + |
| 2 | pExTra-NormanBulbieJr73 | NormanBulbieJr *73* replicate 2 | Non-toxic | + |
| 3 | pExTra-NormanBulbieJr73 | NormanBulbieJr *73* replicate 3 | Non-toxic | + |

*Key: NG (no growth) - (no pink color) +(faint pink color) ++(obvious pink color) +++ (dark pink color)

Images taken after 4 days at 37 ºC on 7H11 agar

Gene *74*; Score *0*

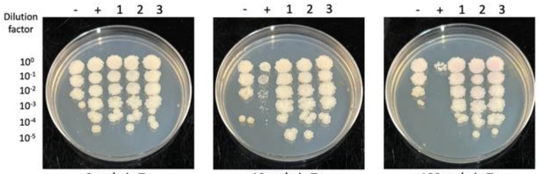


| Lane | Plasmid name | Gene name, replicate | Toxic/Non-toxic | Colony color on 100 ng/ml aTc plate* |
|---|---|---|---|---|
| - Non-toxic control | pExTra03 | Fruitloop *52* mutant | Non-toxic | + |
| + Toxic control | pExTra02 | Fruitloop *52* | Toxic | - |
| 1 | pExTra-NormanBulbieJr74 | NormanBulbieJr *74* replicate 1 | Non-toxic | + |
| 2 | pExTra-NormanBulbieJr74 | NormanBulbieJr *74* replicate 2 | Non-toxic | + |
| 3 | pExTra-NormanBulbieJr74 | NormanBulbieJr *74* replicate 3 | Non-toxic | + |

*Key: NG (no growth) - (no pink color) +(faint pink color) ++(obvious pink color) +++ (dark pink color)

Images taken after 4 days at 37 ºC on 7H11 agar

Gene *75*; Score *0*

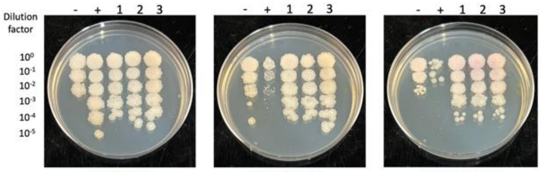


| Lane | Plasmid name | Gene name, replicate | Toxic/Non-toxic | Colony color on 100 ng/ml aTc plate* |
|---|---|---|---|---|
| - Non-toxic control | pExTra03 | Fruitloop *52* mutant | Non-toxic | - |
| + Toxic control | pExTra02 | Fruitloop *52* | Toxic | - |
| 1 | pExTra-NormanBulbieJr75 | NormanBulbieJr *75* replicate 1 | Non-toxic | + |
| 2 | pExTra-NormanBulbieJr75 | NormanBulbieJr *75* replicate 2 | Non-toxic | + |
| 3 | pExTra-NormanBulbieJr75 | NormanBulbieJr *75* replicate 3 | Non-toxic | + |

*Key: NG (no growth) - (no pink color) +(faint pink color) ++(obvious pink color) +++ (dark pink color)

Images taken after 4 days at 37 ºC on 7H11 agar

Gene *76*; Score *0*

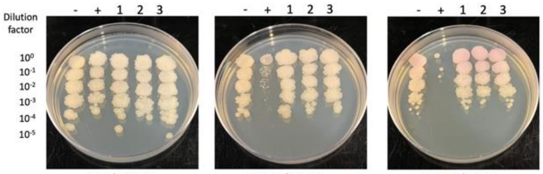


| Lane | Plasmid name | Gene name, replicate | Toxic/Non-toxic | Colony color on 100 ng/ml aTc plate* |
|---|---|---|---|---|
| - Non-toxic control | pExTra03 | Fruitloop *52* mutant | Non-toxic | + |
| + Toxic control | pExTra02 | Fruitloop *52* | Toxic | - |
| 1 | pExTra-NormanBulbieJr76 | NormanBulbieJr *76* replicate 1 | Non-toxic | ++ |
| 2 | pExTra-NormanBulbieJr76 | NormanBulbieJr *76* replicate 2 | Non-toxic | ++ |
| 3 | pExTra-NormanBulbieJr76 | NormanBulbieJr *76* replicate 3 | Non-toxic | ++ |

*Key: NG (no growth) - (no pink color) +(faint pink color) ++(obvious pink color) +++ (dark pink color)

Images taken after 4 days at 37 ºC on 7H11 agar

Gene *77*; Score *0*

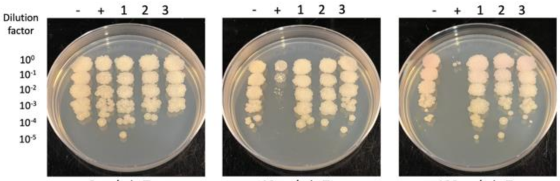


| Lane | Plasmid name | Gene name, replicate | Toxic/Non-toxic | Colony color on 100 ng/ml aTc plate* |
|---|---|---|---|---|
| - Non-toxic control | pExTra03 | Fruitloop *52* mutant | Non-toxic | ++ |
| + Toxic control | pExTra02 | Fruitloop *52* | Toxic | - |
| 1 | pExTra-NormanBulbieJr77 | NormanBulbieJr *77* replicate 1 | Non-toxic | + |
| 2 | pExTra-NormanBulbieJr77 | NormanBulbieJr *77* replicate 2 | Non-toxic | + |
| 3 | pExTra-NormanBulbieJr77 | NormanBulbieJr *77* replicate 3 | Non-toxic | + |

*Key: NG (no growth) - (no pink color) +(faint pink color) ++(obvious pink color) +++ (dark pink color)

Images taken after 4 days at 37 ºC on 7H11 agar

Gene *78*; Score *0*

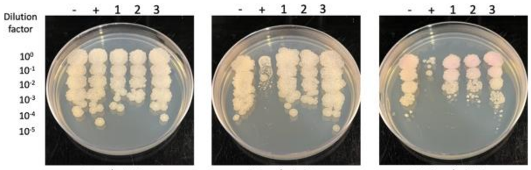


| Lane | Plasmid name | Gene name, replicate | Toxic/Non-toxic | Colony color on 100 ng/ml aTc plate* |
|---|---|---|---|---|
| - Non-toxic control | pExTra03 | Fruitloop *52* mutant | Non-toxic | + |
| + Toxic control | pExTra02 | Fruitloop *52* | Toxic | - |
| 1 | pExTra-NormanBulbieJr78 | NormanBulbieJr *78* replicate 1 | Non-toxic | + |
| 2 | pExTra-NormanBulbieJr78 | NormanBulbieJr *78* replicate 2 | Non-toxic | + |
| 3 | pExTra-NormanBulbieJr78 | NormanBulbieJr *78* replicate 3 | Non-toxic | + |

*Key: NG (no growth) - (no pink color) +(faint pink color) ++(obvious pink color) +++ (dark pink color)

Images taken after 3 days at 37 ºC on 7H11 agar

Gene *79*; Score *0*

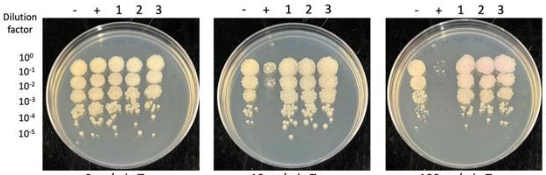


| Lane | Plasmid name | Gene name, replicate | Toxic/Non-toxic | Colony color on 100 ng/ml aTc plate* |
|---|---|---|---|---|
| - Non-toxic control | pExTra03 | Fruitloop *52* mutant | Non-toxic | - |
| + Toxic control | pExTra02 | Fruitloop *52* | Toxic | - |
| 1 | pExTra-NormanBulbieJr79 | NormanBulbieJr *79* replicate 1 | Non-toxic | + |
| 2 | pExTra-NormanBulbieJr79 | NormanBulbieJr *79* replicate 2 | Non-toxic | + |
| 3 | pExTra-NormanBulbieJr79 | NormanBulbieJr *79* replicate 3 | Non-toxic | + |

*Key: NG (no growth) - (no pink color) +(faint pink color) ++(obvious pink color) +++ (dark pink color)

Images taken after 4 days at 37 ºC on 7H11 agar

Gene *80*; Score *0*

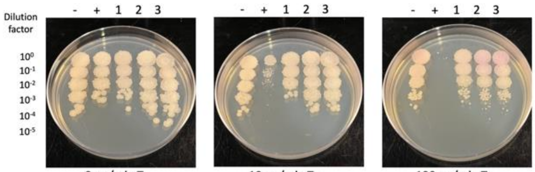


| Lane | Plasmid name | Gene name, replicate | Toxic/Non-toxic | Colony color on 100 ng/ml aTc plate* |
|---|---|---|---|---|
| - Non-toxic control | pExTra03 | Fruitloop *52* mutant | Non-toxic | + |
| + Toxic control | pExTra02 | Fruitloop *52* | Toxic | - |
| 1 | pExTra-NormanBulbieJr80 | NormanBulbieJr *80* replicate 1 | Non-toxic | + |
| 2 | pExTra-NormanBulbieJr80 | NormanBulbieJr *80* replicate 2 | Non-toxic | ++ |
| 3 | pExTra-NormanBulbieJr80 | NormanBulbieJr *80* replicate 3 | Non-toxic | ++ |

*Key: NG (no growth) - (no pink color) +(faint pink color) ++(obvious pink color) +++ (dark pink color)

Figure S1

#### Gene *81*; Score *0*

Images taken after 4 days at 37 ºC on 7H11 agar

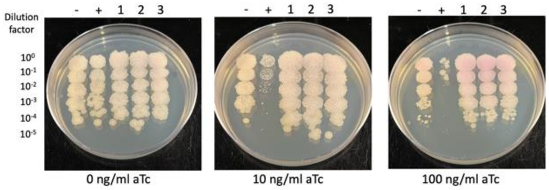


| Lane | Plasmid name | Gene name, replicate | Toxic/Non-toxic | Colony color on 100 ng/ml aTc plate* |
|---|---|---|---|---|
| - Non-toxic control | pExTra03 | Fruitloop *52* mutant | Non-toxic | + |
| + Toxic control | pExTra02 | Fruitloop *52* | Toxic | - |
| 1 | pExTra-NormanBulbieJr81 | NormanBulbieJr 81 replicate 1 | Non-toxic | ++ |
| 2 | pExTra-NormanBulbieJr81 | NormanBulbieJr 81 replicate 2 | Non-toxic | ++ |
| 3 | pExTra-NormanBulbieJr81 | NormanBulbieJr 81 replicate 3 | Non-toxic | ++ |

*Key: NG (no growth) - (no pink color) +(faint pink color) ++(obvious pink color) +++ (dark pink color)

#### Gene *85*; Score *0*

Images taken after 3 days at 37 ºC on 7H11 agar

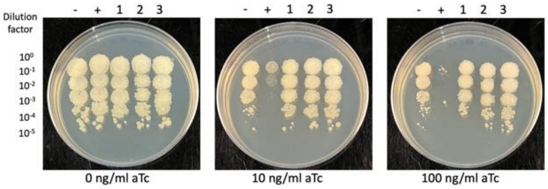


| Lane | Plasmid name | Gene name, replicate | Toxic/Non-toxic | Colony color on 100 ng/ml aTc plate* |
|---|---|---|---|---|
| - Non-toxic control | pExTra03 | Fruitloop *52* mutant | Non-toxic | - |
| + Toxic control | pExTra02 | Fruitloop *52* | Toxic | - |
| 1 | pExTra-NormanBulbieJr85 | NormanBulbieJr *85* replicate 1 | Non-Toxic | - |
| 2 | pExTra-NormanBulbieJr85 | NormanBulbieJr *85* replicate 2 | Non-Toxic | - |
| 3 | pExTra-NormanBulbieJr85 | NormanBulbieJr *85* replicate 3 | Non-Toxic | - |

*Key: NG (no growth) - (no pink color) +(faint pink color) ++(obvious pink color) +++ (dark pink color)

#### Gene *82*; Score *0*

Images taken after 4 days at 37 ºC on 7H11 agar

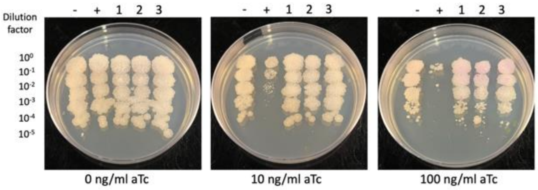


| Lane | Plasmid name | Gene name, replicate | Toxic/Non-toxic | Colony color on 100 ng/ml aTc plate* |
|---|---|---|---|---|
| - Non-toxic control | pExTra03 | Fruitloop *52* mutant | Non-toxic | + |
| + Toxic control | pExTra02 | Fruitloop *52* | Toxic | - |
| 1 | pExTra-NormanBulbieJr82 | NormanBulbieJr 82 replicate 1 | Non-toxic | ++ |
| 2 | pExTra-NormanBulbieJr82 | NormanBulbieJr 82 replicate 2 | Non-toxic | ++ |
| 3 | pExTra-NormanBulbieJr82 | NormanBulbieJr 82 replicate 3 | Non-toxic | ++ |

*Key: NG (no growth) - (no pink color) +(faint pink color) ++(obvious pink color) +++ (dark pink color)

#### Gene *86*; Score *0*

Images taken after 3 days at 37 ºC on 7H11 agar

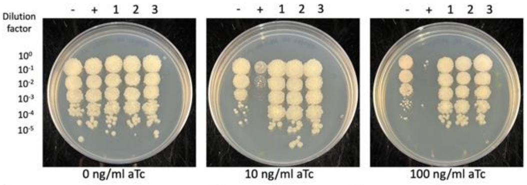


| Lane | Plasmid name | Gene name, replicate | Toxic/Non-toxic | Colony color on 100 ng/ml aTc plate* |
|---|---|---|---|---|
| - Non-toxic control | pExTra03 | Fruitloop *52* mutant | Non-toxic | + |
| + Toxic control | pExTra02 | Fruitloop *52* | Toxic | - |
| 1 | pExTra-NormanBulbieJr86 | NormanBulbieJr *86* replicate 1 | Non-toxic | - |
| 2 | pExTra-NormanBulbieJr86 | NormanBulbieJr *86* replicate 2 | Non-toxic | - |
| 3 | pExTra-NormanBulbieJr86 | NormanBulbieJr *86* replicate 3 | Non-toxic | - |

*Key: NG (no growth) - (no pink color) +(faint pink color) ++(obvious pink color) +++ (dark pink color)

#### Gene *83*; Score *0*

Images taken after 4 days at 37 ºC on 7H11 agar

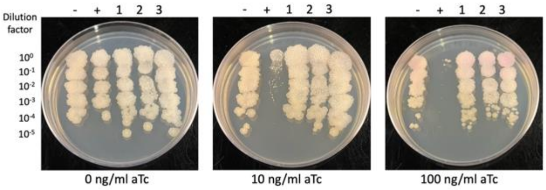


| Lane | Plasmid name | Gene name, replicate | Toxic/Non-toxic | Colony color on 100 ng/ml aTc plate* |
|---|---|---|---|---|
| - Non-toxic control | pExTra03 | Fruitloop *52* mutant | Non-toxic | + |
| + Toxic control | pExTra02 | Fruitloop *52* | Toxic | - |
| 1 | pExTra-NormanBulbieJr83 | NormanBulbieJr 83 replicate 1 | Non-toxic | + |
| 2 | pExTra-NormanBulbieJr83 | NormanBulbieJr 83 replicate 2 | Non-toxic | + |
| 3 | pExTra-NormanBulbieJr83 | NormanBulbieJr 83 replicate 3 | Non-toxic | + |

*Key: NG (no growth) - (no pink color) +(faint pink color) ++(obvious pink color) +++ (dark pink color)

#### Gene *87*; Score *0*

Images taken after 3 days at 37 ºC on 7H11 agar

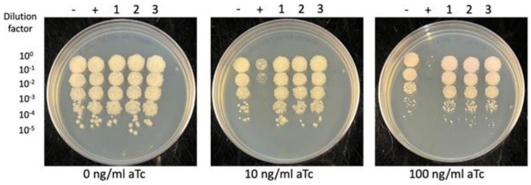


| Lane | Plasmid name | Gene name, replicate | Toxic/Non-toxic | Colony color on 100 ng/ml aTc plate* |
|---|---|---|---|---|
| - Non-toxic control | pExTra03 | Fruitloop *52* mutant | Non-toxic | + |
| + Toxic control | pExTra02 | Fruitloop *52* | Toxic | - |
| 1 | pExTra-NormanBulbieJr87 | NormanBulbieJr *87* replicate 1 | Non-toxic | + |
| 2 | pExTra-NormanBulbieJr87 | NormanBulbieJr *87* replicate 2 | Non-toxic | + |
| 3 | pExTra-NormanBulbieJr87 | NormanBulbieJr *87* replicate 3 | Non-toxic | + |

*Key: NG (no growth) - (no pink color) +(faint pink color) ++(obvious pink color) +++ (dark pink color)

#### Gene *84*; Score *0*

Images taken after 4 days at 37 ºC on 7H11 agar

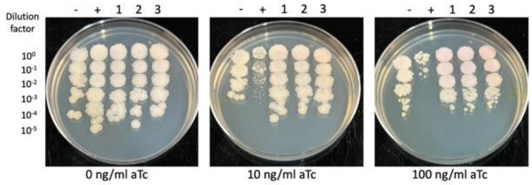


| Lane | Plasmid name | Gene name, replicate | Toxic/Non-toxic | Colony color on 100 ng/ml aTc plate* |
|---|---|---|---|---|
| - Non-toxic control | pExTra03 | Fruitloop *52* mutant | Non-toxic | + |
| + Toxic control | pExTra02 | Fruitloop *52* | Toxic | - |
| 1 | pExTra-NormanBulbieJr84 | NormanBulbieJr *84* replicate 1 | Non-toxic | + |
| 2 | pExTra-NormanBulbieJr84 | NormanBulbieJr *84* replicate 2 | Non-toxic | + |
| 3 | pExTra-NormanBulbieJr84 | NormanBulbieJr *84* replicate 3 | Non-toxic | + |

*Key: NG (no growth) - (no pink color) +(faint pink color) ++(obvious pink color) +++ (dark pink color)

#### Gene *88*; Score *0*

Images taken after 4 days at 37 ºC on 7H11 agar

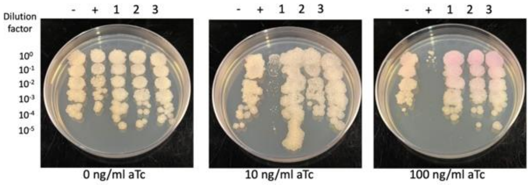


| Lane | Plasmid name | Gene name, replicate | Toxic/Non-toxic | Colony color on 100 ng/ml aTc plate* |
|---|---|---|---|---|
| - Non-toxic control | pExTra03 | Fruitloop *52* mutant | Non-toxic | + |
| + Toxic control | pExTra02 | Fruitloop *52* | Toxic | - |
| 1 | pExTra-NormanBulbieJr88 | NormanBulbieJr *88* replicate 1 | Non-toxic | ++ |
| 2 | pExTra-NormanBulbieJr88 | NormanBulbieJr *88* replicate 2 | Non-toxic | ++ |
| 3 | pExTra-NormanBulbieJr88 | NormanBulbieJr *88* replicate 3 | Non-toxic | ++ |

*Key: NG (no growth) - (no pink color) +(faint pink color) ++(obvious pink color) +++ (dark pink color)

### Figure S1

#### Gene *89*; Score *0*

Images taken after 3 days at 37 ºC on 7H11 agar

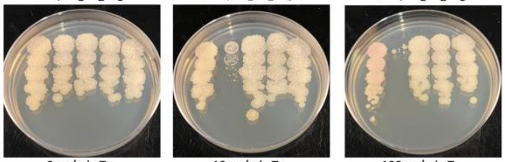


| Lane | Plasmid name | Gene name, replicate | Toxic/Non-toxic | Colony color on 100 ng/ml aTc plate* |
|---|---|---|---|---|
| - Non-toxic control | pExTra03 | Fruitloop *52* mutant | Non-toxic | + |
| + Toxic control | pExTra02 | Fruitloop *52* | Toxic | - |
| 1 | pExTra-NormanBulbieJr89 | NormanBulbieJr *89* replicate 1 | Non-toxic | - |
| 2 | pExTra-NormanBulbieJr89 | NormanBulbieJr *89* replicate 2 | Non-toxic | - |
| 3 | pExTra-NormanBulbieJr89 | NormanBulbieJr *89* replicate 3 | Non-toxic | - |

*Key: NG (no growth) - (no pink color) +(faint pink color) ++(obvious pink color) +++ (dark pink color)

#### Gene *93*; Score *0*

Images taken after 4 days at 37 ºC on 7H11 agar

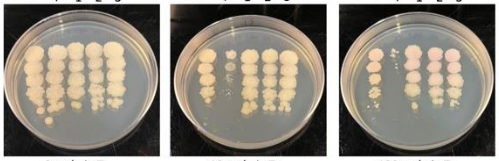


| Lane | Plasmid name | Gene name, replicate | Toxic/Non-toxic | Colony color on 100 ng/ml aTc plate* |
|---|---|---|---|---|
| - Non-toxic control | pExTra03 | Fruitloop *52* mutant | Non-toxic | + |
| + Toxic control | pExTra02 | Fruitloop *52* | Toxic | - |
| 1 | pExTra-NormanBulbieJr93 | NormanBulbieJr *93* replicate 1 | Non-toxic | ++ |
| 2 | pExTra-NormanBulbieJr93 | NormanBulbieJr *93* replicate 2 | Non-toxic | ++ |
| 3 | pExTra-NormanBulbieJr93 | NormanBulbieJr *93* replicate 3 | Non-toxic | ++ |

*Key: NG (no growth) - (no pink color) +(faint pink color) ++(obvious pink color) +++ (dark pink color)

#### Gene *90*; Score *2*

Images taken after 3 days at 37 ºC on 7H11 agar

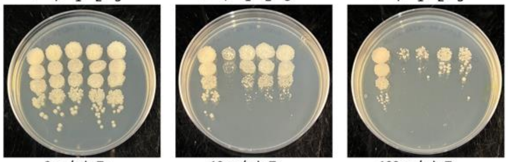


| Lane | Plasmid name | Gene name, replicate | Toxic/Non-toxic | Colony color on 100 ng/ml aTc plate* |
|---|---|---|---|---|
| - Non-toxic control | pExTra03 | Fruitloop *52* mutant | Non-toxic | - |
| + Toxic control | pExTra02 | Fruitloop *52* | Toxic | - |
| 1 | pExTra-NormanBulbieJr90 | NormanBulbieJr *90* replicate 1 | Toxic | - |
| 2 | pExTra-NormanBulbieJr90 | NormanBulbieJr *90* replicate 2 | Toxic | - |
| 3 | pExTra-NormanBulbieJr90 | NormanBulbieJr *90* replicate 3 | Toxic | - |

*Key: NG (no growth) - (no pink color) +(faint pink color) ++(obvious pink color) +++ (dark pink color)

#### Gene *94*; Score *0*

Images taken after 4 days at 37 ºC on 7H11 agar

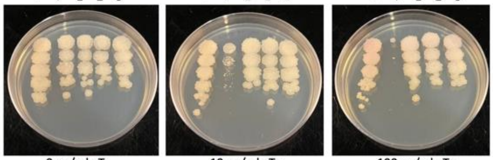


| Lane | Plasmid name | Gene name, replicate | Toxic/Non-toxic | Colony color on 100 ng/ml aTc plate* |
|---|---|---|---|---|
| - Non-toxic control | pExTra03 | Fruitloop *52* mutant | Non-toxic | ++ |
| + Toxic control | pExTra02 | Fruitloop *52* | Toxic | - |
| 1 | pExTra-NormanBulbieJr94 | NormanBulbieJr *94* replicate 1 | Non-toxic | + |
| 2 | pExTra-NormanBulbieJr94 | NormanBulbieJr *94* replicate 2 | Non-toxic | + |
| 3 | pExTra-NormanBulbieJr94 | NormanBulbieJr *94* replicate 3 | Non-toxic | + |

*Key: NG (no growth) - (no pink color) +(faint pink color) ++(obvious pink color) +++ (dark pink color)

#### Gene *91*; Score *0*

Images taken after 3 days at 37 ºC on 7H11 agar

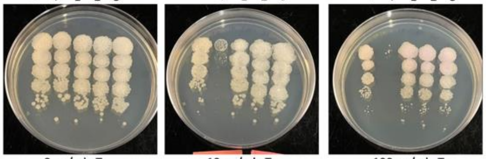


| Lane | Plasmid name | Gene name, replicate | Toxic/Non-toxic | Colony color on 100 ng/ml aTc plate* |
|---|---|---|---|---|
| - Non-toxic control | pExTra03 | Fruitloop *52* mutant | Non-toxic | + |
| + Toxic control | pExTra02 | Fruitloop *52* | Toxic | - |
| 1 | pExTra-NormanBulbieJr91 | NormanBulbieJr *91* replicate 1 | Non-toxic | + |
| 2 | pExTra-NormanBulbieJr91 | NormanBulbieJr *91* replicate 2 | Non-toxic | + |
| 3 | pExTra-NormanBulbieJr91 | NormanBulbieJr *91* replicate 3 | Non-toxic | + |

*Key: NG (no growth) - (no pink color) +(faint pink color) ++(obvious pink color) +++ (dark pink color)

#### Gene *95*; Score *0*

Images taken after 3 days at 37 ºC on 7H11 agar

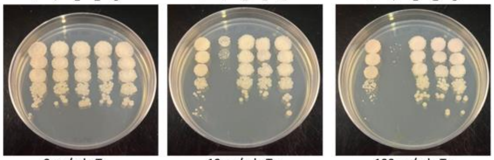


| Lane | Plasmid name | Gene name, replicate | Toxic/Non-toxic | Colony color on 100 ng/ml aTc plate* |
|---|---|---|---|---|
| - Non-toxic control | pExTra03 | Fruitloop *52* mutant | Non-toxic | ++ |
| + Toxic control | pExTra02 | Fruitloop *52* | Toxic | - |
| 1 | pExTra-NormanBulbieJr95 | NormanBulbieJr *95* replicate 1 | Non-toxic | + |
| 2 | pExTra-NormanBulbieJr95 | NormanBulbieJr *95* replicate 2 | Non-toxic | + |
| 3 | pExTra-NormanBulbieJr95 | NormanBulbieJr *95* replicate 3 | Non-toxic | + |

*Key: NG (no growth) - (no pink color) +(faint pink color) ++(obvious pink color) +++ (dark pink color)

#### Gene *92*; Score *0*

Images taken after 4 days at 37 ºC on 7H11 agar

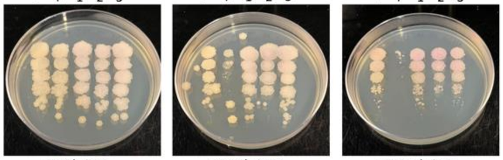


| Lane | Plasmid name | Gene name, replicate | Toxic/Non-toxic | Colony color on 100 ng/ml aTc plate* |
|---|---|---|---|---|
| - Non-toxic control | pExTra03 | Fruitloop *52* mutant | Non-toxic | + |
| + Toxic control | pExTra02 | Fruitloop *52* | Toxic | - |
| 1 | pExTra-NormanBulbieJr92 | NormanBulbieJr *92* replicate 1 | Non-toxic | + |
| 2 | pExTra-NormanBulbieJr92 | NormanBulbieJr *92* replicate 2 | Non-toxic | + |
| 3 | pExTra-NormanBulbieJr92 | NormanBulbieJr *92* replicate 3 | Non-toxic | + |

*Key: NG (no growth) - (no pink color) +(faint pink color) ++(obvious pink color) +++ (dark pink color)

#### Gene *96*; Score *0*

Images taken after 4 days at 37 ºC on 7H11 agar

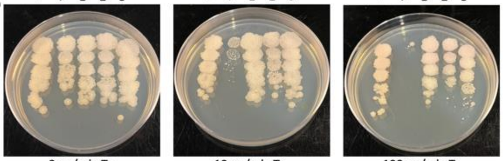


| Lane | Plasmid name | Gene name, replicate | Toxic/Non-toxic | Colony color on 100 ng/ml aTc plate* |
|---|---|---|---|---|
| - Non-toxic control | pExTra03 | Fruitloop *52* mutant | Non-toxic | + |
| + Toxic control | pExTra02 | Fruitloop *52* | Toxic | - |
| 1 | pExTra-NormanBulbieJr96 | NormanBulbieJr *96* replicate 1 | Non-toxic | + |
| 2 | pExTra-NormanBulbieJr96 | NormanBulbieJr *96* replicate 2 | Non-toxic | + |
| 3 | pExTra-NormanBulbieJr96 | NormanBulbieJr *96* replicate 3 | Non-toxic | + |

*Key: NG (no growth) - (no pink color) +(faint pink color) ++(obvious pink color) +++ (dark pink color)

### Figure S1

Gene *97*; Score *0*

Images taken after 4 days at 37 ºC on 7H11 agar

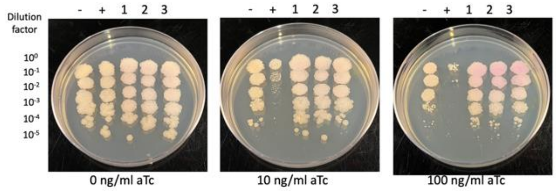


| Lane | Plasmid name | Gene name, replicate | Toxic/Non-toxic | Colony color on 100 ng/ml aTc plate* |
|---|---|---|---|---|
| - Non-toxic control | pExTra03 | Fruitloop *52* mutant | Non-toxic | - |
| + Toxic control | pExTra02 | Fruitloop *52* | Toxic | - |
| 1 | pExTra-NormanBulbieJr97 | NormanBulbieJr *97* replicate 1 | Non-toxic | ++ |
| 2 | pExTra-NormanBulbieJr97 | NormanBulbieJr *97* replicate 2 | Non-toxic | ++ |
| 3 | pExTra-NormanBulbieJr97 | NormanBulbieJr *97* replicate 3 | Non-toxic | ++ |

*Key: NG (no growth) - (no pink color) +(faint pink color) ++(obvious pink color) +++ (dark pink color)

Gene *101*; Score *0*

Images taken after 3 days at 37 ºC on 7H11 agar

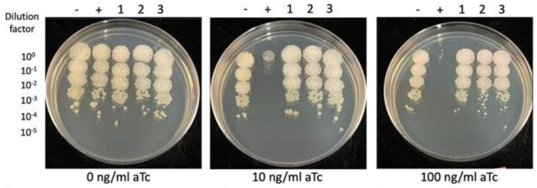


| Lane | Plasmid name | Gene name, replicate | Toxic/Non-toxic | Colony color on 100 ng/ml aTc plate* |
|---|---|---|---|---|
| - Non-toxic control | pExTra03 | Fruitloop *52* mutant | Non-toxic | + |
| + Toxic control | pExTra02 | Fruitloop *52* | Toxic | - |
| 1 | pExTra-NormanBulbieJr101 | NormanBulbieJr *101* replicate 1 | Non-toxic | + |
| 2 | pExTra-NormanBulbieJr101 | NormanBulbieJr *101* replicate 2 | Non-toxic | + |
| 3 | pExTra-NormanBulbieJr101 | NormanBulbieJr *101* replicate 3 | Non-toxic | + |

*Key: NG (no growth) - (no pink color) +(faint pink color) ++(obvious pink color) +++ (dark pink color)

Gene *98*; Score *0*

Images taken after 3 days at 37 ºC on 7H11 agar

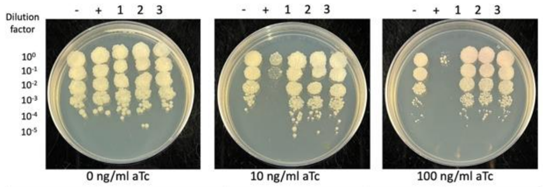


| Lane | Plasmid name | Gene name, replicate | Toxic/Non-toxic | Colony color on 100 ng/ml aTc plate* |
|---|---|---|---|---|
| - Non-toxic control | pExTra03 | Fruitloop *52* mutant | Non-toxic | + |
| + Toxic control | pExTra02 | Fruitloop *52* | Toxic | - |
| 1 | pExTra-NormanBulbieJr98 | NormanBulbieJr *98* replicate 1 | Non-toxic | + |
| 2 | pExTra-NormanBulbieJr98 | NormanBulbieJr *98* replicate 2 | Non-toxic | + |
| 3 | pExTra-NormanBulbieJr98 | NormanBulbieJr *98* replicate 3 | Non-toxic | + |

*Key: NG (no growth) - (no pink color) +(faint pink color) ++(obvious pink color) +++ (dark pink color)

Gene *99*; Score *2*

Images taken after 4 days at 37 ºC on 7H11 agar

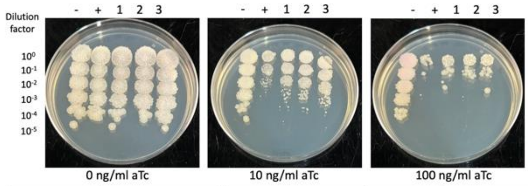


| Lane | Plasmid name | Gene name, replicate | Toxic/Non-toxic | Colony color on 100 ng/ml aTc plate* |
|---|---|---|---|---|
| - Non-toxic control | pExTra03 | Fruitloop *52* mutant | Non-toxic | + |
| + Toxic control | pExTra02 | Fruitloop *52* | Toxic | - |
| 1 | pExTra-NormanBulbieJr99 | NormanBulbieJr *99* replicate 1 | Toxic | - |
| 2 | pExTra-NormanBulbieJr99 | NormanBulbieJr *99* replicate 2 | Toxic | - |
| 3 | pExTra-NormanBulbieJr99 | NormanBulbieJr *99* replicate 3 | Toxic | - |

*Key: NG (no growth) - (no pink color) +(faint pink color) ++(obvious pink color) +++ (dark pink color)

Gene *100*; Score *0*

Images taken after 3 days at 37 ºC on 7H11 agar

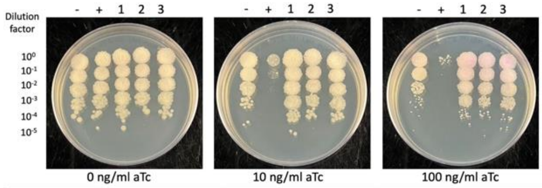


| Lane | Plasmid name | Gene name, replicate | Toxic/Non-toxic | Colony color on 100 ng/ml aTc plate* |
|---|---|---|---|---|
| - Non-toxic control | pExTra03 | Fruitloop *52* mutant | Non-toxic | - |
| + Toxic control | pExTra02 | Fruitloop *52* | Toxic | - |
| 1 | pExTra-NormanBulbieJr100 | NormanBulbieJr *100* replicate 1 | Non-toxic | ++ |
| 2 | pExTra-NormanBulbieJr100 | NormanBulbieJr *100* replicate 2 | Non-toxic | ++ |
| 3 | pExTra-NormanBulbieJr100 | NormanBulbieJr *100* replicate 3 | Non-toxic | ++ |

*Key: NG (no growth) - (no pink color) +(faint pink color) ++(obvious pink color) +++ (dark pink color)

### Supplemental Figure 2

Images taken after 2 days at 37 ºC

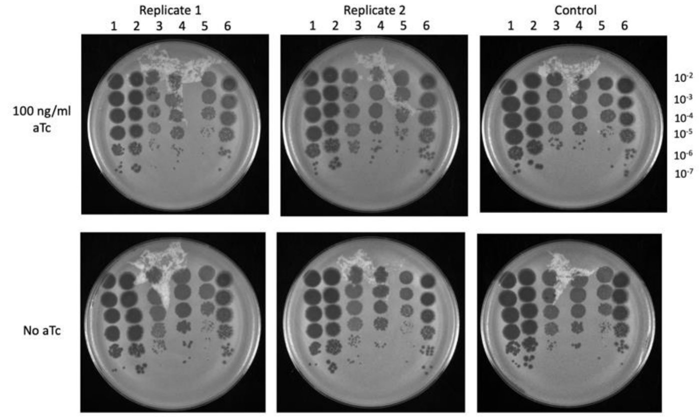


Table 1: *M. smegmatis* lawns

| | Plasmid name | Gene name |
|---|---|---|
| Replicates | pExTra-NormanBublieJr39 | NormanBublieJr *39* |
| Control | pExTra03 | *Fruitloop* 52-MUT |

Table 2: Phage lysates

| | 1 | 2 | 3 | 4 | 5 | 6 |
|---|---|---|---|---|---|---|
| Phage name | NormanBublieJr | Girr | Akhila | Zapner | Squirty | Tchen |
| Titer on control plate* | ~1.33x $10^{10}$ pfu/ml | ~1.0 x $10^{10}$ pfu/ml | ~4.0 x $10^{9}$ pfu/ml | ~1.67 x $10^{9}$ pfu/ml | ~4.0 x $10^{8}$ pfu/ml | ~2.67 x $10^{9}$ pfu/ml |
| E.O.P (replicate 1) | ~1 | ~1 | ~1 | ~1 | ~1 | ~1 |
| E.O.P (replicate 2) | ~1 | ~1 | ~1 | ~1 | ~1 | ~1 |

* The 100 ng/ml aTc plate results were used to calculate E.O.P.

E.O.P.= replicate plate titer/ control plate titer

Images taken after 2 days at 37 ºC

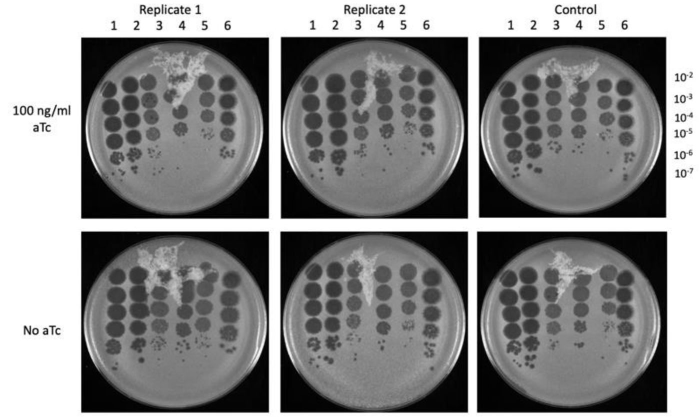


Table 1: *M. smegmatis* lawns

| | Plasmid name | Gene name |
|---|---|---|
| Replicates | pExTra-NormanBublieJr40 | NormanBublieJr *40* |
| Control | pExTra03 | *Fruitloop* 52-MUT |

Table 2: Phage lysates

| | 1 | 2 | 3 | 4 | 5 | 6 |
|---|---|---|---|---|---|---|
| Phage name | NormanBublieJr | Girr | Akhila | Zapner | Squirty | Tchen |
| Titer on control plate* | ~1.33x $10^{10}$ pfu/ml | ~1.0 x $10^{10}$ pfu/ml | ~4.0 x $10^{9}$ pfu/ml | ~1.67 x $10^{9}$ pfu/ml | ~4.0 x $10^{8}$ pfu/ml | ~2.67 x $10^{9}$ pfu/ml |
| E.O.P (replicate 1) | ~1 | ~1 | ~1 | ~1 | ~1 | ~1 |
| E.O.P (replicate 2) | ~1 | ~1 | ~1 | ~1 | ~1 | ~1 |

* The 100 ng/ml aTc plate results were used to calculate E.O.P.

E.O.P.= replicate plate titer/ control plate titer

Images taken after 2 days at 37 ºC

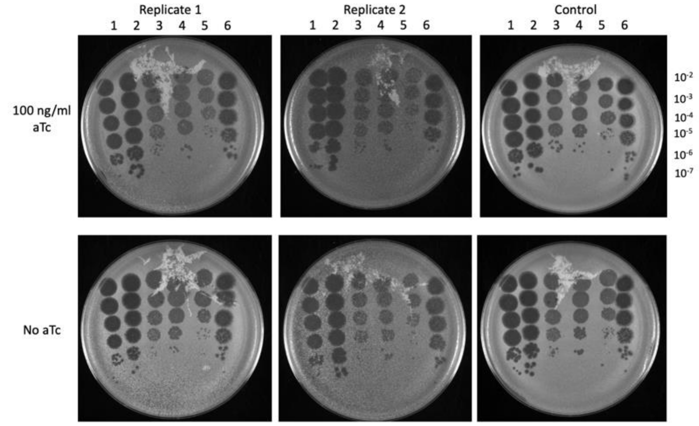


Table 1: *M. smegmatis* lawns

| | Plasmid name | Gene name |
|---|---|---|
| Replicates | pExTra-NormanBublieJr41 | NormanBublieJr *41* |
| Control | pExTra03 | *Fruitloop* 52-MUT |

Table 2: Phage lysates

| | 1 | 2 | 3 | 4 | 5 | 6 |
|---|---|---|---|---|---|---|
| Phage name | NormanBublieJr | Girr | Akhila | Zapner | Squirty | Tchen |
| Titer on control plate* | ~1.33x $10^{10}$ pfu/ml | ~1.0 x $10^{10}$ pfu/ml | ~4.0 x $10^{9}$ pfu/ml | ~1.67 x $10^{9}$ pfu/ml | ~4.0 x $10^{8}$ pfu/ml | ~2.67 x $10^{9}$ pfu/ml |
| E.O.P (replicate 1) | ~1 | ~1 | ~1 | ~1 | ~1 | ~1 |
| E.O.P (replicate 2) | ~1 | ~1 | ~1 | ~1 | ~1 | ~1 |

* The 100 ng/ml aTc plate results were used to calculate E.O.P.

E.O.P.= replicate plate titer/ control plate titer

### Figure S2

Images taken after 2 days at 37 ºC

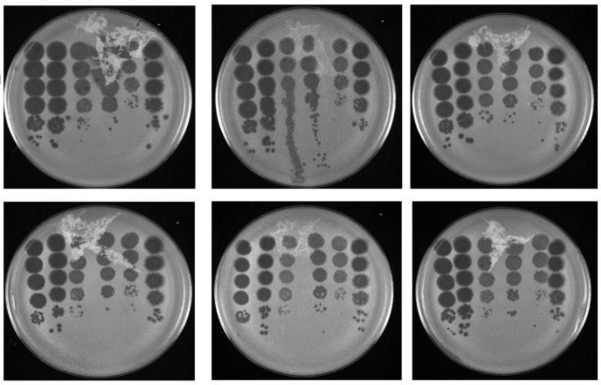


Table 1: *M. smegmatis* lawns

| | Plasmid name | Gene name |
|---|---|---|
| Replicates | pExTra-NormanBublieJr42 | NormanBublieJr *42* |
| Control | pExTra03 | *Fruitloop* 52-MUT |

Table 2: Phage lysates

| | 1 | 2 | 3 | 4 | 5 | 6 |
|---|---|---|---|---|---|---|
| Phage name | NormanBublieJr | Girr | Akhila | Zapner | Squirty | Tchen |
| Titer on control plate* | ~1.33x $10^{10}$ pfu/ml | ~1.0 x $10^{10}$ pfu/ml | ~4.0 x $10^{9}$ pfu/ml | ~1.67 x $10^{9}$ pfu/ml | ~4.0 x $10^{8}$ pfu/ml | ~2.67 x $10^{9}$ pfu/ml |
| E.O.P (replicate 1) | ~1 | ~1 | ~1 | ~1 | ~1 | ~1 |
| E.O.P (replicate 2) | ~1 | ~1 | ~1 | ~1 | ~1 | ~1 |

* The 100 ng/ml aTc plate results were used to calculate E.O.P.

E.O.P.= replicate plate titer/ control plate titer

Images taken after 2 days at 37 ºC

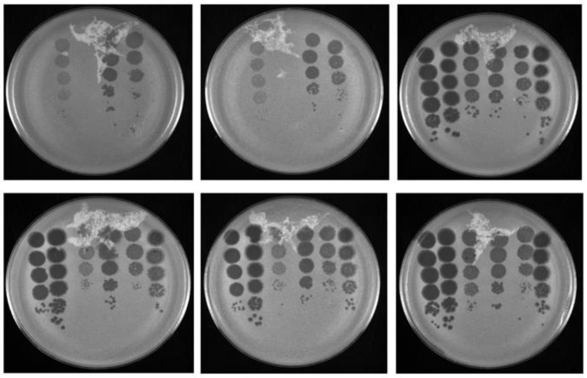


Table 1: *M. smegmatis* lawns

| | Plasmid name | Gene name |
|---|---|---|
| Replicates | pExTra-NormanBublieJr45 | NormanBublieJr *45* |
| Control | pExTra03 | *Fruitloop* 52-MUT |

Table 2: Phage lysates

| | 1 | 2 | 3 | 4 | 5 | 6 |
|---|---|---|---|---|---|---|
| Phage name | NormanBublieJr | Girr | Akhila | Zapner | Squirty | Tchen |
| Titer on control plate* | ~1.33x $10^{10}$ pfu/ml | ~1.0 x $10^{10}$ pfu/ml | ~4.0 x $10^{9}$ pfu/ml | ~1.67 x $10^{9}$ pfu/ml | ~4.0 x $10^{8}$ pfu/ml | ~2.67 x $10^{9}$ pfu/ml |
| E.O.P (replicate 1) | 0 | 0.367 | 0 | 0 | 6.67 | 0.25 |
| E.O.P (replicate 2) | 0 | 0.333 | 0 | 0 | 4.17 | 0.125 |

* The 100 ng/ml aTc plate results were used to calculate E.O.P.

E.O.P.= replicate plate titer/ control plate titer

Images taken after 2 days at 37 ºC

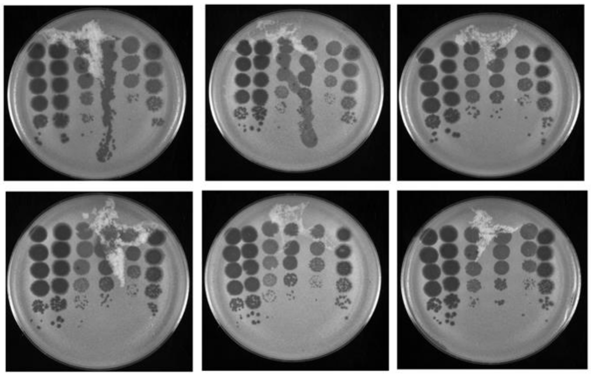


Table 1: *M. smegmatis* lawns

| | Plasmid name | Gene name |
|---|---|---|
| Replicates | pExTra-NormanBublieJr46 | NormanBublieJr *46* |
| Control | pExTra03 | *Fruitloop* 52-MUT |

Table 2: Phage lysates

| | 1 | 2 | 3 | 4 | 5 | 6 |
|---|---|---|---|---|---|---|
| Phage name | NormanBublieJr | Girr | Akhila | Zapner | Squirty | Tchen |
| Titer on control plate* | ~1.33x $10^{10}$ pfu/ml | ~1.0 x $10^{10}$ pfu/ml | ~4.0 x $10^{9}$ pfu/ml | ~1.67 x $10^{9}$ pfu/ml | ~4.0 x $10^{8}$ pfu/ml | ~2.67 x $10^{9}$ pfu/ml |
| E.O.P (replicate 1) | ~1 | ~1 | ~1 | ~1 | ~1 | ~1 |
| E.O.P (replicate 2) | ~1 | ~1 | ~1 | ~1 | ~1 | ~1 |

* The 100 ng/ml aTc plate results were used to calculate E.O.P.

E.O.P.= replicate plate titer/ control plate titer

Figure S2

Images taken after 2 days at 37 ºC

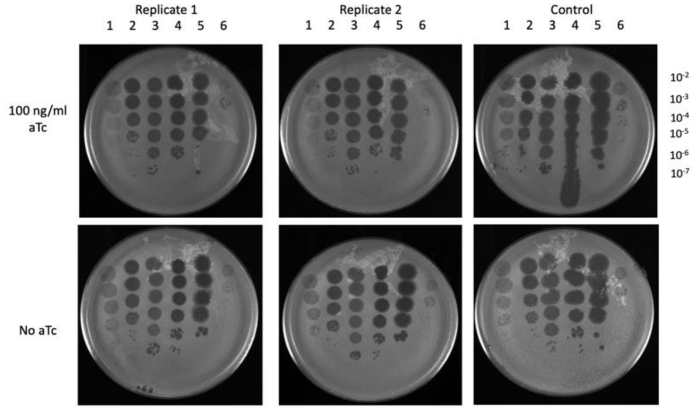


Table 1: *M. smegmatis* lawns

| | Plasmid name | Gene name |
|---|---|---|
| Replicates | pExTra-NormanBublieJr39 | NormanBublieJr *39* |
| Control | pExTra03 | *Fruitloop* 52-MUT |

Table 2: Phage lysates

| | 7 | 8 | 9 | 10 | 11 | 12 |
|---|---|---|---|---|---|---|
| Phage name | ZoeJ | Charlie | Tortellini | Lebron | Bxb1 | Typha |
| Titer on control plate* | ~$1.67 \times 10^{10}$ pfu/ml | ~$1.33 \times 10^{10}$ pfu/ml | ~$3.0 \times 10^{10}$ pfu/ml | ~$1.0 \times 10^{10}$ pfu/ml | ~$3.33 \times 10^{9}$ pfu/ml | ~$2.33 \times 10^{7}$ pfu/ml |
| E.O.P (replicate 1) | ~1 | ~1 | ~1 | ~1 | ~1 | ~1 |
| E.O.P (replicate 2) | ~1 | ~1 | ~1 | ~1 | ~1 | ~1 |

* The 100 ng/ml aTc plate results were used to calculate E.O.P.

E.O.P.= replicate plate titer/ control plate titer

Images taken after 2 days at 37 ºC

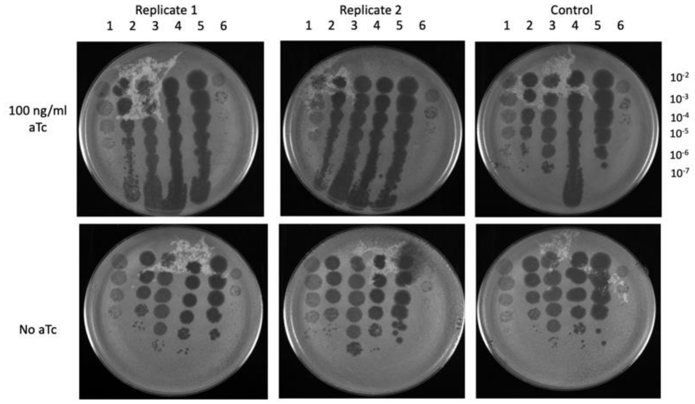


Table 1: *M. smegmatis* lawns

| | Plasmid name | Gene name |
|---|---|---|
| Replicates | pExTra-NormanBublieJr40 | NormanBublieJr *40* |
| Control | pExTra03 | *Fruitloop* 52-MUT |

Table 2: Phage lysates

| | 7 | 8 | 9 | 10 | 11 | 12 |
|---|---|---|---|---|---|---|
| Phage name | ZoeJ | Charlie | Tortellini | Lebron | Bxb1 | Typha |
| Titer on control plate* | ~$1.67 \times 10^{10}$ pfu/ml | ~$1.33 \times 10^{10}$ pfu/ml | ~$3.0 \times 10^{10}$ pfu/ml | ~$1.0 \times 10^{10}$ pfu/ml | ~$3.33 \times 10^{9}$ pfu/ml | ~$2.33 \times 10^{7}$ pfu/ml |
| E.O.P (replicate 1) | ~1 | ~1 | ~1 | ~1 | ~1 | ~1 |
| E.O.P (replicate 2) | ~1 | ~1 | ~1 | ~1 | ~1 | ~1 |

* The 100 ng/ml aTc plate results were used to calculate E.O.P.

E.O.P.= replicate plate titer/ control plate titer

Images taken after 2 days at 37 ºC

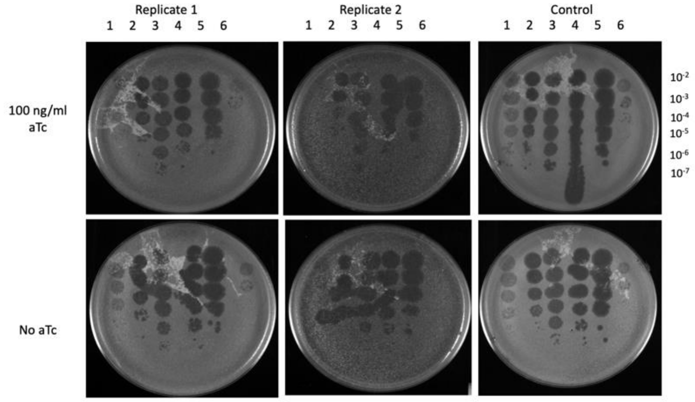


Table 1: *M. smegmatis* lawns

| | Plasmid name | Gene name |
|---|---|---|
| Replicates | pExTra-NormanBublieJr41 | NormanBublieJr *41* |
| Control | pExTra03 | *Fruitloop* 52-MUT |

Table 2: Phage lysates

| | 7 | 8 | 9 | 10 | 11 | 12 |
|---|---|---|---|---|---|---|
| Phage name | ZoeJ | Charlie | Tortellini | Lebron | Bxb1 | Typha |
| Titer on control plate* | ~$1.67 \times 10^{10}$ pfu/ml | ~$1.33 \times 10^{10}$ pfu/ml | ~$3.0 \times 10^{10}$ pfu/ml | ~$1.0 \times 10^{10}$ pfu/ml | ~$3.33 \times 10^{9}$ pfu/ml | ~$2.33 \times 10^{7}$ pfu/ml |
| E.O.P (replicate 1) | ~1 | ~1 | ~1 | ~1 | ~1 | ~1 |
| E.O.P (replicate 2) | ~1 | ~1 | ~1 | ~1 | ~1 | ~1 |

* The 100 ng/ml aTc plate results were used to calculate E.O.P.

E.O.P.= replicate plate titer/ control plate titer

### Figure S2

Images taken after 2 days at 37 ºC

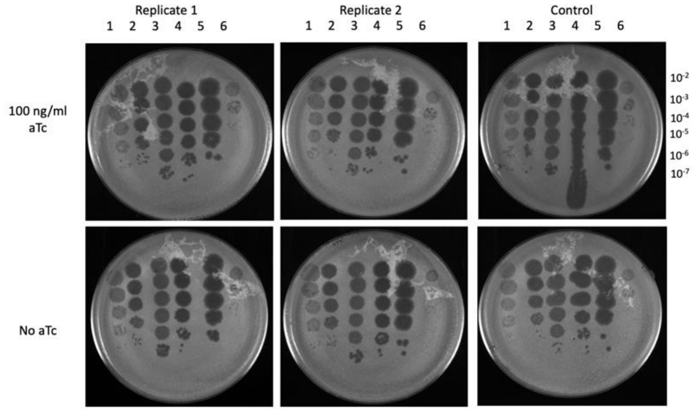


Table 1: *M. smegmatis* lawns

| | Plasmid name | Gene name |
|---|---|---|
| Replicates | pExTra-NormanBublieJr42 | NormanBublieJr *42* |
| Control | pExTra03 | *Fruitloop* 52-MUT |

Table 2: Phage lysates

| | 7 | 8 | 9 | 10 | 11 | 12 |
|---|---|---|---|---|---|---|
| Phage name | ZoeJ | Charlie | Tortellini | Lebron | Bxb1 | Typha |
| Titer on control plate* | ~1.67x $10^{10}$ pfu/ml | ~1.33 x $10^{10}$ pfu/ml | ~3.0 x $10^{10}$ pfu/ml | ~1.0 x $10^{10}$ pfu/ml | ~3.33 x $10^{9}$ pfu/ml | ~2.33 x $10^{7}$ pfu/ml |
| E.O.P (replicate 1) | ~1 | ~1 | ~1 | ~1 | ~1 | ~1 |
| E.O.P (replicate 2) | ~1 | ~1 | ~1 | ~1 | ~1 | ~1 |

* The 100 ng/ml aTc plate results were used to calculate E.O.P.

E.O.P.= replicate plate titer/ control plate titer

Images taken after 2 days at 37 ºC

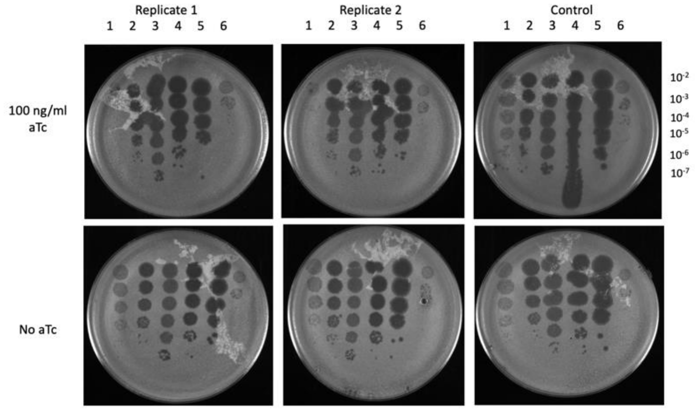


Table 1: *M. smegmatis* lawns

| | Plasmid name | Gene name |
|---|---|---|
| Replicates | pExTra-NormanBublieJr45 | NormanBublieJr *45* |
| Control | pExTra03 | *Fruitloop* 52-MUT |

Table 2: Phage lysates

| | 7 | 8 | 9 | 10 | 11 | 12 |
|---|---|---|---|---|---|---|
| Phage name | ZoeJ | Charlie | Tortellini | Lebron | Bxb1 | Typha |
| Titer on control plate* | ~1.67x $10^{10}$ pfu/ml | ~1.33 x $10^{10}$ pfu/ml | ~3.0 x $10^{10}$ pfu/ml | ~1.0 x $10^{10}$ pfu/ml | ~3.33 x $10^{9}$ pfu/ml | ~2.33 x $10^{7}$ pfu/ml |
| E.O.P (replicate 1) | ~4.0x $10^{-6}$ pfu/ml | ~1 | ~1 | ~1 | ~1 | ~1 |
| E.O.P (replicate 2) | ~4.0x $10^{-6}$ pfu/ml | ~1 | ~1 | ~1 | ~1 | ~1 |

* The 100 ng/ml aTc plate results were used to calculate E.O.P.

E.O.P.= replicate plate titer/ control plate titer

Images taken after 2 days at 37 ºC

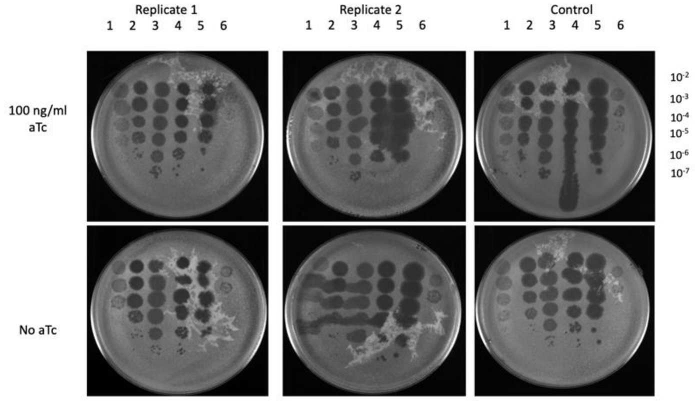


Table 1: *M. smegmatis* lawns

| | Plasmid name | Gene name |
|---|---|---|
| Replicates | pExTra-NormanBublieJr46 | NormanBublieJr *46* |
| Control | pExTra03 | *Fruitloop* 52-MUT |

Table 2: Phage lysates

| | 7 | 8 | 9 | 10 | 11 | 12 |
|---|---|---|---|---|---|---|
| Phage name | ZoeJ | Charlie | Tortellini | Lebron | Bxb1 | Typha |
| Titer on control plate* | ~1.67x $10^{10}$ pfu/ml | ~1.33 x $10^{10}$ pfu/ml | ~3.0 x $10^{10}$ pfu/ml | ~1.0 x $10^{10}$ pfu/ml | ~3.33 x $10^{9}$ pfu/ml | ~2.33 x $10^{7}$ pfu/ml |
| E.O.P (replicate 1) | ~1 | ~1 | ~1 | ~1 | ~1 | ~1 |
| E.O.P (replicate 2) | ~1 | ~1 | ~1 | ~1 | ~1 | ~1 |

* The 100 ng/ml aTc plate results were used to calculate E.O.P.

E.O.P.= replicate plate titer/ control plate titer

**Supplemental Figure 3**

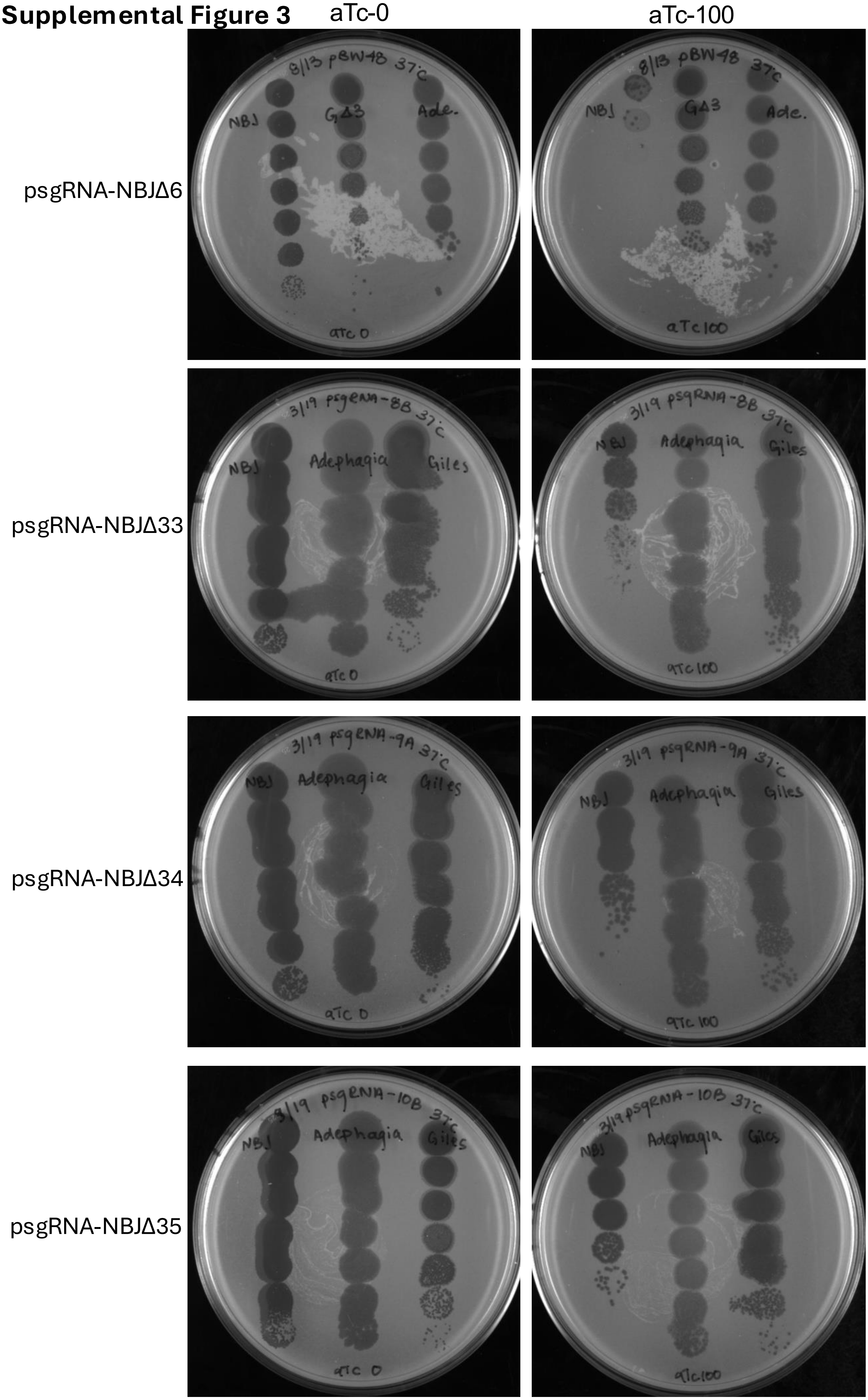

Figure S3

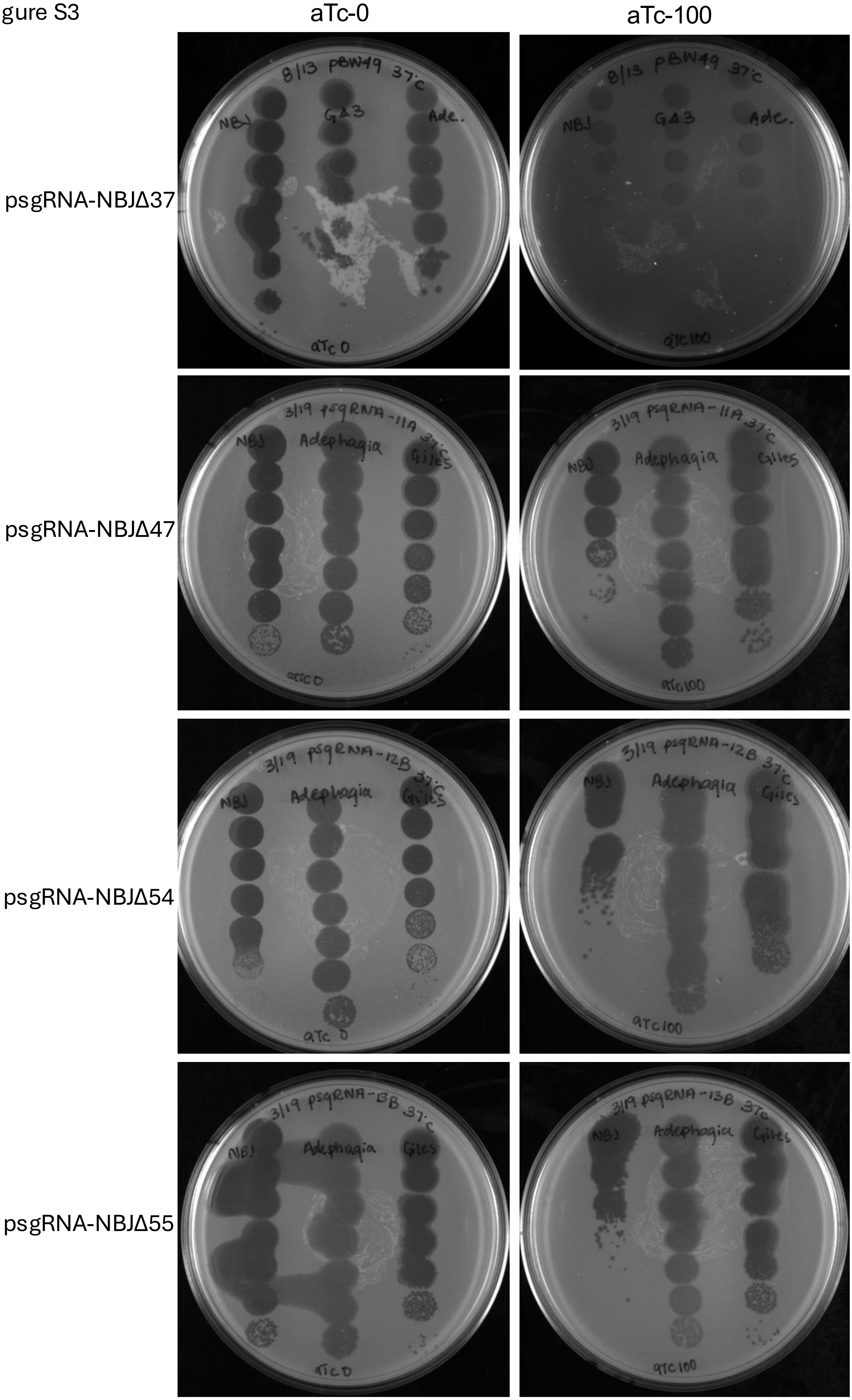
aTc-0
aTc-100
psgRNA-NBJΔ37
psgRNA-NBJΔ47
psgRNA-NBJΔ54
psgRNA-NBJΔ55

Figure S3

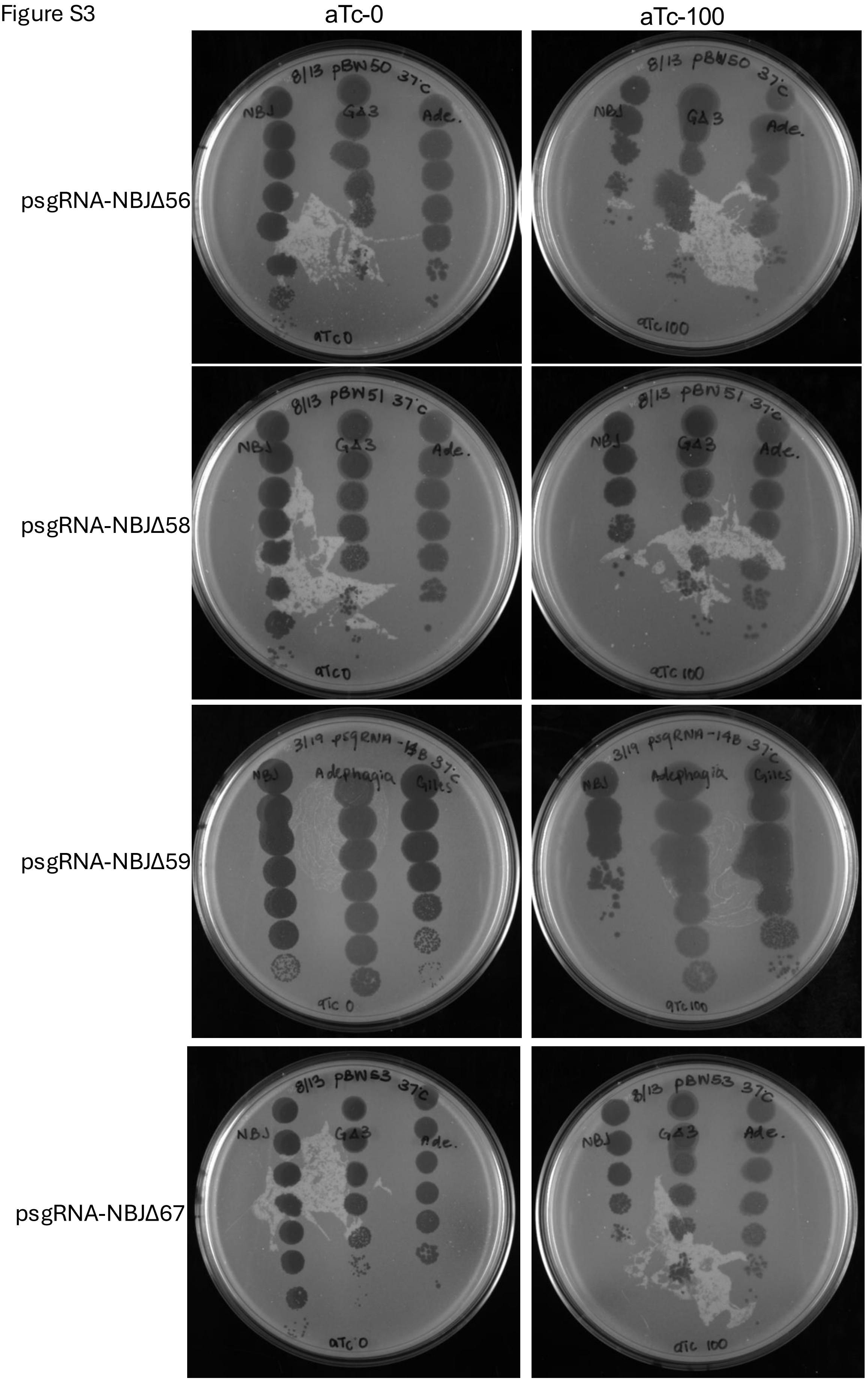
aTc-0
aTc-100
psgRNA-NBJΔ56
psgRNA-NBJΔ58
psgRNA-NBJΔ59
psgRNA-NBJΔ67

Figure S3

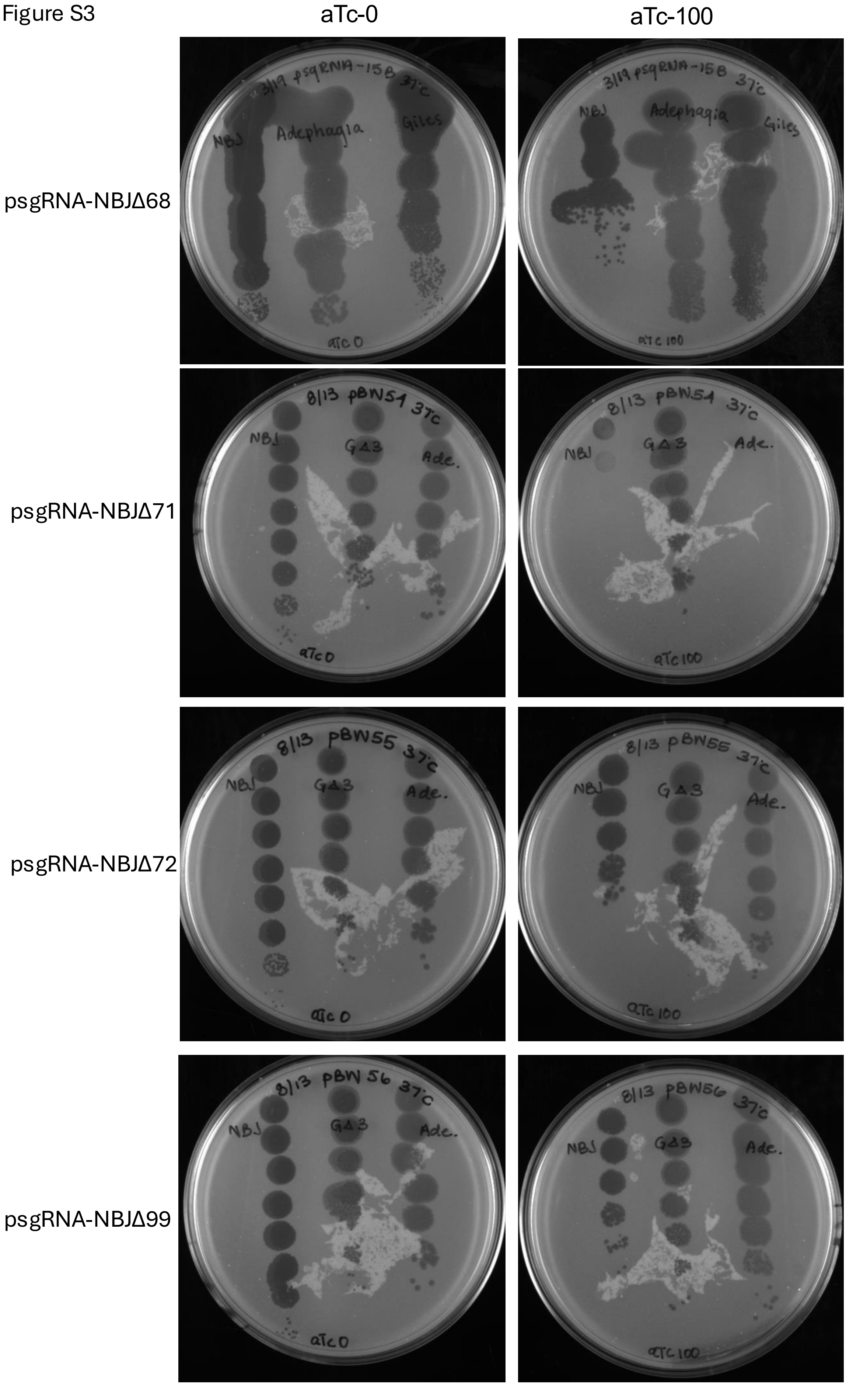
aTc-0
aTc-100
psgRNA-NBJΔ68
psgRNA-NBJΔ71
psgRNA-NBJΔ72
psgRNA-NBJΔ99
3/19 psgRNA-15B 37°C
NBJ
Adephagia
Giles
aTc 0
aTc 100
8/13 pBW54 37°C
GΔ3
Ade.
8/13 pBW55 37°C
8/13 pBW56 37°C

Figure S3

aTc-0 | aTc-100

psgRNA-NBJΔ101

psgRNA-NBJΔ45

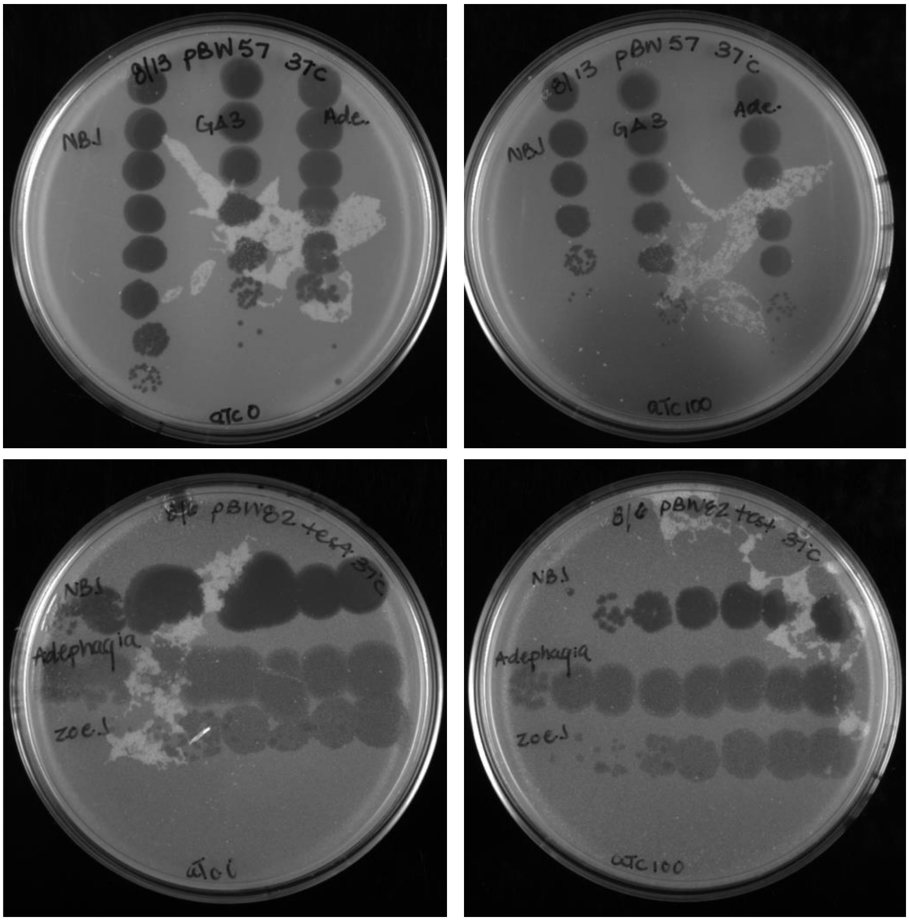

**Supplemental Figure 4**

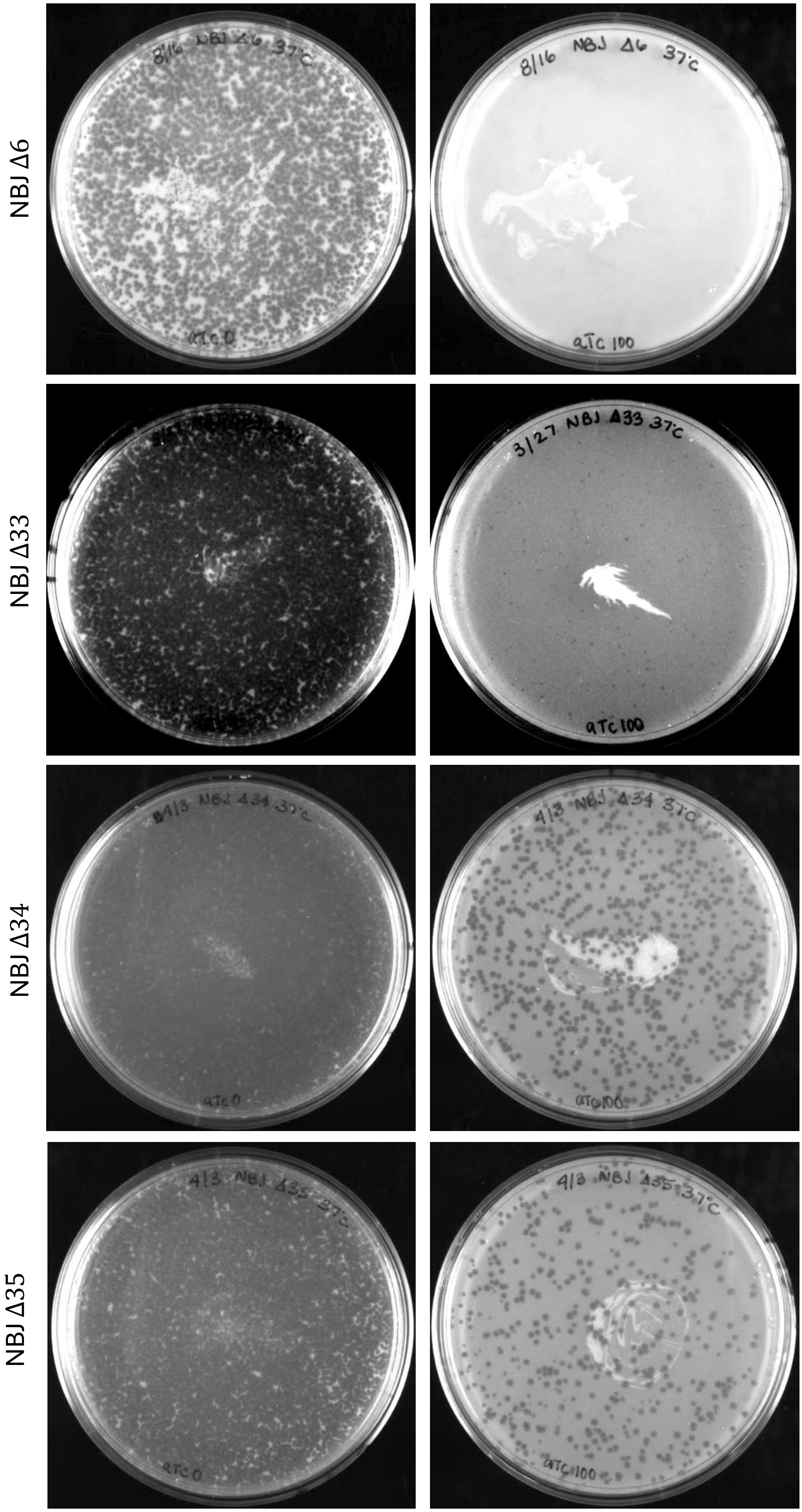

Figure S4

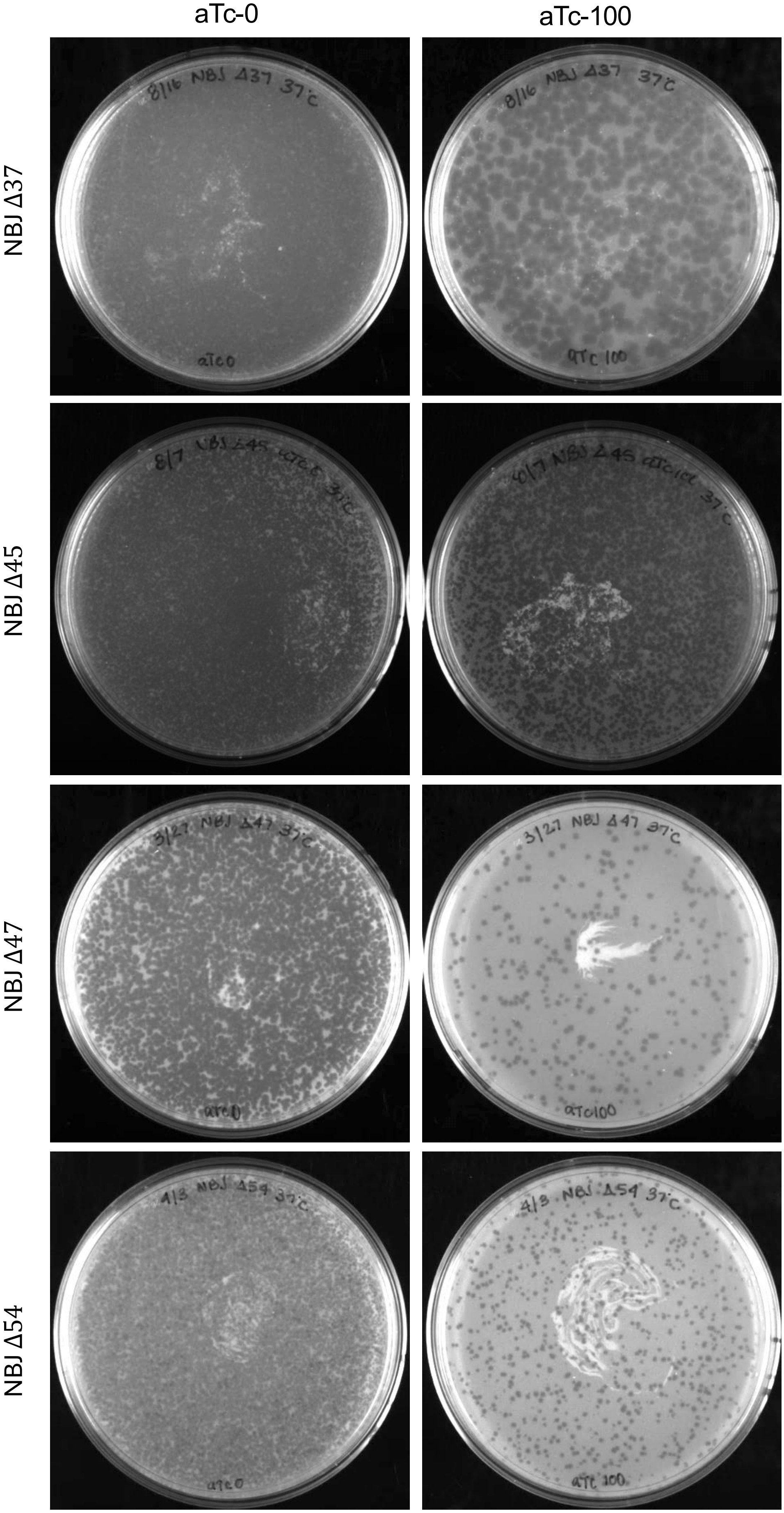

Figure S4

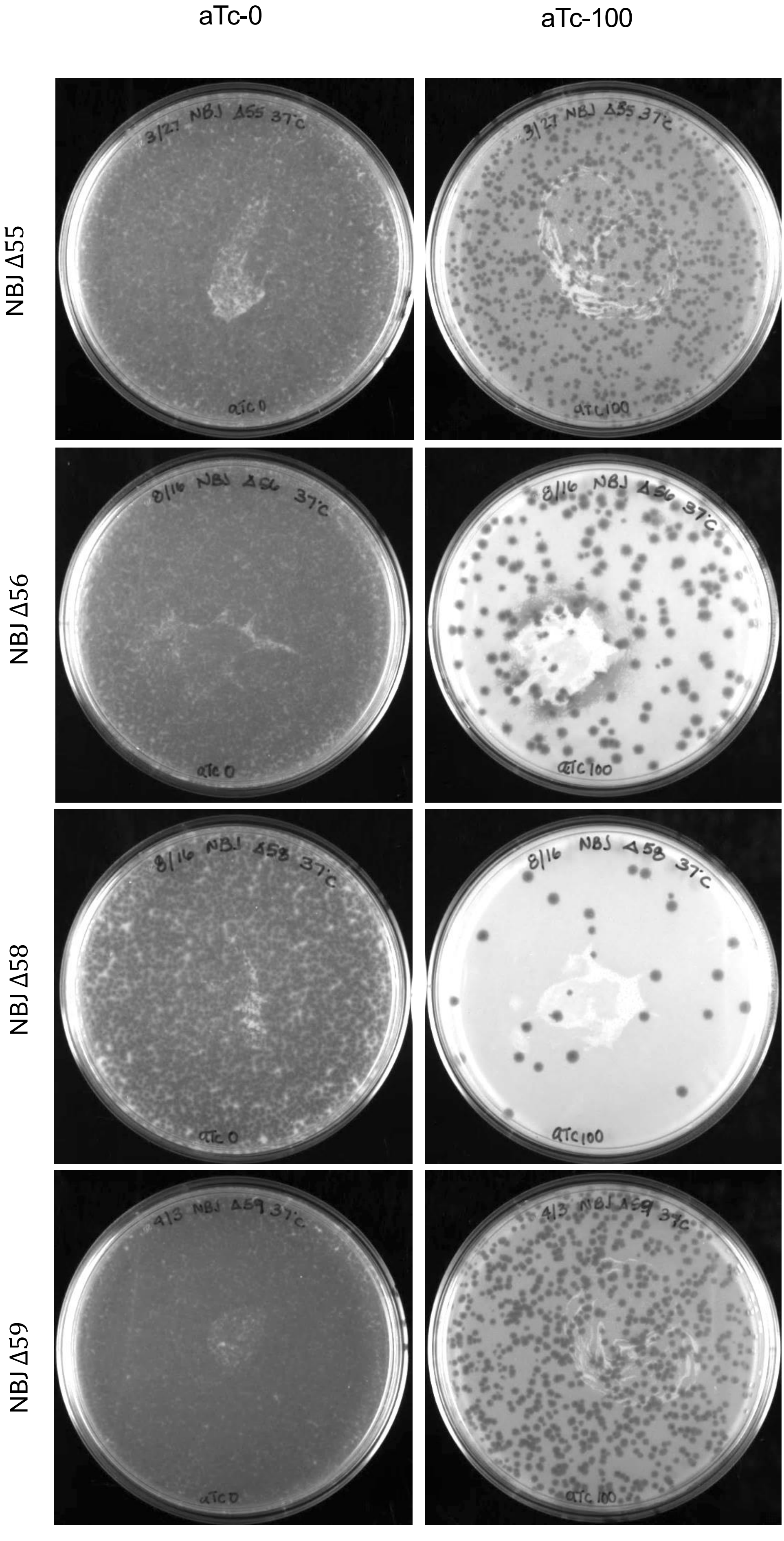

Figure S4

aTc-0 aTc-100

NBJ Δ67

NBJ Δ68

NBJ Δ71

NBJ Δ72

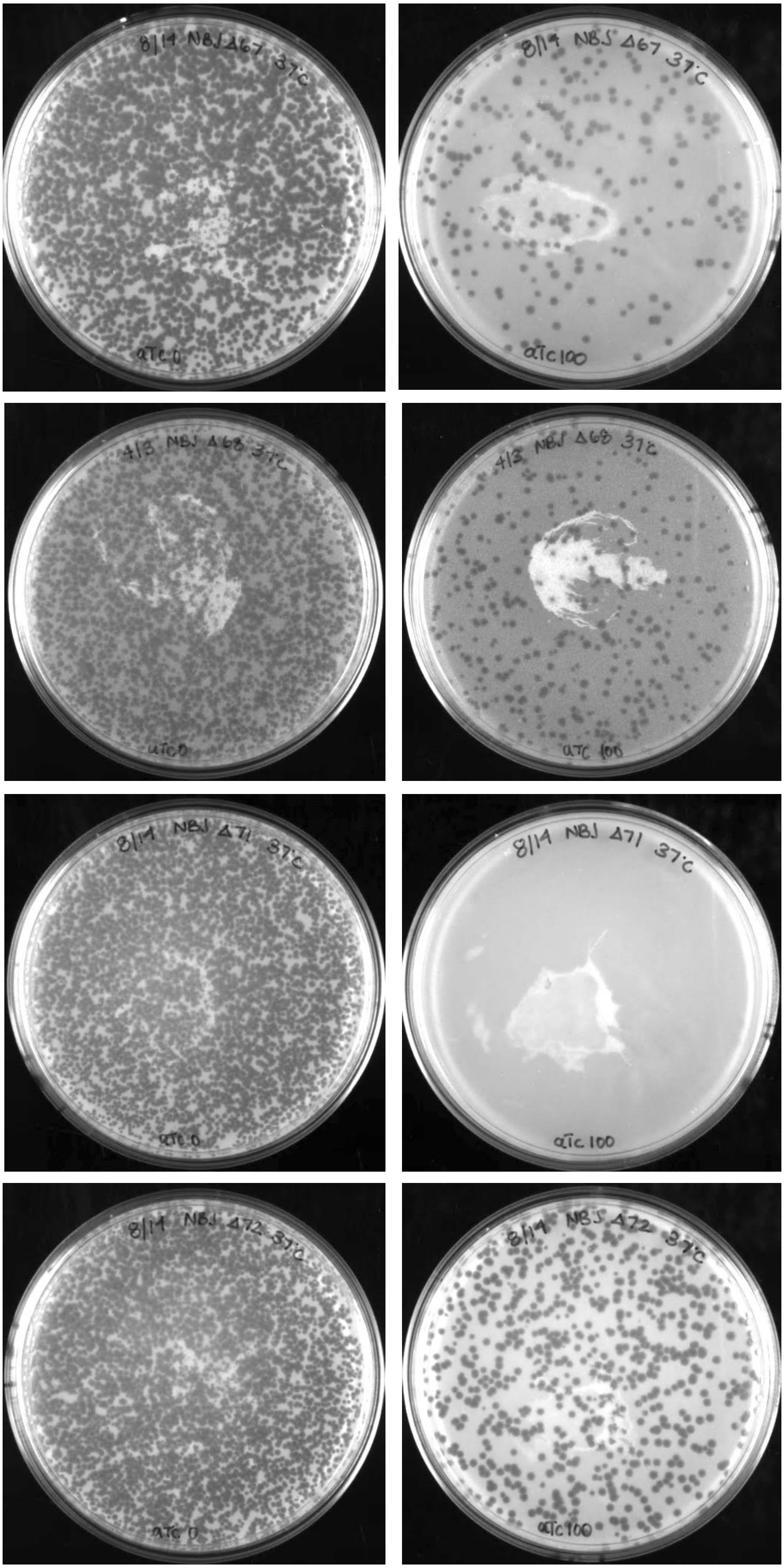

Figure S4

aTc-0 aTc-100

NBJ Δ99

NBJ Δ101

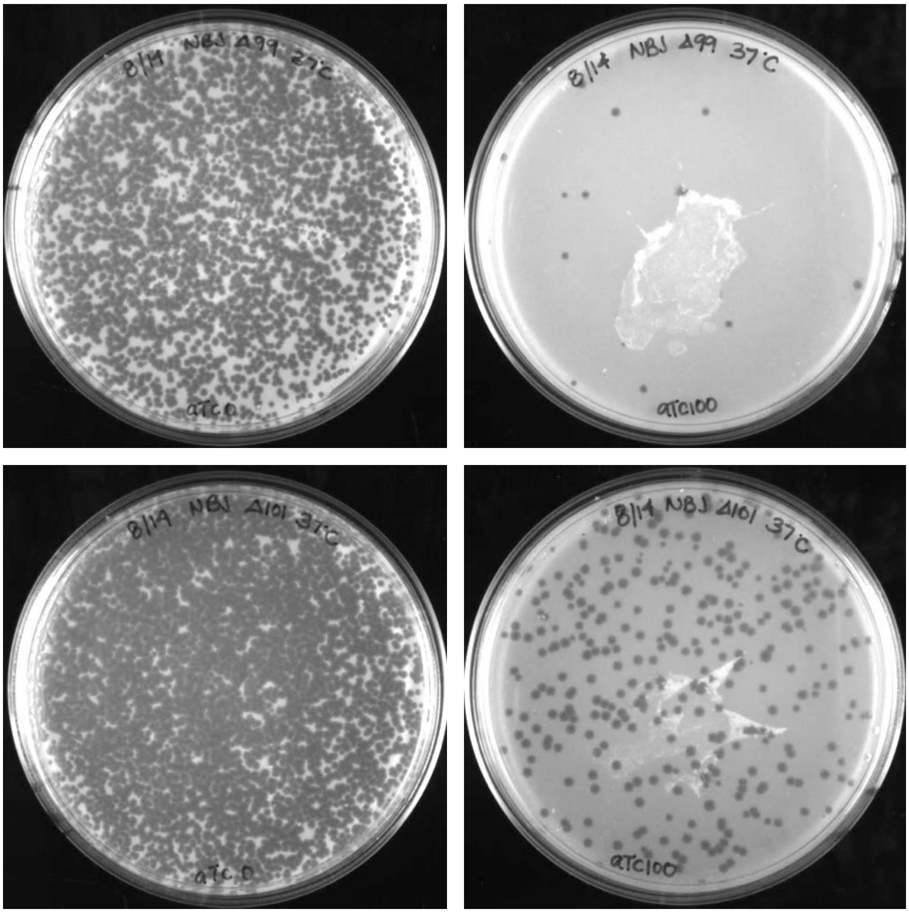

**Supplemental Figure 5**

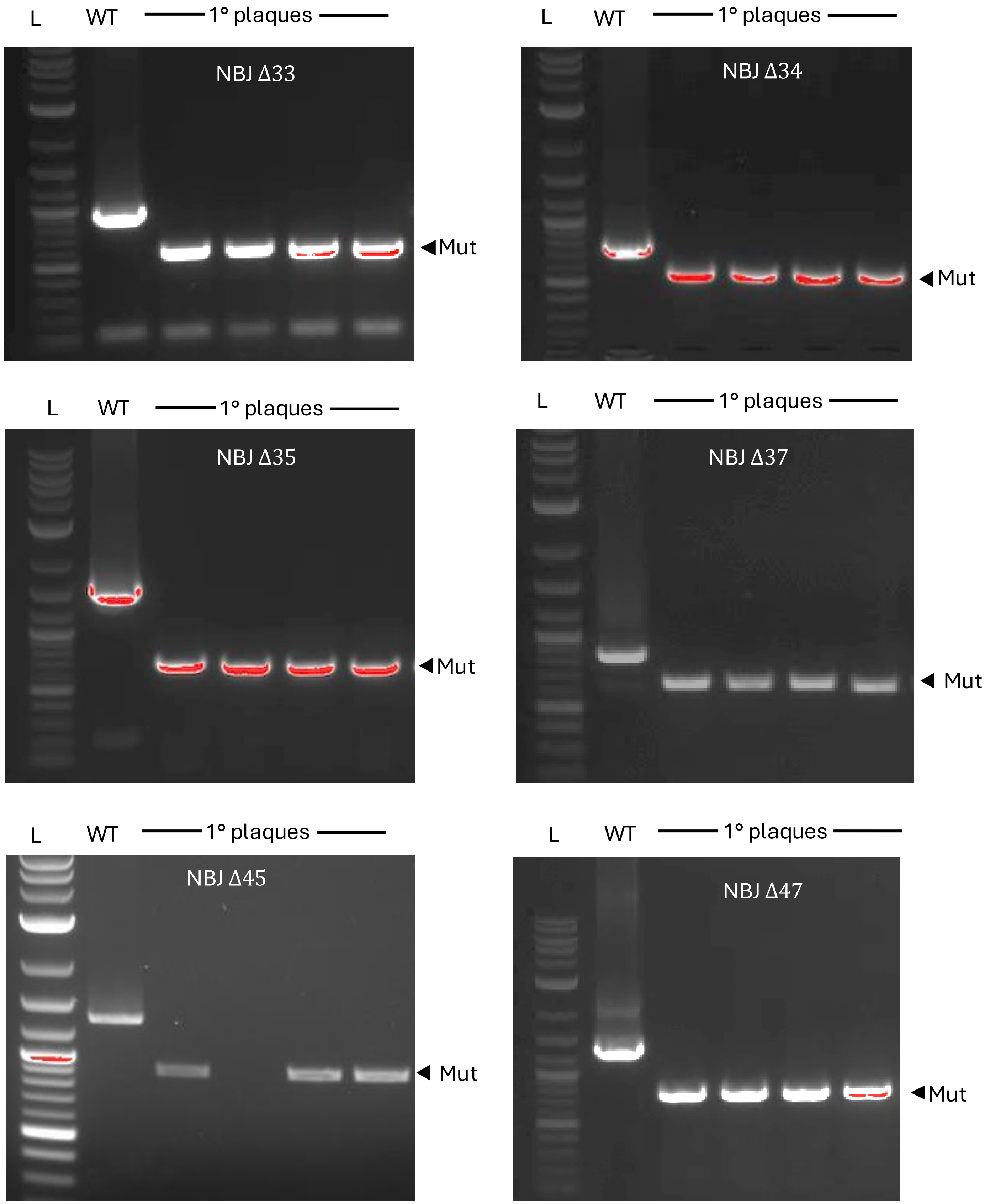

Figure S5

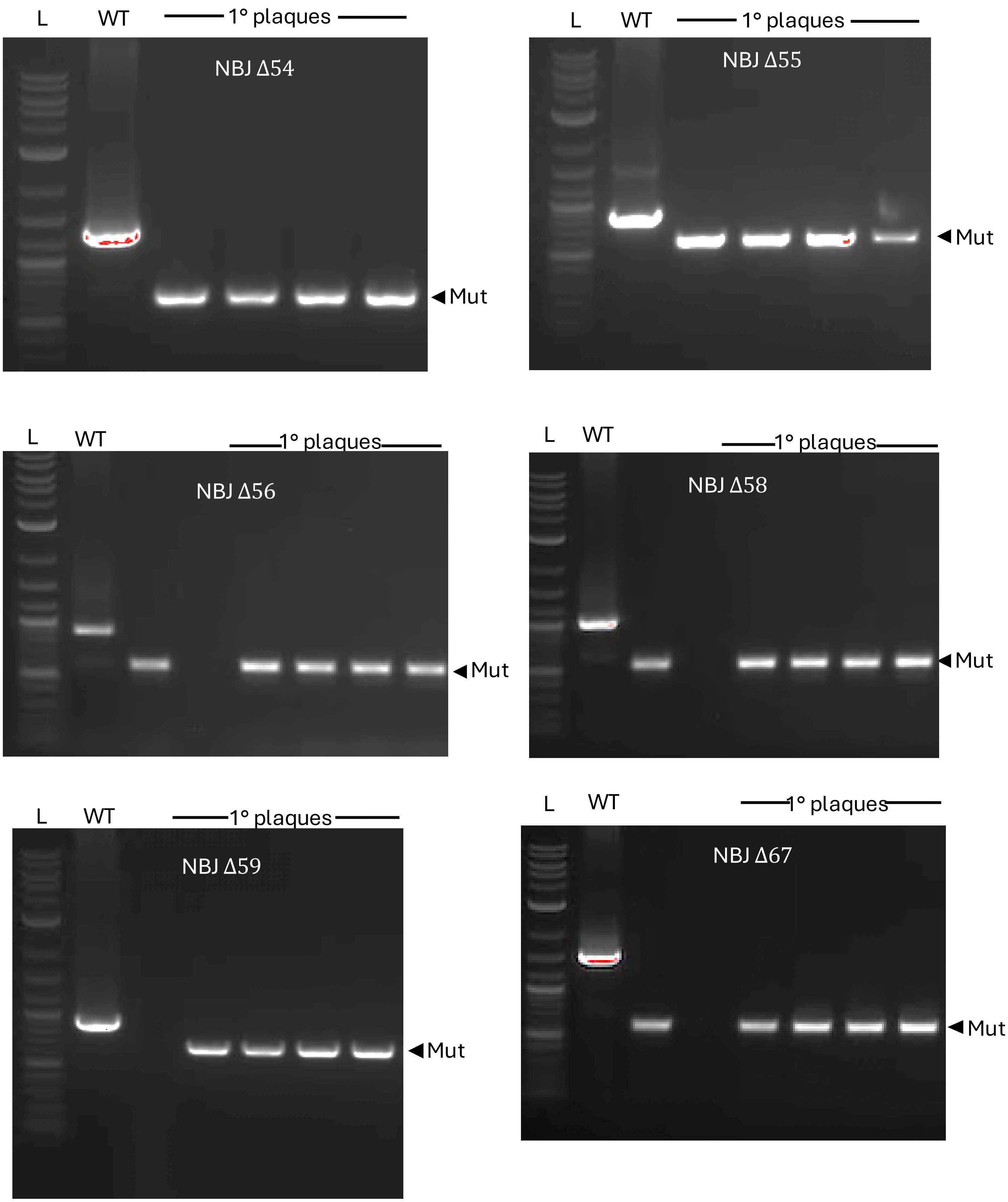
L
WT
1° plaques
NBJ Δ54
◀Mut
L
WT
1° plaques
NBJ Δ55
◀Mut
L
WT
1° plaques
NBJ Δ56
◀Mut
L
WT
1° plaques
NBJ Δ58
◀Mut
L
WT
1° plaques
NBJ Δ59
◀Mut
L
WT
1° plaques
NBJ Δ67
◀Mut

Figure S5

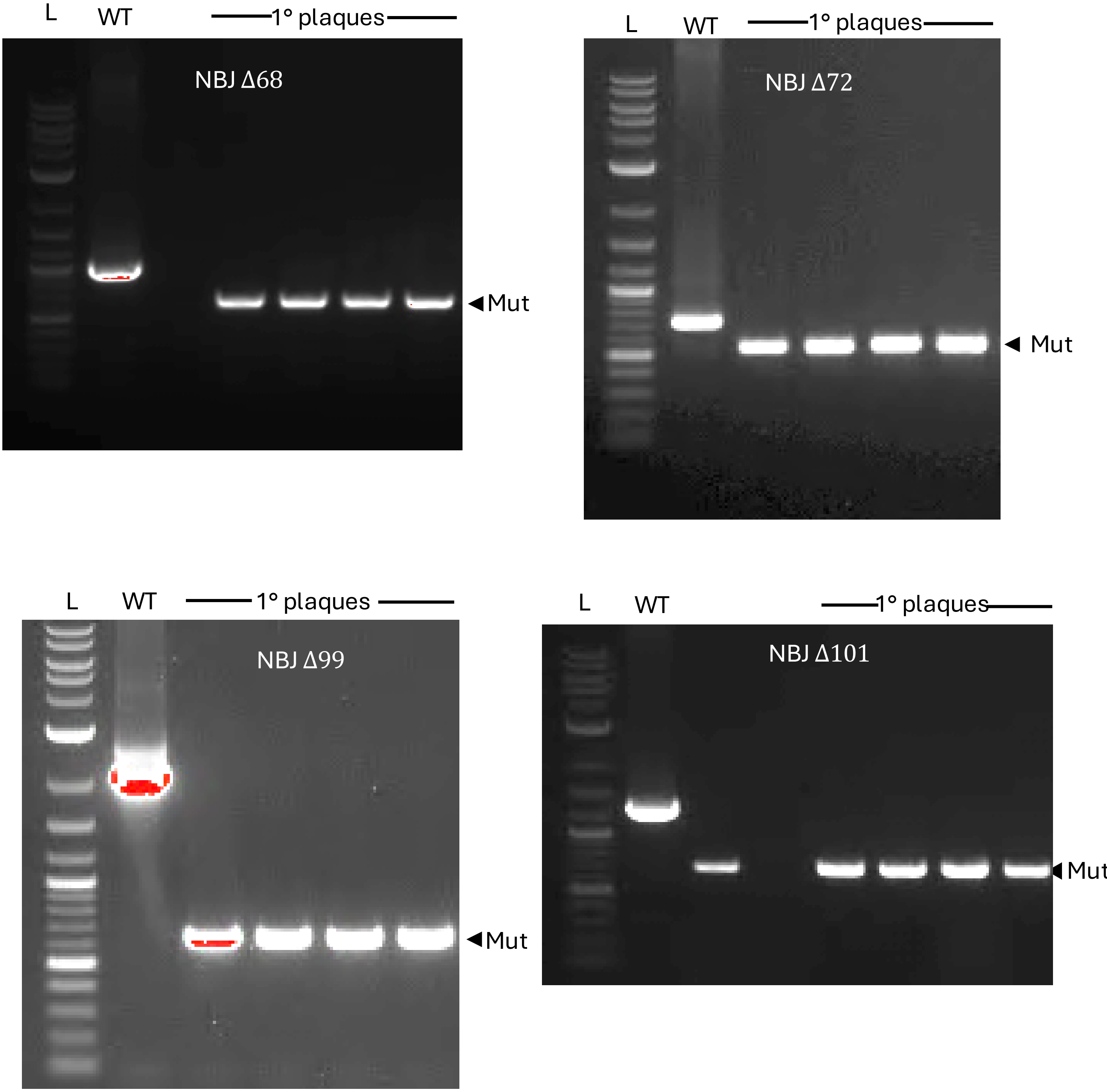
L
WT
1° plaques
NBJ Δ68
Mut
L
WT
1° plaques
NBJ Δ72
Mut
L
WT
1° plaques
NBJ Δ99
Mut
L
WT
1° plaques
NBJ Δ101
Mut

**Table S1: Oligonucleotides used in this study**

| OLIGO NAME | OLIGO SEQUENCE (5' TO 3') |
|---|---|
| ONORMANBULBIEJR1_F | atgcggaggaatcacttccatATGCCACCTGTACCTAAAG |
| ONORMANBULBIEJR1_R | tgcaggatccgactcgagtgtcgacTCAGGTCACAAGCTTCAAAC |
| ONORMANBULBIEJR2_F | atgcggaggaatcacttccatATGGCTGTTTTGCAGGTCC |
| ONORMANBULBIEJR2_R | tgcaggatccgactcgagtgtcgacTCAGTAGATCCGTCTAGGC |
| ONORMANBULBIEJR3_F | atgcggaggaatcacttccatATGACTGCTTCAACGCCAG |
| ONORMANBULBIEJR3_R | tgcaggatccgactcgagtgtcgacTCAGCGTGATCCATCTTCC |
| ONORMANBULBIEJR4_F | atgcggaggaatcacttccatATGGATCACGCTGAGTATGC |
| ONORMANBULBIEJR4_R | tgcaggatccgactcgagtgtcgacTCAGCGCTGGATGCTTC |
| ONORMANBULBIEJR5_F | atgcggaggaatcacttccatATGTCTGATGATGTGACAGC |
| ONORMANBULBIEJR5_R | tgcaggatccgactcgagtgtcgacTCAGTGGAGTTCTCCACG |
| ONORMANBULBIEJR6_F | atgcggaggaatcacttccatATGGCTTTCAACAACTTCATTC |
| ONORMANBULBIEJR6_R | tgcaggatccgactcgagtgtcgacTCAGCTGCCCGTCTTATTG |
| ONORMANBULBIEJR7_F | atgcggaggaatcacttccatATGTTGCTTGCTACCGC |
| ONORMANBULBIEJR7_R | tgcaggatccgactcgagtgtcgacTCATAGCCGGTGAATCG |
| ONORMANBULBIEJR8_F | atgcggaggaatcacttccatATGACGTTTCCAACCGC |
| ONORMANBULBIEJR8_R | tgcaggatccgactcgagtgtcgacTCACACCTTCCGAAGTTC |
| ONORMANBULBIEJR9_F | atgcggaggaatcacttccataTGGCTAACGGTCCAACG |
| ONORMANBULBIEJR9_R | tgcaggatccgactcgagtgtcgactcaGTCACCATACGCGG |
| ONORMANBULBIEJR10_F | atgcggaggaatcacttccatATGGTGACTGATTCAGCG |
| ONORMANBULBIEJR10_R | tgcaggatccgactcgagtgtcgacTCAGATGTACTGAACACCG |
| ONORMANBULBIEJR11_F | atgcggaggaatcacttccataTGACGCAGCCATTGACC |
| ONORMANBULBIEJR11_R | tgcaggatccgactcgagtgtcgactcaGCTGCCGTCCGAGTAC |
| ONORMANBULBIEJR13_F | atgcggaggaatcacttccatATGTCTGTGAAGAAACCTGAAAAC |
| ONORMANBULBIEJR13_R | tgcaggatccgactcgagtgtcgacTCACCAGCCGAACAGATC |
| ONORMANBULBIEJR12_F | atgcggaggaatcacttccatATGTCTGTGAAGAAACCTGAAAAC |
| ONORMANBULBIEJR12_R | tgcaggatccgactcgagtgtcgacTCACTCGCTATCTGTCTC |
| ONORMANBULBIEJR14_F | atgcggaggaatcacttccataTGCCGATCTATGTGGAC |
| ONORMANBULBIEJR14_R | tgcaggatccgactcgagtgtcgactcaCCCTCCCGGCATGAC |
| ONORMANBULBIEJR15_F | atgcggaggaatcacttccataTGGCTAAGAAGCATTACCCCG |
| ONORMANBULBIEJR15_R | tgcaggatccgactcgagtgtcgactcaCATCGGGTAGCGGCG |
| ONORMANBULBIEJR16_F | atgcggaggaatcacttccataTGTCGAAGTTTGAACGCG |
| ONORMANBULBIEJR16_R | tgcaggatccgactcgagtgtcgactcaTCCCTGCGGTGACAG |
| ONORMANBULBIEJR17_F | atgcggaggaatcacttccatATGGAGCGTGCCCTAATG |
| ONORMANBULBIEJR17_R | tgcaggatccgactcgagtgtcgacTCATGACAGCGGAAGAACC |
| ONORMANBULBIEJR18_F | atgcggaggaatcacttccatATGACTGATTCGTTTGATCCG |
| ONORMANBULBIEJR18_R | tgcaggatccgactcgagtgtcgacTCATTCGTCCGTCGC |
| ONORMANBULBIEJR19_F | atgcggaggaatcacttccatATGACAATTCAAGGCTACTTCG |
| ONORMANBULBIEJR19_R | tgcaggatccgactcgagtgtcgacTCAGCCCGCCCCGATTTG |
| ONORMANBULBIEJR20_F | atgcggaggaatcacttccatATGGCGGGCTGGTGG |
| ONORMANBULBIEJR20_R | tgcaggatccgactcgagtgtcgacTCATATTGGTGACAGAGCGAC |

| | |
|---|---|
| ONORMANBULBIEJR21_F | atgcggaggaatcacttccatATGGGCATTCCCAATGC |
| ONORMANBULBIEJR21_R | tgcaggatccgactcgagtgtcgacTCAAGCGACCACGCG |
| ONORMANBULBIEJR22_F | atgcggaggaatcacttccataTGCGGATCAAGACCGATCACC |
| ONORMANBULBIEJR22_R | tgcaggatccgactcgagtgtcgactcaGGGCTGGTCCCTCAAC |
| ONORMANBULBIEJR23_F | atgcggaggaatcacttccatATGGCTTACGACAAGCAG |
| ONORMANBULBIEJR23_R | tgcaggatccgactcgagtgtcgacTCACGGCACCACCAC |
| ONORMANBULBIEJR24_F | atgcggaggaatcacttccatATGAAAGTTTGGAACGGC |
| ONORMANBULBIEJR24_R | tgcaggatccgactcgagtgtcgacTCATTCCCACTCGACCAG |
| ONORMANBULBIEJR25_F | atgcggaggaatcacttccataTGACAATCATCGACCGTATGATCG |
| ONORMANBULBIEJR25_R | tgcaggatccgactcgagtgtcgactcaACGGTTCGGGTCACC |
| ONORMANBULBIEJR26_F | atgcggaggaatcacttccatATGCTACGCAACACCATC |
| ONORMANBULBIEJR26_R | tgcaggatccgactcgagtgtcgacTCACCAGGTGTCGGTG |
| ONORMANBULBIEJR27_F | atgcggaggaatcacttccatATGAAGATCCACGTTCAATCC |
| ONORMANBULBIEJR27_R | tgcaggatccgactcgagtgtcgacTCATGACAGTCTCCTGACC |
| ONORMANBULBIEJR28_F | atgcggaggaatcacttccatATGACCAAACGAGTAGCG |
| ONORMANBULBIEJR28_R | tgcaggatccgactcgagtgtcgacTCAGATGTAGAAGAGGGTG |
| ONORMANBULBIEJR29_F | atgcggaggaatcacttccatATGGACCGTCTCGGAATC |
| ONORMANBULBIEJR29_R | tgcaggatccgactcgagtgtcgacTCACCGTTTGCTCCCG |
| ONORMANBULBIEJR30_F | atgcggaggaatcacttccatATGACCACGAAAGATCAAGTC |
| ONORMANBULBIEJR30_R | tgcaggatccgactcgagtgtcgacTCAGGACTTGCCCTTG |
| ONORMANBULBIEJR31_F | atgcggaggaatcacttccatATGCGCATCGACGGG |
| ONORMANBULBIEJR31_R | tgcaggatccgactcgagtgtcgacTCATGTGCGTAGGTAGTCG |
| ONORMANBULBIEJR32_F | atgcggaggaatcacttccatATGCTGACACGTTCATTCTG |
| ONORMANBULBIEJR32_R | tgcaggatccgactcgagtgtcgacTCATGCGGCGGTGAC |
| ONORMANBULBIEJR33_F | atgcggaggaatcacttccatATGATCTGGGAATCGGTGC |
| ONORMANBULBIEJR33_R | tgcaggatccgactcgagtgtcgacTCACCGGTCGCGTCG |
| ONORMANBULBIEJR34_F | atgcggaggaatcacttccatATGTCACTCTTGGCCG |
| ONORMANBULBIEJR34_R | tgcaggatccgactcgagtgtcgacTCAGTCAGCGACATGATG |
| ONORMANBULBIEJR35_F | atgcggaggaatcacttccatATGTCGCTGACTAGCGAC |
| ONORMANBULBIEJR35_R | tgcaggatccgactcgagtgtcgacTCACACCCCTCTCACAG |
| ONORMANBULBIEJR36_F | atgcggaggaatcacttccatATGAACGTTCGAGTGTGC |
| ONORMANBULBIEJR36_R | tgcaggatccgactcgagtgtcgacTCAGCAGACATCACGAC |
| ONORMANBULBIEJR37_F | atgcggaggaatcacttccataTGATGTCTGCTGACCCTG |
| ONORMANBULBIEJR37_R | tgcaggatccgactcgagtgtcgactcaCTCACCAGACGACTC |
| ONORMANBULBIEJR38_F | atgcggaggaatcacttccatATGAGTGATCCGCAGTTG |
| ONORMANBULBIEJR38_R | tgcaggatccgactcgagtgtcgacTCAAGCCAGGACGTAAAC |
| ONORMANBULBIEJR39_F | atgcggaggaatcacttccataTGGCTGACGCTTTGAAAC |
| ONORMANBULBIEJR39_R | tgcaggatccgactcgagtgtcgactcaGTTTGGGTACACCCG |
| ONORMANBULBIEJR40_F | atgcggaggaatcacttccatATGAGTGTCCTCGCTTG |
| ONORMANBULBIEJR40_R | tgcaggatccgactcgagtgtcgacTCAGCCATTGGCTTCTTC |
| ONORMANBULBIEJR41_F | atgcggaggaatcacttccatATGAACACCGATGATCGTTG |
| ONORMANBULBIEJR41_R | tgcaggatccgactcgagtgtcgacTCATCGGCTTGCTTCTTC |
| ONORMANBULBIEJR42_F | atgcggaggaatcacttccatATGTGCACCGCTTGC |

| | |
|---|---|
| ONORMANBULBIEJR42_R | tgcaggatccgactcgagtgtcgacTCAGTTGCACATCTCGCATC |
| ONORMANBULBIEJR43_F | atgcggaggaatcacttccataTGACTCACTCACCGGGAAAC |
| ONORMANBULBIEJR43_R | tgcaggatccgactcgagtgtcgactcaCTCCCCGCCCTGAAG |
| ONORMANBULBIEJR44_F | atgcggaggaatcacttccatATGGCAACTAAGAAACGCAG |
| ONORMANBULBIEJR44_R | tgcaggatccgactcgagtgtcgacTCATTTTGTTTCTAACGCAAACG |
| ONORMANBULBIEJR45_F | atgcggaggaatcacttccatATGACCACCAATGATCTGTC |
| ONORMANBULBIEJR45_R | tgcaggatccgactcgagtgtcgacTCACTCCGTCATGGTC |
| ONORMANBULBIEJR46_F | atgcggaggaatcacttccataTGCCCGGCGCAACGTCC |
| ONORMANBULBIEJR46_R | tgcaggatccgactcgagtgtcgactcaCGCATCCGGTTGCGAG |
| ONORMANBULBIEJR47_F | atgcggaggaatcacttccataTGCACCTGTTTGCACGC |
| ONORMANBULBIEJR47_R | tgcaggatccgactcgagtgtcgac tcaGTCCTTGCGGCGTTG |
| ONORMANBULBIEJR48_F | atgcggaggaatcacttccatATGGTCAAACAGTCCTACG |
| ONORMANBULBIEJR48_R | tgcaggatccgactcgagtgtcgacTCATGCAGCGGGGTTTTTC |
| ONORMANBULBIEJR49_F | atgcggaggaatcacttccatATGTCTGAACTACAGCATACC |
| ONORMANBULBIEJR49_R | tgcaggatccgactcgagtgtcgacTCATCGCGTTTCCCTC |
| ONORMANBULBIEJR50_F | atgcggaggaatcacttccatATGAGCACTCCCAGATGG |
| ONORMANBULBIEJR50_R | tgcaggatccgactcgagtgtcgacTCACGCGGACCTCGG |
| ONORMANBULBIEJR51_F | atgcggaggaatcacttccatATGAACCTCACTGAGTATGAGC |
| ONORMANBULBIEJR51_R | tgcaggatccgactcgagtgtcgacTCATCGCCGTCCCGAC |
| ONORMANBULBIEJR52_F | atgcggaggaatcacttccatATGAGCCAGGACGTC |
| ONORMANBULBIEJR52_R | tgcaggatccgactcgagtgtcgacTCACAACCGGCCCTC |
| ONORMANBULBIEJR53_F | atgcggaggaatcacttccatATGAGTACGTCTGCTCCTAAG |
| ONORMANBULBIEJR53_R | tgcaggatccgactcgagtgtcgacTCACGCTGTCTCCCC |
| ONORMANBULBIEJR54_F | atgcggaggaatcacttccatATGAATCTTGTTGAGCGTTTG |
| ONORMANBULBIEJR54_R | tgcaggatccgactcgagtgtcgacTCATCTCGGGGTGATG |
| ONORMANBULBIEJR55_F | atgcggaggaatcacttccataTGCTAGATCGAGATTCTAAACCC |
| ONORMANBULBIEJR55_R | tgcaggatccgactcgagtgtcgactcaTGCTGCTTCTCCTGTC |
| ONORMANBULBIEJR56_F | atgcggaggaatcacttccatATGAGGCGCAGTGAGAAG |
| ONORMANBULBIEJR56_R | tgcaggatccgactcgagtgtcgacTCACACCCACCCAGTTC |
| ONORMANBULBIEJR57_F | atgcggaggaatcacttccataTGATGGCGAACTCCCCG |
| ONORMANBULBIEJR57_R | tgcaggatccgactcgagtgtcgactcaCGCGCCCCACCTC |
| ONORMANBULBIEJR58_F | atgcggaggaatcacttccatATGAGCATCGACTGGTTC |
| ONORMANBULBIEJR58_R | tgcaggatccgactcgagtgtcgacTCATGCAACGGACTCC |
| ONORMANBULBIEJR59_F | atgcggaggaatcacttccatATGACCCTGCTCGATC |
| ONORMANBULBIEJR59_R | tgcaggatccgactcgagtgtcgacTCATGCCACCTGATCC |
| ONORMANBULBIEJR60_F | atgcggaggaatcacttccataTGAGCAACGGGAACAG |
| ONORMANBULBIEJR60_R | tgcaggatccgactcgagtgtcgactcaCGAAGCCTCCAAC |
| ONORMANBULBIEJR61_F | atgcggaggaatcacttccataTGATGTGCGTGTGCGGC |
| ONORMANBULBIEJR61_R | tgcaggatccgactcgagtgtcgactcaGTTTTCGTCCTTGTCTCGATAT |
| ONORMANBULBIEJR62_F | atgcggaggaatcacttccataTGGTTGTTGATACACGGG |
| ONORMANBULBIEJR62_R | tgcaggatccgactcgagtgtcgactcaTGCTGCTGTCCAC |
| ONORMANBULBIEJR63_F | atgcggaggaatcacttccataTGAGCGATCCGACTG |
| ONORMANBULBIEJR63_R | tgcaggatccgactcgagtgtcgactcaTTGTTCGATCTCCTTCAAC |

| | |
|---|---|
| ONORMANBULBIEJR64_F | atgcggaggaatcacttccataTGACCCTCAAAACCCG |
| ONORMANBULBIEJR64_R | tgcaggatccgactcgagtgtcgactcaTGAGGCGTCCGC |
| ONORMANBULBIEJR65_F | atgcggaggaatcacttccataTGACCCGCCGGTTTAC |
| ONORMANBULBIEJR65_R | tgcaggatccgactcgagtgtcgactcaCTTAGCAGCCTCCACAG |
| ONORMANBULBIEJR66_F | atgcggaggaatcacttccataTGACCGGCCACGTGACG |
| ONORMANBULBIEJR66_R | tgcaggatccgactcgagtgtcgactcaTTTACCTCTCACTCTTCTCATC |
| ONORMANBULBIEJR67_F | atgcggaggaatcacttccataTGACCGACTACATCACC |
| ONORMANBULBIEJR67_R | tgcaggatccgactcgagtgtcgactcaTGACGCGTCATCC |
| ONORMANBULBIEJR68_F | atgcggaggaatcacttccataTGACCGCCGAGTCG |
| ONORMANBULBIEJR68_R | tgcaggatccgactcgagtgtcgactcaTGGAACCTGCCAGTTG |
| ONORMANBULBIEJR69_F | atgcggaggaatcacttccataTGATCACCGTTGCTTGCG |
| ONORMANBULBIEJR69_R | tgcaggatccgactcgagtgtcgactcaCGAGGCCGCCTC |
| ONORMANBULBIEJR70_F | atgcggaggaatcacttccatATGGCCCACGTTTTGTATC |
| ONORMANBULBIEJR70_R | tgcaggatccgactcgagtgtcgacTCAGGAATCACCTCCAGG |
| ONORMANBULBIEJR71_F | atgcggaggaatcacttccataTGGGAAGGAAAGCCAC |
| ONORMANBULBIEJR71_R | tgcaggatccgactcgagtgtcgactcaCAACGCCACACC |
| ONORMANBULBIEJR72_F | atgcggaggaatcacttccataTGAAGGACTGGCGTGGC |
| ONORMANBULBIEJR72_R | tgcaggatccgactcgagtgtcgactcaGTGGCCTTGGGAACC |
| ONORMANBULBIEJR73_F | atgcggaggaatcacttccataTGAGCCTCGAGATTCGC |
| ONORMANBULBIEJR73_R | tgcaggatccgactcgagtgtcgactcaGTCCGGCCACCTG |
| ONORMANBULBIEJR74_F | atgcggaggaatcacttccataTGACGTTGTTTGTGTCG |
| ONORMANBULBIEJR74_R | tgcaggatccgactcgagtgtcgactcaTTCGTCGCCTTTCG |
| ONORMANBULBIEJR75_F | atgcggaggaatcacttccataTGAACAACCCCGAGTTG |
| ONORMANBULBIEJR75_R | tgcaggatccgactcgagtgtcgactcaTTCCTCCGCCTCTTC |
| ONORMANBULBIEJR76_F | atgcggaggaatcacttccataTGTTTCCGATTACCGACAC |
| ONORMANBULBIEJR76_R | tgcaggatccgactcgagtgtcgactcaTCTGGGGTCATCGTC |
| ONORMANBULBIEJR77_F | atgcggaggaatcacttccataTGGAGGAATCTGGAGACC |
| ONORMANBULBIEJR77_R | tgcaggatccgactcgagtgtcgactcaTGCCTCGCTCCATC |
| ONORMANBULBIEJR78_F | atgcggaggaatcacttccataTGGAGCGAGGCATG |
| ONORMANBULBIEJR78_R | tgcaggatccgactcgagtgtcgactcaCCGCGTAACCTC |
| ONORMANBULBIEJR79_F | atgcggaggaatcacttccataTGATTCAGGTTCATTGCAAG |
| ONORMANBULBIEJR79_R | tgcaggatccgactcgagtgtcgactcaTGGTTGTCCTTTCGTG |
| ONORMANBULBIEJR80_F | atgcggaggaatcacttccataTGACCAACGAGTTACGTG |
| ONORMANBULBIEJR80_R | tgcaggatccgactcgagtgtcgactcaCTTGTCTTCCCCCTC |
| ONORMANBULBIEJR81_F | atgcggaggaatcacttccataTGAGTTGGGTCGAAGGC |
| ONORMANBULBIEJR81_R | tgcaggatccgactcgagtgtcgactcaTTGTTCCCCCTCGG |
| ONORMANBULBIEJR82_F | atgcggaggaatcacttccataTGAGCGACGCGCAG |
| ONORMANBULBIEJR82_R | tgcaggatccgactcgagtgtcgactcaCTCATCGGGTATCTCCATC |
| ONORMANBULBIEJR83_F | atgcggaggaatcacttccataTGAGTGATGTGGTAGAGC |
| ONORMANBULBIEJR83_R | tgcaggatccgactcgagtgtcgactcaTCGTTCCCCCTC |
| ONORMANBULBIEJR84_F | atgcggaggaatcacttccataTGAACGGCAAACGC |
| ONORMANBULBIEJR84_R | tgcaggatccgactcgagtgtcgactcaTTTTTCCTCGCCCTC |
| ONORMANBULBIEJR85_F | atgcggaggaatcacttccataTGAGCACTCCTGAGC |

| | |
|---|---|
| ONORMANBULBIEJR85_R | tgcaggatccgactcgagtgtcgactcaAAAGGGAGGAGCC |
| ONORMANBULBIEJR86_F | atgcggaggaatcacttccataTGAGCGGCCGCAAGATC |
| ONORMANBULBIEJR86_R | tgcaggatccgactcgagtgtcgactcaCTCCGGTTTGTGGGATTC |
| ONORMANBULBIEJR87_F | atgcggaggaatcacttccataTGACCGTGATCTACATCTTC |
| ONORMANBULBIEJR87_R | tgcaggatccgactcgagtgtcgactcaTTTGTGGTGTCCTTTG |
| ONORMANBULBIEJR88_F | atgcggaggaatcacttccataTGAGCGTCTACGCACTG |
| ONORMANBULBIEJR88_R | tgcaggatccgactcgagtgtcgactcaCTTCTCAGGCCTCCTAG |
| ONORMANBULBIEJR89_F | atgcggaggaatcacttccataTGGAGGTGCCCATGAGC |
| ONORMANBULBIEJR89_R | tgcaggatccgactcgagtgtcgactcaTAGGGTTTCCTGAAGTTCGTCG |
| ONORMANBULBIEJR90_F | atgcggaggaatcacttccataTGACCCAACCAGCAGAG |
| ONORMANBULBIEJR90_R | tgcaggatccgactcgagtgtcgactcaTTCGAGGACTCCTGC |
| ONORMANBULBIEJR91_F | atgcggaggaatcacttccataTGAGCACCTTCCCCG |
| ONORMANBULBIEJR91_R | tgcaggatccgactcgagtgtcgactcaTGGCCGGTTTCTCC |
| ONORMANBULBIEJR92_F | atgcggaggaatcacttccataTGACCCGACGTTTTCG |
| ONORMANBULBIEJR92_R | tgcaggatccgactcgagtgtcgactcaCCCGAACCACCAG |
| ONORMANBULBIEJR93_F | atgcggaggaatcacttccataTGAGGCGCGCAATCGCCG |
| ONORMANBULBIEJR93_R | tgcaggatccgactcgagtgtcgactcaCTCCTTCACCCGTCCG |
| ONORMANBULBIEJR94_F | atgcggaggaatcacttccataTGGATCCCCAGAAGTTCCGC |
| ONORMANBULBIEJR94_R | tgcaggatccgactcgagtgtcgactcaGTCCGCCGCCTCGTAC |
| ONORMANBULBIEJR95_F | atgcggaggaatcacttccataTGCCGCTCAAACACC |
| ONORMANBULBIEJR95_R | tgcaggatccgactcgagtgtcgactcaTTTGGGATATTCGCCTG |
| ONORMANBULBIEJR96_F | atgcggaggaatcacttccataTGAGCAGCCTCACAGAC |
| ONORMANBULBIEJR96_R | tgcaggatccgactcgagtgtcgactcaCTGTTCGACCGCTTTC |
| ONORMANBULBIEJR97_F | atgcggaggaatcacttccataTGATCCACATCGAAGTTGAC |
| ONORMANBULBIEJR97_R | tgcaggatccgactcgagtgtcgactcaTCCATTGGTGCAGTC |
| ONORMANBULBIEJR98_F | atgcggaggaatcacttccataTGGATGAGGCAGCCC |
| ONORMANBULBIEJR98_R | tgcaggatccgactcgagtgtcgactcaCTCGTACCATTCGCC |
| ONORMANBULBIEJR99_F | atgcggaggaatcacttccataTGACCGACGTCGTGATC |
| ONORMANBULBIEJR99_R | tgcaggatccgactcgagtgtcgactcaTTTGTGCCTCCACCAG |
| ONORMANBULBIEJR100_F | atgcggaggaatcacttccatATGGACCAGAACCTGAAAC |
| ONORMANBULBIEJR100_R | tgcaggatccgactcgagtgtcgacTCATCGTGGCCTTATGC |
| ONORMANBULBIEJR101_F | atgcggaggaatcacttccatATGACCCACATCATCGG |
| ONORMANBULBIEJR101_R | tgcaggatccgactcgagtgtcgacTCAGATCGGTTGGGTG |
| ONORMANBULBIEJR102_F | atgcggaggaatcacttccatATGCCACGCGCGCCTAAG |
| ONORMANBULBIEJR102_R | tgcaggatccgactcgagtgtcgacTCAGGTGTTCGATCCGCGG |
| ONBJ_14IA | TCAACCAGGTGTTCAACAAGG |
| ONBJ_14IB | ATTGGAATCATTCGAACACG |
| ONBJ_14IC | TTCTGCGGGCTGTACAACC |
| ONBJ_15IA | AGACTTCGACACAGACACC |
| ONBJ_16IA | ATGGATTTGTGGCTTCCAGG |
| ONBJ_18IA | GACTGGACCTTGATGTCTGG |
| ONBJ_18IB | CGTTGATCCACTCATCGG |
| ONBJ_19IA | TCCAACTTCTCGTTGATCGG |

| | |
|---|---|
| PEXTRA_SEQF | GTACCCGTGTGTACGACCAGC |
| PEXTRA_UNIVERSALF | ATGAGCACGATCCGCATGCG |
| PEXTRA_UNIVERSALR | CCCTTCGAGACCATAGATCTGTTCC |
| PEXTRA_F | GTCGACACTCGAGTCGGATCCTG |
| PEXTRA_R | ATGGAAGTGATTCCTCCGCATGC |
| ONBJ54_RT-PCR_F | GACGCCATCGACTCCACCAAGC |
| ONBJ54_RT-PCR_R | CTATCTCGGGGTGATGCGGAAC |
| ODH458 | atatatcatatgGGCGCCGCCGGTGTTGCCCTG |
| ODH459 | atatatACTAGTttaACCCTCAGTAGTGTCCAACGG |
| ODH462 | CATGAACGCCGCTGTGGGTaGTGTTGGGGCGAACTTCATTTC |
| ODH463 | GAAATGAAGTTCGCCCCAACACtACCCACAGCGGCGTTCATG |

**Table S2: Oligonucleotide sequences used for BRED**

| Oligo Name | Gene Targets | Usage | Oligo Sequence (5' to 3') |
|---|---|---|---|
| oBW54 | *33* | substrate amplification | CATGCTGACACGTTCATTCTGG |
| oBW55 | *33* | substrate amplification | TGGTGCCGCCACGCTTGCGCGTTC |
| oBW56 | *34* | substrate amplification | ACAGATCGACGCGAAAACCG |
| oBW57 | *34* | substrate amplification | GCGTCCACCTTTGTCCCATTCG |
| oBW58 | *35* | substrate amplification | AGGGAACACCGAATGTCACTCTTGG |
| oBW59 | *35* | substrate amplification | GTGATCACAGTCGTAGCACCCAG |
| oBW60 | *47* | substrate amplification | TGGTGGCTAACTTTCGCGCACTGG |
| oBW61 | *47* | substrate amplification | GTAAGGGGCAGAGACTCCGCTTTCG |
| oBW62 | *54* | substrate amplification | ATGGGTCCTGCTGGTGGTTTGG |
| oBW63 | *54* | substrate amplification | CCAGTTCACTCATGACCGCC |
| oBW64 | *55* | substrate amplification | GTGTCACGCGCAGACCTCACC |
| oBW65 | *55* | substrate amplification | GATGCTTCGTACATGGTGGC |
| oBW66 | *59* | substrate amplification | CAGACAGGCTGTTTCAGGAATGCC |
| oBW67 | *59* | substrate amplification | CCGTCATCAAGCAAATATTGGGCG |
| oBW68 | *68* | substrate amplification | TCCAGCTTGGTTTCGTCGAG |
| oBW69 | *68* | substrate amplification | CTCATGTCATCAGGGTCGATACC |
| oBW102 | *6* | substrate amplification | GTGGAATGAGTTCGTCAACAAGC |
| oBW103 | *6* | substrate amplification | GCTTCGACTTCATCCACCACG |
| oBW104 | *37* | substrate amplification | GGCGAGGACAAGCTGATGG |
| oBW105 | *37* | substrate amplification | CGGAACGACTCGGTATATCG |
| oBW106 | *56* | substrate amplification | CCAAACAACCTGGGCTGACC |
| oBW107 | *56* | substrate amplification | GGCACGTTCCTTCTCTGAG |
| oBW108 | *58* | substrate amplification | AGCGTGGCGAGCTGGCACCG |
| oBW109 | *58* | substrate amplification | TCCCGCTCCGACAACCCCC |
| oBW111 | *67* | substrate amplification | ACCAGCACTGCCACCGCATG |
| oBW112 | *67* | substrate amplification | ATCACCTGAACAGCCCCTTCACG |

| | | | |
|---|---|---|---|
| oBW113 | *71* | substrate amplification | CCGGTCACGTTTTCGGCGG |
| oBW114 | *71* | substrate amplification | GATTGCGGGGGCTGTACGCC |
| oBW115 | *72* | substrate amplification | CCCATATCAATCACCCGAACC |
| oBW116 | *72* | substrate amplification | ATCAGATGACCGACTTCGCC |
| oBW117 | *99* | substrate amplification | ACACGAGGACACCACAATCC |
| oBW118 | *99* | substrate amplification | TTGGTGTGGGCCACGATGC |
| oBW119 | *101* | substrate amplification | CCGTGGAGCTGGTGGAGGC |
| oBW120 | *101* | substrate amplification | CGTGATCGGGGTTGGATGTG |
| oBW70 | *33* | flanking PCR | CCGGCATGTTCTTCGCGAAACG |
| oBW71 | *33* | flanking PCR | GTCAGCGACATGATGGTTCC |
| oBW72 | *34* | flanking PCR | TGTTCGAGGGCAACGCAAGG |
| oBW73 | *34* | flanking PCR | CCGCTTGGGATAGCAGGTTGTGGAC |
| oBW74 | *35* | flanking PCR | CGGCTTCAAACAGATTCAACGGGAC |
| oBW75 | *35* | flanking PCR | CCACACCGACCGCCACAACTCC |
| oBW76 | *47* | flanking PCR | TTCTGTCGTCGTCGTTGTTGTTG |
| oBW77 | *47* | flanking PCR | GTGGGGGCTTCTTCTATGCAGCGG |
| oBW78 | *54* | flanking PCR | GAGTGCCCGCACCTGAATGGGAC |
| oBW79 | *54* | flanking PCR | TGAGGTAGTCGTGCAGAAGCCCAAC |
| oBW80 | *55* | flanking PCR | GCCGCGATCCGACAGCACAC |
| oBW81 | *55* | flanking PCR | CAGGTGTTCAGGGCAGAACGATGC |
| oBW82 | *59* | flanking PCR | TGGCAACATGGAAGCCCACC |
| oBW83 | *59* | flanking PCR | CAACCAGCGGAACTTCTTGACC |
| oBW84 | *68* | flanking PCR | GATGAGCGTCACATTTGCCC |
| oBW85 | *68* | flanking PCR | GCTGCCCTGTCGCGCTGTAGAAG |
| oBW121 | *6* | flanking PCR | GATACCGAGGAAGAGATGCG |
| oBW122 | *6* | flanking PCR | ACCGTCTTGGTTGAAGTCG |
| oBW123 | *37* | flanking PCR | GCACATCAAAAACCATCCCG |
| oBW124 | *37* | flanking PCR | AGAAACGATGGGAAGAAGCC |
| oBW125 | *56* | flanking PCR | TTCTAAACCCTCATGGTGGG |

| | | | |
|---|---|---|---|
| oBW126 | *56* | flanking PCR | TGAACCGTGCCGGTTTCAC |
| oBW127 | *58* | flanking PCR | GGCTTCTCAGAGAAGGAACG |
| oBW128 | *58* | flanking PCR | CGATCGCTTTGAGCAGAGGC |
| oBW130 | *67* | flanking PCR | TCCTGGTAACGAAGCCGAG |
| oBW131 | *67* | flanking PCR | GGTGGAGTTGAAGCATTTCG |
| oBW132 | *71* | flanking PCR | GATCTCTTCTCCCCAGAACC |
| oBW133 | *71* | flanking PCR | GAGGCTCACTGTTCGTCTCC |
| oBW134 | *72* | flanking PCR | GAGCCCCCATTGCGTTG |
| oBW135 | *72* | flanking PCR | GTTGTTCAACAGCAGGTCC |
| oBW136 | *99* | flanking PCR | AACGAGCCTTGGATGCTTCC |
| oBW137 | *99* | flanking PCR | GCCCACGGTTTCCATGAGCC |
| oBW138 | *101* | flanking PCR | ACCAACAACGACGCATCTGG |
| oBW139 | *101* | flanking PCR | TTGTCGAGTGTGTCTGTGC |
| oBW86 | *33* | sgRNA construction | GGGAGCCGTTCCGATCGCTGCGATC |
| oBW87 | *33* | sgRNA construction | AAACGATCGCAGCGATCGGAACGGC |
| oBW88 | *34* | sgRNA construction | GGGAGCCGCGTTTCGCGGACTCCAA |
| oBW89 | *34* | sgRNA construction | AAACTTGGAGTCCGCGAAACGCGGC |
| oBW90 | *35* | sgRNA construction | GGGAGTGGTTTTTGATGTGCGGACG |
| oBW91 | *35* | sgRNA construction | AAACCGTCCGCACATCAAAAACCAC |
| oBW92 | *47* | sgRNA construction | GGGAGTAGTCGTCGCCGGACTTCGG |
| oBW93 | *47* | sgRNA construction | AAACCCGAAGTCCGGCGACGACTAC |
| oBW94 | *54* | sgRNA construction | GGGAGCCTCAGCCTCCGCAGCCACA |
| oBW95 | *54* | sgRNA construction | AAACTGTGGCTGCGGAGGCTGAGGC |
| oBW96 | *55* | sgRNA construction | GGGAGTCCTGTCGTGAGGTAGTCGT |
| oBW97 | *55* | sgRNA construction | AAACACGACTACCTCACGACAGGAC |
| oBW98 | *59* | sgRNA construction | GGGAGTTCGCGCTTTTACCCTTCTC |
| oBW99 | *59* | sgRNA construction | AAACGAGAAGGGTAAAAGCGCGAAC |
| oBW100 | *68* | sgRNA construction | GGGAGGTTGATGGTGTCGCCTTGCT |
| oBW101 | *68* | sgRNA construction | AAACAGCAAGGCGACACCATCAACC |

| | | | |
|---|---|---|---|
| oBW140 | *6* | sgRNA construction | GGGAGCAGCGCGGTCCCGTTGTCCA |
| oBW141 | *6* | sgRNA construction | AAACTGGACAACGGGACCGCGCTGC |
| oBW142 | *37* | sgRNA construction | GGGAGGGATCGCGCCGCGAACAGGG |
| oBW143 | *37* | sgRNA construction | AAACCCCTGTTCGCGGCGCGATCCC |
| oBW144 | *56* | sgRNA construction | GGGAGGTTCCGCTCAGGTGTTCAGG |
| oBW145 | *56* | sgRNA construction | AAACCCTGAACACCTGAGCGGAACC |
| oBW146 | *58* | sgRNA construction | GGGAGGAGTTTCGGGCGCCGCCGCA |
| oBW147 | *58* | sgRNA construction | AAACTGCGGCGGCGCCCGAAACTCC |
| oBW150 | *67* | sgRNA construction | GGGAGAACGCGGCGATAACTTGCCC |
| oBW151 | *67* | sgRNA construction | AAACGGGCAAGTTATCGCCGCGTTC |
| oBW152 | *71* | sgRNA construction | GGGAGGTCGTGCTTGATCCACGACC |
| oBW153 | *71* | sgRNA construction | AAACGGTCGTGGATCAAGCACGACC |
| oBW154 | *72* | sgRNA construction | GGGAGAGCGATCCTGTGCGAATGAG |
| oBW155 | *72* | sgRNA construction | AAACCTCATTCGCACAGGATCGCTC |
| oBW156 | *99* | sgRNA construction | GGGAGGGGTAGCAGTCGTCGTCGAA |
| oBW157 | *99* | sgRNA construction | AAACTTCGACGACGACTGCTACCCC |
| oBW158 | *101* | sgRNA construction | GGGAGAGCGCTGCGTCGAGTTGGTC |
| oBW159 | *101* | sgRNA construction | AAACGACCAACTCGACGCAGCGCTC |
| oBW47 | | psgRNA sequencing | TTTGAATTCTCTGACCAGGG |

**Table S3: sgRNA sequences for CRISPR selection**

| sgRNA | Gene Targets | PAM site | sgRNA Targeting Sequence (5’ to 3’) |
|---|---|---|---|
| sgRNA-1 | *33* | NNGGAAG | CCGTTCCGATCGCTGCGATC |
| sgRNA-2 | *34* | NNAGGAT | CCGCGTTTCGCGGACTCCAA |
| sgRNA-3 | *35* | NNGGAAG | TGGTTTTTGATGTGCGGACG |
| sgRNA-4 | *47* | NNAGAAG | TAGTCGTCGCCGGACTTCGG |
| sgRNA-5 | *54* | NNAGAAT | CCTCAGCCTCCGCAGCCACA |
| sgRNA-6 | *55* | NNAGAAG | TCCTGTCGTGAGGTAGTCGT |
| sgRNA-7 | *59* | NNGGAAA | TTCGCGCTTTTACCCTTCTC |
| sgRNA-8 | *68* | NNAGAAT | GTTGATGGTGTCGCCTTGCT |
| sgRNA-9 | *6* | NNAGAAG | CAGCGCGGTCCCGTTGTCCA |
| sgRNA-10 | *37* | NNAGCAG | GGATCGCGCCGCGAACAGGG |
| sgRNA-11 | *56* | NNAGAAC | GTTCCGCTCAGGTGTTCAGG |
| sgRNA-12 | *58* | NNAGCAT | GAGTTTCGGGCGCCGCCGCA |
| sgRNA-14 | *67* | NNGGAAG | AACGCGGCGATAACTTGCCC |
| sgRNA-15 | *71* | NNAGAAG | GTCGTGCTTGATCCACGACC |
| sgRNA-16 | *72* | NNGGAAA | AGCGATCCTGTGCGAATGAG |
| sgRNA-17 | *99* | NNGGAAG | GGGTAGCAGTCGTCGTCGAA |
| sgRNA-18 | *101* | NNGGAAG | AGCGCTGCGTCGAGTTGGTC |

**Table S4: Summary Table of BRED and CRISPY-BRED deletion studies**

| Gene deletion | BRED | | CRISPY-BRED | Average plaque diameter (mm) ± standard deviation | Efficiency of Lysogeny (%) ± standard deviation (number of independent tests) |
|---|---|---|---|---|---|
| | mixed 1° plaques[1] | mutant 2° plaques[2] | mutant plaques[3] | WT= 2.4 ± 0.5 | WT=22%±10 (*n*=5) |
| *33* | 8/16 | 0/32 | 23/24 | 0.68 ± 0.2 | |
| *34* | 8/16 | 6/24 | 15/16 | 2.2 ± 0.5 | |
| *35* | 7/16 | 3/31 | 16/16 | 2.4 ± 0.4 | |
| *54* | 5/16 | 0/32 | 15/16 | 2.0 ± 0.4 | |
| *59* | 6/16 | 0/40 | 16/16 | 2.5 ± 0.4 | 20.5%±15 (*n=2*) |
| *68* | 1/13 | 1/24 | 16/16 | 2.5 ± 0.6 | 19.5%±4 (*n=2*) |
| *6* | | | *No plaques recovered on aTc* | | |
| *37* | | | 8/8 | 2.4 ± 0.4 | |
| *45* | | | 8/8 | 2.4 ± 0.6 | 12%±1 (*n=3*) |
| *47* | | | 22/24 | 2.4 ± 0.5 | <0.0001% (*n*=5) |
| *56* | | | 16/16 | 2.4 ± 0.5 | |
| *58* | | | 16/16 | 2.0 ± 0.5 | |
| *67* | | | 16/16 | 2.3 ± 0.5 | |
| *71* | | | *No plaques recovered on aTc* | | |
| *72* | | | 16/16 | 2.3 ± 0.5 | 23%±4 (*n=2*) |
| *99* | | | 7/8 | 2.9 ± 0.7 | 16%±1 (*n=2*) |

[1] Primary plaques were isolated in an infectious center assay after electroporation of the recombineering strain (*M. smegmatis*/pJV53) with gDNA and deletion substrate. Mixed plaques were considered positive if a band was detected by PCR with flanking primers corresponding to the wildtype and mutant products. Shown is the ratio of positive plaques out of the total number screened.

[2] Secondary plaques were derived from 1 or 2 mixed primary plaques and screened by PCR with flanking primers. Shown is the ratio of plaques positive for the pure mutant product out of the total number screened.

[3] Plaques were isolated in a plaque assay after electroporation of the recombineering strain (*M. smegmatis*/pJV138) with gDNA and deletion substrate. Plaques were selected on CRISPR-expressing strains on 100 ng/ml aTc and screened by PCR with primers flanking the deletion site. Shown is the ratio of plaques with the mutant product out of the total number screened.
